# Hist2Cell: Deciphering Fine-grained Cellular Architectures from Histology Images

**DOI:** 10.1101/2024.02.17.580852

**Authors:** Weiqin Zhao, Zhuo Liang, Xianjie Huang, Yuanhua Huang, Lequan Yu

**Affiliations:** Department of Statistics and Actuarial Science, The University of Hong Kong, Hong Kong SAR, China; School of Biomedical Sciences, The University of Hong Kong, Hong Kong SAR, China; Center for Translational Stem Cell Biology, Hong Kong Science and Technology Park, Hong Kong SAR, China

**Author notes:** Contributing authors (Weiqin Zhao, Zhuo Liang, Xianjie Huang).

**Keywords:** Cell Type Prediction, Spatial Transcriptomics, Computational Pathology, Graph Learning

## Abstract

Histology images, with low cost, are unleashing great power of predicting cellular phenotypes in tissue, thanks to the emerging spatial transcriptomics serving as annotations. Recent efforts aimed to predict individual gene expression, suffering from low accuracy and high variability, while no methods are tailored to predict fine-grained transcriptional cell types -the most critical phenotype. Here, we present Hist2Cell, a Vision Graph-Transformer framework, to accurately resolve fine-grained transcriptional cell types (up to 40 cell types) directly from histology images and further create cellular maps of diverse tissues at a customizable resolution. Specifically, trained on human lung and breast cancer spatial transcriptome datasets, Hist2Cell accurately predicts the abundance of each cell type across space in new patient samples with Pearson Correlation Coefficient of biological informative cell types over 0.80, and effectively capturing their colocalization directly from histology images. Moreover, without re-training, it robustly generalizes to large-scale histology cancer cohorts from TCGA, highlighting recurrent cell co-localization and supporting precise survival prediction by revealing distinct tissue micro-environments and insightful cell type-patient mortality relationship. Therefore, Hist2Cell enables cost-efficient histology analysis for large-scale studies of spatial biology and precise cancer prognosis in real-world clinical diagnostics.

## 1 Introduction

The cellular architecture of tissues, where distinct cell types are organized in space, underlies cell-cell communication, organ function, and pathology, enabling our broad and deep understanding of diverse cellular behaviors and biology. For example, cell-cell communication plays an essential role in various biological processes and functional regulations [1, 2], such as immune cooperation in a tumor microenvironment, organ development and stem cell niche maintenance, and wound healing. Although previous digital pathology-based methods [3–5] could segment and identify different cells, they can only identify coarse-grained cell type groups (usually **2** to **4** major cell types) and provide limited information for detailed analysis. Fortunately, the emerging spatial transcriptomics (ST) technologies provide key opportunities to map fine-grained transcriptional (for instance, **80** cell types in lung tissue [6]) resident cell types and cell signaling *in situ* in a scalable manner by probing the full transcriptome, for example, Slide-seqV2 [7] and Stereo-seq [8]. Recently, several computational methods have been proposed for analyzing ST data to reveal cell variability in a tissue context. The most critical step is to characterize the fine-grained transcriptional cell type proportion (or abundance) in each ST spot and one of the most popular strategies is to integrate ST with the reference transcriptome signatures of cell types obtained from relevant scRNA-seq profiles. These methods started from general data interaction methods, like Seurat v3 [9] but soon embraced tailored designs for ST by modeling the read counts with negative binomial distributions (stereoscope [10], RCTD [11], and cell2location [12]), or via non-negative matrix factorization (spotlight [13]), or as a mapping from individual cells (e.g., Tangram [14] and TransImp [15]). Of note, among these methods, cell2location uniquely provides an estimate of the absolute cell abundance on top of its proportion; also, its Bayesian treatment enables a higher sensitivity and resolution by accounting for uncertainty in technical sources of variation and borrowing statistical strength across locations; therefore, it was employed for cell type abundance estimation here.

Considering that current sequencing-based ST technologies generally suffer from low spatial resolution and low RNA capture efficiency (and sequencing depth), recent efforts have also been made to take advantage of the paired histology image to enhance ST analysis. Prominent examples include XFuse, which uses a variational autoencoder to generate super-resolution transcriptome by sharing a latent variable between ST and its image [16], while TESLA [17] and iSTAR [18] further accelerate the computational efficiency of this task via graph-based smoothing. Additionally, SpatialScope [19] utilizes histology to segment cell nuclei and achieves a single-cell level transcriptome resolution through a deep generative model (i.e., diffusion model).

However, a key limitation of these existing imputation-like approaches is the requirement of estimating from spatial transcriptome data from the same or consecutive tissue sections of the same patient, which hinders large-scale studies due to the expensive costs. Relatively, histology images are much cheaper and easier to acquire and are routinely used in clinics [20]. Previous studies have shown that histological images are highly correlated with gene expression and cell type identification, and have been used to predict bulk gene expression profiles [21], coarse-grained cell type abundance [3], and genomic abberations [22]. Furthermore, powered by the ST technologies that provide a large amount of high-quality training data, multiple attempts were made to directly predict the expression of a set of critical genes or the full transcriptome from a cell-size path of image [21, 23–26]. However, these methods that aim to predict the expression of a small group of individual genes were generally reported with limited accuracy, probably due to the lack of predictive information for most genes. Therefore, their predictions are unlikely to serve as reliable sources for estimating fine-grained transcriptional cell abundances and how to identify fine-grained transcriptional cell-type spatial variations and infer cell-cell communication directly from histology images is underexplored and remains an open problem. Moreover, existing methods are also incapable of creating more detailed cellular wiring diagrams for higher-resolution analysis when estimating from spatial transcriptome data, as they ignore the information provided by the histology images.

In this paper, we present Hist2Cell, a Graph-Transformer framework to resolve fine-grained transcriptional (up to 80) cell types directly from hematoxylin-and-eosin-stained histology images and create cellular maps of diverse tissues with a **one-stage** pipeline. Hist2Cell was trained on spatial transcriptome data sets for human lung [6] and human breast cancer [27] with spot-level cell abundance estimations from cell2location [12] algorithm. For held-out test patients, Hist2Cell can accurately predict the spatial variations in the localization of **40 fine-grained transcriptional cell types** -at a resolution of around 100µm, serving as important references for biological research. Specifically, it can predict substantial variation in cell localization within regions labeled as tumors or different tissue structures by clinicians, demonstrating that Hist2Cell captures significant tissue heterogeneity. By feeding the predictions to an existing toolbox like SpatialDM [28], Hist2Cell can also reveal accurate cell-type colocalizations, figuring out the prioritization of communicating cells as well as the identification of interaction spots in the spatial context. Hist2Cell is more accurate than deep learning methods that predict cell abundance solely from a single histology image spot or a limited neighboring context, as it leverages both local and global correlations of histology image spots with the proposed Graph-Transformer architecture while maintaining data diversity with a random subgraph sampling strategy.

As independent external tests, Hist2Cell accurately predicts the spatial distribution of **15 fine-grained transcriptional cell types** and their localization in breast cancer data from an external source [21] without any framework modification or fine-tuning. This suggests that Hist2Cell can robustly deal with the unseen samples despite potential batch effects or technique differences of imaging. As a result, Hist2Cell can provide consensus cellular analysis for biologists on the diverse large cohort studies (e.g., TCGA cohorts) that current ST technology cannot be scaled due to high cost. More importantly, Hist2Cell’s fine-grained transcriptional cell abundance predictions could support precise cancer prognosis. In this study, we demonstrate that it outperforms the previous state-of-the-art pathology models in survival risk prediction tasks on three common cancers and presents promising potential to serve as a cheap yet comparable alternative to expensive bulk RNA-seq for cancer prognosis in real clinical settings. Moreover, leveraging integrated gradients, Hist2Cell uncovers the relationships between different cell populations and patient mortality, which could contribute to validating existing cancer research as well as providing new biological insights. In addition, Hist2Cell could provide the above analysis/application under a higher spatial resolution (e.g., 16×, near single-cell-level) than the original spatial transcriptome technique by utilizing low-resolution cellular maps or directly predicting from histology images. In general, Hist2Cell paves the way for reliable investigations of cellular organization and communication in diverse tissues and precise survival prediction for cancer prognosis in a cost-efficient manner, with potential broader applications in diagnostics and personalized medicine.

To avoid confusion with the mentioned previous works, a detailed comparison among Hist2Cell and previous works is provided in Fig 1c and Supplementary Table 1. Briefly, Hist2Cell is the first method that leverages ST as training to infer fine-grained transcriptional (up to 80) cell types as more predictable phenotypes in new tissue/patient samples. All existing methods either predict only 2 to 4 cell types (Group 1: conventional digital pathology), focus on noisy gene expression (Group 2: ST-Net and variants/HE2RNA), or impute the training samples (Group 3: iStar/XFuse).

**Fig. 1:**
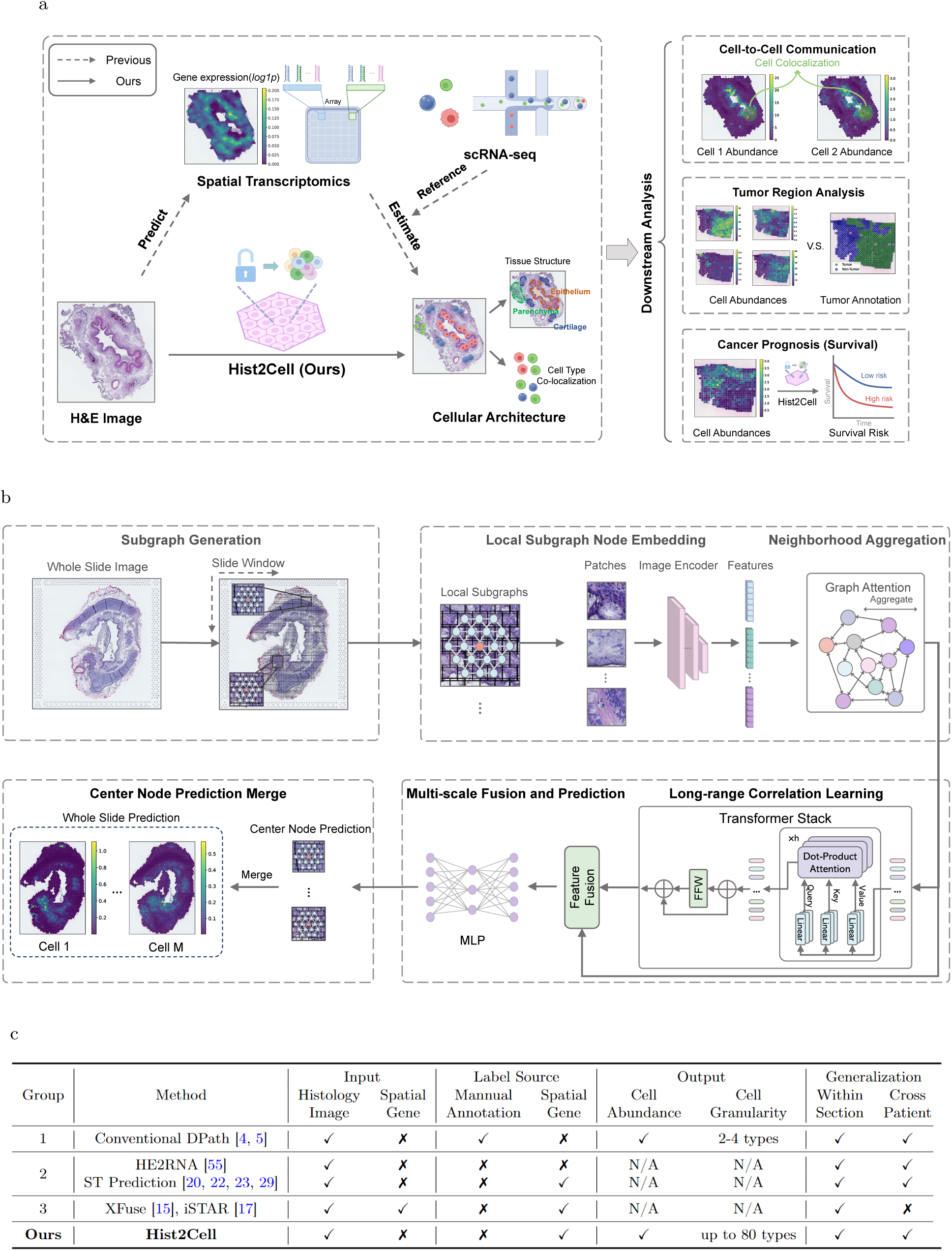
Overview of Hist2Cell. **a,** Left: An illustration demonstrating how Hist2Cell directly predicts spatial cell abundance from histology images (one-stage), in contrast to how previous works estimate spatial cell abundance from spatial transcriptomics data using scRNA-seq data as a reference (two-stage). Right: The three major downstream tasks based on spatial cell abundance predictions by Hist2Cell -encompassing cell-cell colocalization analysis, tumor region analysis, and cancer prognosis (survival risk prediction). **b,** Hist2Cell initially formulates the whole slide image into a spatial graph, followed by sampling local subgraphs as input for the model. It employs a feature encoder, Graph Attention layers, and transformer layers to learn and extract the morphological features of the tissue, as well as to discern both local and long correlations among histology spots. Multi-scale features from these components are fused for fine-grained spot cell abundance prediction. The prediction for the central node of the sampled subgraphs is used as the final output during the inference phase. **c,** Conceptual comparison between our Hist2Cell with conventional digital pathology (DPath), HE2RNA/ST prediction works, and XFuse/iStar works.

## 2 Results

A schematic figure illustrating the workflow of Hist2Cell is presented in Figure 1, along with a summary of example downstream tasks depicted. Initially, Hist2Cell crops the entire slide image into a multitude of spot patches, with spot size 150 × 150*µ*m^2^ according to the ST platforms. Treating spots as nodes and their spatial neighboring relationships as edges, each whole slide image is formulated into an independent graph. Subsequently, during the iteration of constructed graphs, multiple subgraphs are randomly sampled from a certain whole slide image to serve as training inputs for Hist2Cell. Each subgraph consists of a central node and its k-hop spatial neighborhood. Following this, an image encoder extracts node embeddings from the images constituting each subgraph. Hist2Cell then employs a Graph-Transformer architecture to learn and integrate both short-range local and long-range global relationships among a batch of subgraphs from the same whole slide image, thereby enhancing the prediction of cell abundance for each node. The image encoder and the Graph-Transformer architecture are concurrently trained in an end-to-end fashion, minimizing the distance between the cell abundance predictions and ground truth labels estimated from the spatial transcriptome data. In the inference phase, Hist2Cell uses the predictions of the center nodes as the conclusive prediction and implements a sliding-window strategy to ascertain the cell abundance prediction for all spots. To assess the effectiveness of Hist2Cell, we study healthy human lung and human breast cancer tissues and large-scale TCGA histology image datasets.

### 2.1 Hist2Cell Maps Cellular Architectures and Detects Key Colocalized Cell Types in Human Lung Tissues

To validate Hist2Cell, we initially applied the framework to data from 5 proximal-to-distal locations in healthy human lungs from 4 donors [6]. This dataset provides 11 hematoxylin-and-eosin-stained histology slides, resulting in a total of 20,770 spots with 80 fine-grained transcriptional cell-type abundances estimated by the cell2location algorithm according to the authors. More details about the 80 cell types can be found in the Supplementary Fig. 1. All performance comparisons, visualizations, and analyses are reported using leave-one-donor-out cross-validation, where we iteratively trained Hist2Cell on three of the donors and made predictions on the remaining held-out donor.

#### Cell Type Identification

As depicted in Fig.2.c, the cell type identification performance of Hist2Cell was compared with methods designed for predicting local gene expression from histology images including STNet [21], DeepSpaCE [29], Hist2ST [23], and THItoGene [24]. For these ST-prediction-based methods, we follow their default two-stage setting as first predicting spatial gene expressions and then using these predictions to estimate cell abundance. As shown in the figure, Hist2Cell demonstrated more accurate predictions of cell type abundances across locations compared to other methods, as evidenced by the overall Pearson correlations of all folds. Specifically, the analysis revealed that for **79 out of 80** cell types, the predicted cell abundance from Hist2Cell positively correlated with the provided cell abundance labels across all patients, with an average Pearson’s correlation of 0.31 among all 80 cell types and patients, demonstrating a performance increase in PCC of approximately **50%** improvement compared with the best baseline methods. We also examined the top 30% cell types across 11 slides, and the average Pearson’s correlation reached 0.50 vs 0.43 by the best counterpart as shown in the updated Supplementary Fig. 19 and Supplementary Fig. 20. Moreover, despite the divergence of different cross-validated slides, we found out a group of **40 fine-grained transcriptional cell types** can be predicted with high to moderate accuracy (Pearson’s correlation above 0.25 for at least 50% slides.) by Hist2Cell. These results underscore the superiority of our methodology and the one-stage prediction **strategy advantage**, suggesting that Hist2Cell can directly decode fine-grained cellular architectures directly from histology images by modeling the tissue morphology feature, local context interactions, and long-range correlations with the slide. Specifically, we also adopt the proposed one-stage prediction strategy of Hist2Cell and only adopt the network architecture of the baseline methods as the compared counterpart, and the results are shown in Supplementary Fig 3. The adapted baseline methods gained instant performance increase, illustrating the efficacy of avoiding the noise contained in the ST predictions, furthermore, our approach also surpasses "adapted" methods, highlighting the **model advantage** of our graph-transformer architecture and subgraph sampling-based training paradigm. Particularly, we highlight that Hist2Cell showed significantly high positive correlations with several key and informative cell types, including CD4+ effector memory T cells (CD4 EM Effector) with Pearson’s R of **0.68** and Jason-Shannon Divergence of 0.24, CD8 EM with Pearson’s R of **0.79** and Jason-Shannon Divergence of 0.31, ciliated cells with Pearson’s R of **0.62** and Jason-Shannon Divergence of 0.24, and gamma-delta T (gdT) cells with Pearson’s R of **0.68** and Jason-Shannon Divergence of 0.25 (Fig. 2.a). Note that these highly predictable cell types have critical functions in the human airway system, for example, CD8 EM cells are essential in directly eliminating infected or malignant cells in situ [30], and ciliated cells play a pivotal role in airway homeostasis by trapping and expelling microorganisms, mucus and other debris [31]. Systematically, we also found Hist2Cell excelled in predicting the abundance for cell types exhibiting a higher spatial auto-correlation denoted by Moran’s *I* (i.e., spatially variable cell types, as detailed in Fig. 2.d). In other words, on top of the reasonably high performance across all cell types, Hist2Cell achieves even more accurate prediction for cell types that may have specific spatial patterns, which often relate to biological significance [32–34]. Beyond the quantitative assessment, visualization of the predicted cell type abundances in tissue also shows that Hist2Cell accurately identifies spatial patterns of cell type distributions, with fewer false positives compared to other methods, for example in ciliated cells, IgA plasma cells, and SMG Serous cells (Fig. 2.f).

**Fig. 2:**
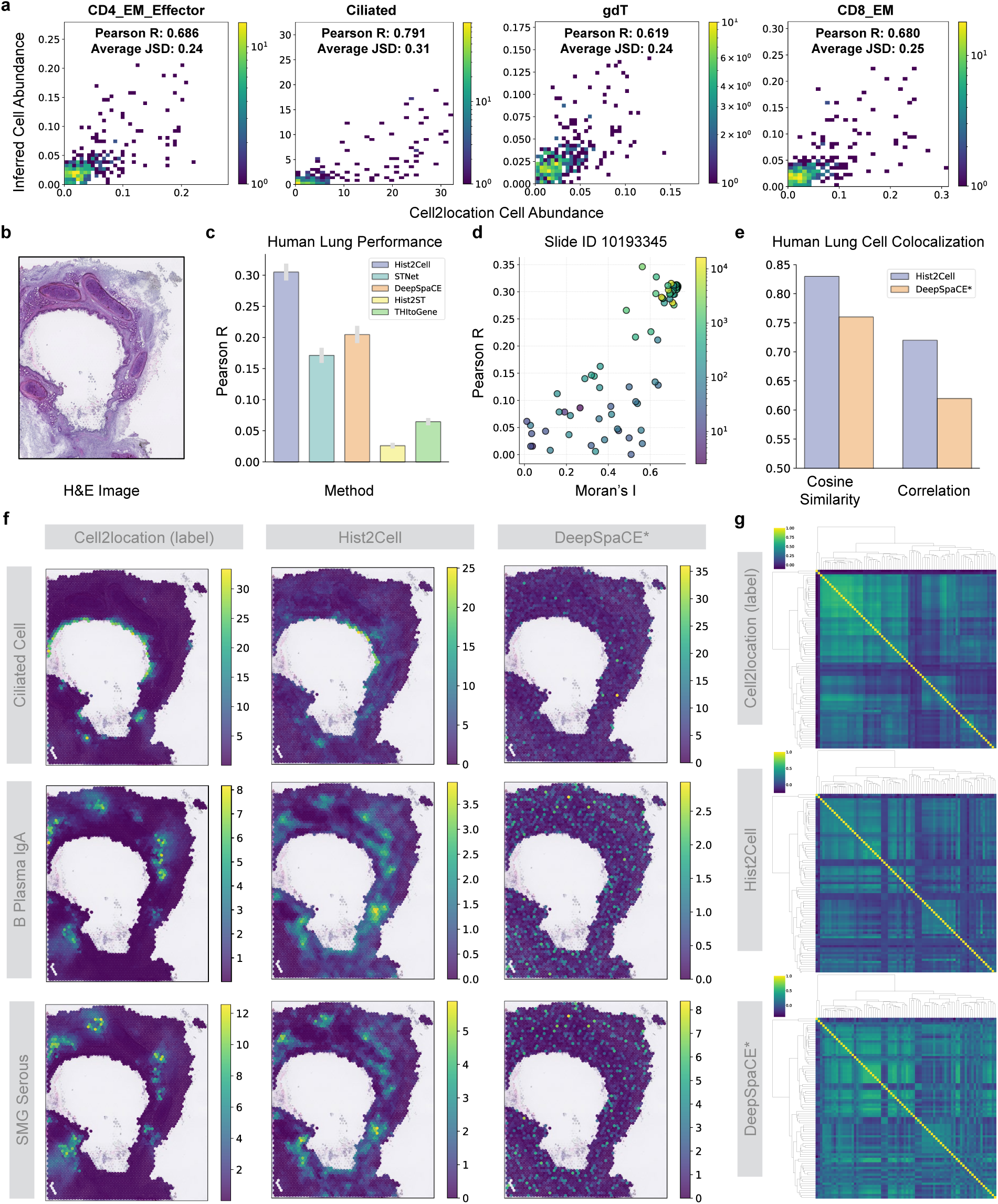
Hist2Cell can map fine-grained human lung cellular architectures. **a,** 2D histogram plots showcasing the concordance of cell abundance between ground truth (x-axis) and Hist2Cell’s prediction (y-axis) across all testing spots in the healthy human lung slide. Color denotes 2D histogram counts. Pear-son’s R denotes Pearson’s correlation coefficient, and JSD denotes Jensen–Shannon divergence. Hist2Cell showed significantly high positive correlations with several key cell types. **b,** Example H&E image for slide (ID) 9258464. **c,** Histogram depicting the average Pearson’s R values for cell abundance prediction in the leave-one-donor-out cross-validation experiment conducted on the healthy human lung dataset. Hist2Cell demonstrated more accurate predictions of fine-grained cell type abundances across locations compared to other ST prediction baselines. **d,** Scatter plot for one test slide illustrating the relationship between spatial auto-correlation (Moran’s *I*; x-axis) and Hist2Cell’s prediction performance (y-axis) in the healthy human lung dataset. Hist2Cell excelled in predicting cell types exhibiting higher spatial auto-correlation. **e,** His-togram depicting the cosine similarity and correlation between Moran’s R for cell type pairs calculated from different models’ predictions and ground truth cell abundances. **f,** Related to (b), the visualizations comparing the spatial cell abundances as determined by ground truth, Hist2Cell, and DeepSpaCE* predic-tions for select key cell types. Hist2Cell show less false positives than DeepSpaCE*. **g,** Related to (e), the clustermaps colored by the bivariate statistic Moran’s R for cell type pairs. Top: results from ground truth cell type abundances estimated from the spatial transcriptomics data using cell2location algorithm. Middle: results from predictions of Hist2CellBottom: results from predictions of DeepSpaCE* (the best performed adapted ST prediction baseline). Hist2Cell provided a more consistent cell-cell colocalization analysis with the ground truth cell abundances.

#### Cell Type Colocalization Patterns

Next, we examined the pair-wise colocalization among these 80 cell types by performing a spatial correlation with bi-variant Moran’s R as implemented in SpatialDM [28]. Briefly, this statistic calculates the correlation between the abundances of two cell types while considering spatial auto-correlation. Overall, we found that the patterns of colocalization among these 80 cell types are highly consistent between the quantified abundance from ST and predicted ones from histology by our Hist2Cell (Fig 2.g). By contrast, discordance is evident if using the cell type abundance predicted by the counterpart method DeepSpaCE* (the best performed adapted baseline method in Supplementary Fig 3, * represents adapted baselines using one-stage prediction strategy). For a direct numerical comparison, we also visualized the cosine similarity and correlation between the colocalization calculated from the predictions of different models and the ground truth cell abundances, and Hist2Cell excelled in both metrics as shown in Fig 2.e. As a prominent example, we noticed that Hist2Cell identifies colocalization between IgA plasma cells and submucosal glands (SMGs) in human airways, which is one of the major discoveries in the original study [6]. Impressively, with the observed cell type abundance from ST, IgA plasma was only ranked as the 1st colocalized cell type for SMGs, and IgA plasma was ranked the 3rd for SMGs’ colocalized partner if using predicted cell type abundance by our method (Fig 2.f). These results suggest that the global colocalization patterns and the individual cell type pairs with critical interaction can be identified by our method, by only using histology images. Detailed cluster maps can be found in Supplementary Fig. 13 to Fig. 15.

#### Expert Manual Annotation Evaluation

By comparing Hist2Cell predictions with manual annotations from the original study [6] of the dataset (Fig 3.a), we conclude that Hist2Cell’s outputs can serve as reliable references for biological research, such as the discovery of macro and micro-anatomical tissue compartments. Specifically, Hist2Cell’s accurately predicted cell types correspond to their known locations, like ciliated epithelial cells in the lumen of the airway surrounded by basal cells, and alveolar types 1 and 2 (AT1 and AT2) cells in lung parenchyma, as shown in Fig 3.b. Furthermore, Hist2Cell accurately localized one fibroblast population, enriched in the airways, to its specific region around the airway epithelium (peribronchial fibroblasts-PB-fibro), as depicted in Fig 3.c. This finding also validates the clinical value of Hist2Cell, as PB-fibro is recognized as a key cell type in lung disease [6]. In Supplementary Fig. 11, we also compare the predictions of DeepSpaCE* with the manual annotation, which shows poor alignments between the cell types and the tissue compartments.

**Fig. 3:**
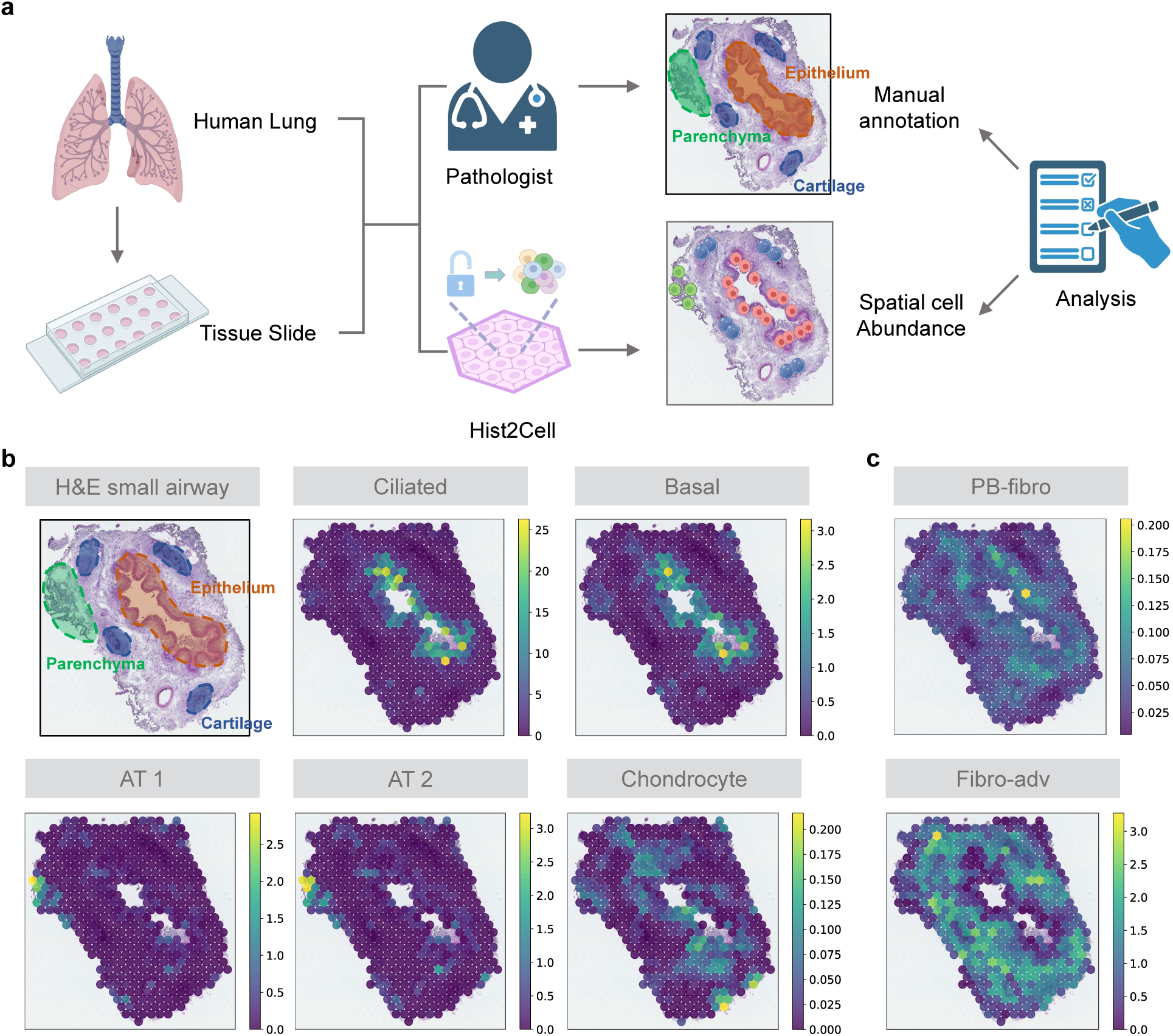
**a,** Analysis workflow incoporating Hist2Cell and manual annotation. **b,** Mapping with Hist2Cell on Visium ST of a bronchi section demonstrates the alignment of cell types with their expected anatomical structures. H&E staining and cell abundance predictions by Hist2Cell (denoted as density scores) for ciliated, basal epithelium, AT1, AT2, and chondrocyte cell types are overlaid on a histology image. Dotted lines demarcate the epithelium (pink), parenchyma (green), and cartilage (brown). **c,** Cell abundance predictions by Hist2Cell indicate that PB-fibro cells localize around the airway epithelium.

### 2.2 Hist2Cell can Generalize to External Unseen Breast Cancer Samples

To evaluate the generalizability and clinical applications of Hist2Cell, we conducted an external validation experiment using human breast cancer tissues. Specifically, the Hist2Cell model was trained on the her2st breast cancer dataset (13,620 spots in 36 sections from 8 patients) [27] to predict cell abundance for 39 fine-grained cell types and was then directly applied to the STNet breast cancer dataset (30,655 spots in 69 sections from 23 patients) [21] without any re-training. All performance comparisons, visualizations, and analyses are reported using the external STNet dataset. More details about the 39 cell types can be found in the Supplementary Fig. 2.

#### Cross-lab Generalization in Cell Prediction

The overall Pearson correlations on the external dataset revealed that Hist2Cell provides more accurate predictions of cell type abundances across various locations compared to other methods and their variants (see Fig.4.c and Supplementary Fig 4). Specifically, on the external dataset, Hist2Cell’s predictions positively correlated with all 39 cell types across all samples, exhibiting an average Pearson’s correlation of 0.29. On the contrary, STNet (the best-performed baseline method in Fig.4.c) only has an average Pearson’s correlation of 0.19 on the external dataset. With the top 30% cell types across all slides, the average Pearson’s correlation reached 0.54 vs 0.44 by the best counterpart as shown in the updated Supplementary Fig. 21 and Supplementary Fig. 22. Moreover, despite the domain shift of different labs, we found out a group of **15 fine-grained transcriptional cell types** can be predicted with high to moderate accuracy (Pearson’s correlation above 0.25 for at least 50% slides.) by Hist2Cell. This suggests that Hist2Cell possesses superior generalization capabilities on external data despite technical differences in sample preparation and imaging techniques of different datasets. The favorable generalization of Hist2Cell might be attributed to its one-stage prediction strategy, model architecture, and the random subgraph sampling strategy. The one-stage prediction strategy avoided learning the compounded errors and shortcuts in the spatial gene expression and directly learned the relationship between the histological morphology features related to the fine-grained cell types to enhance the generalization of the model. The model architecture integrates both local and global correlations within tissue slides, facilitating the learning of more robust histological image representations. Additionally, the random subgraph sampling strategy preserves data diversity throughout the training process. Moreover, we need to highlight that, Hist2Cell maintains significantly high positive correlations with several key cell types on the external dataset (Fig. 4.a), such as fibroblast as a common component in the tumor microenvironment of breast cancer [35] with Pearson’s R of **0.71** and Jason-Shannon Divergence of 0.16, luminal epithelial cells of the mammary gland that can produce basal cells upon oncogenic stress [36] with Pearson’s R of **0.87** and Jason-Shannon Divergence of 0.08, lgA plasma cells as the major player of the tumor-immune interaction [37] with Pearson’s R of **0.69** and Jason-Shannon Divergence of 0.22, and vascular-associated smooth muscle cells with Pearson’s R of **0.74** and Jason-Shannon Divergence of 0.17. Hist2Cell also achieves elevated accuracy in predicting spatially variable cell types (i.e., higher Moran’s *I*; Supplementary Fig. 8) on the external dataset, corroborating findings from the human lung dataset. Beyond quantitative assessment, the ability of Hist2Cell and other methods to accurately identify locations with conspicuously high cell abundance of specific cell types was also evaluated on the external dataset. Visualizations in Fig. 4.e demonstrate that Hist2Cell accurately identifies the presence of cell types such as IgA plasma cells, CD4-positive helper T cells, and luminal epithelial cells of the mammary gland, with fewer false positives than other methods on the external dataset. When comparing these visualizations with manual pathology tumor/normal annotations (Fig. 4.b), it was observed that the inferred cellular architectures closely match human expert annotations, showing the capability of capturing intratumor heterogeneity of Hist2Cell.

**Fig. 4:**
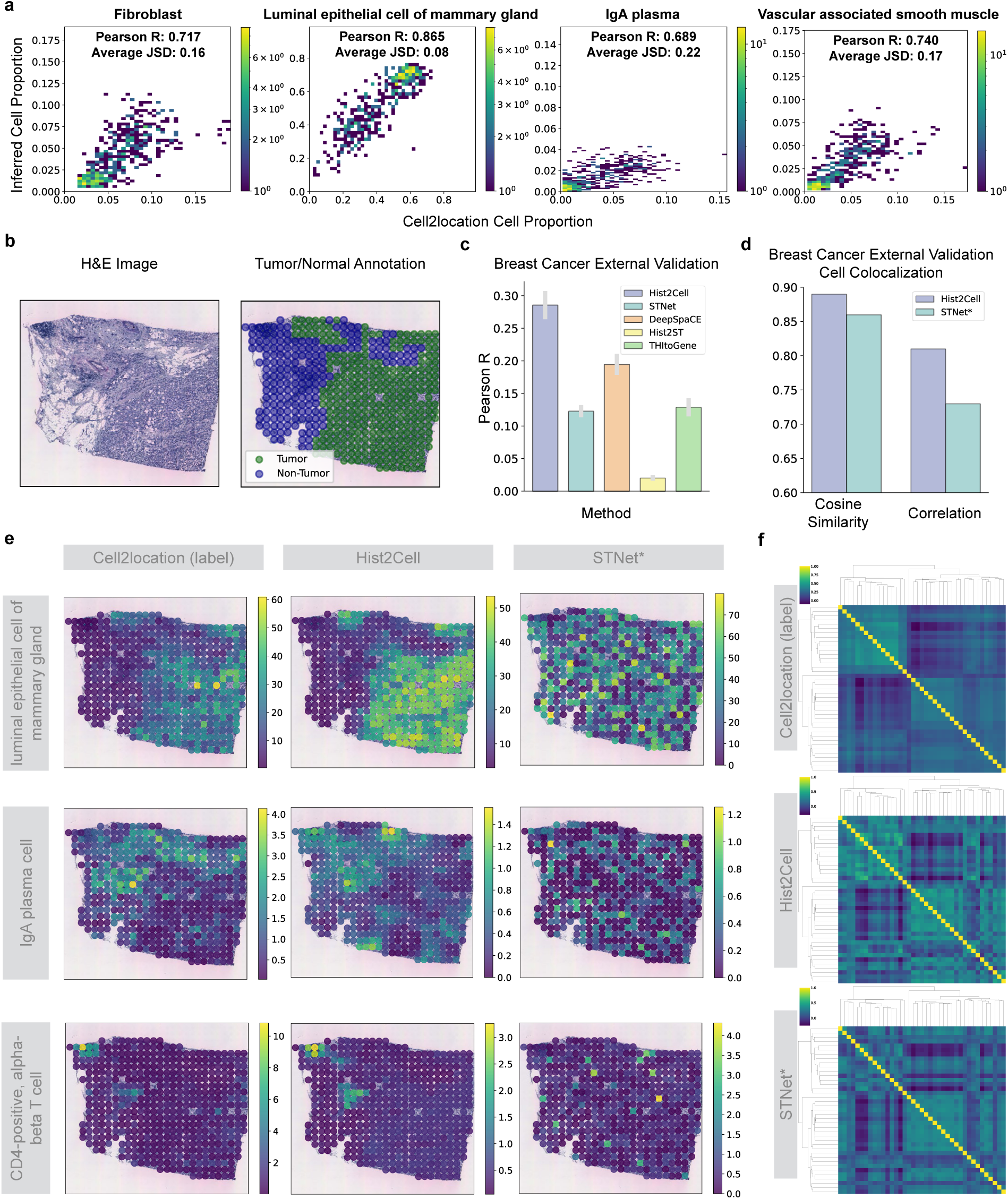
Hist2Cell can generalize to external unseen breast cancer samples. **a,** 2D histogram plots showcasing the agreement between the ground truth cell proportion (x-axis) and that predicted by Hist2Cell (y-axis) across all testing spots in the external breast cancer slide. **b,e** H&E images, manual tumor/normal annotations of slide (ID) 23508D2, and visualizations comparing the spatial cell abundances as determined by ground truth, Hist2Cell and STNet* for certain key cell types. Hist2Cell showed fewer false positives compared to STNet*. **c,** A histogram representing the average Pearson’s R values for cell abundance prediction in the external breast cancer dataset. Hist2Cell demonstrated superior accuracy and generalizability in predicting fine-grained cell type abundances across different locations compared to other ST prediction baselines. **d,** Histogram depicting the cosine similarity and correlation between Moran’s R for cell type pairs calculated from different models’ predictions and ground truth cell abundances. **f,** Related to d, clustermaps colored by the bivariate statistic Moran’s R for various cell type pairs. The top shows results from ground truth cell type abundances estimated from spatial transcriptomics data using the cell2location algorithm. The middle displays results from predictions by Hist2Cell. The bottom presents results from predictions by STNet*(the best-performed adapted ST prediction baseline). Hist2Cell provided more consistent cell colocalization patterns compared to the ground truth cell abundances.

#### Recognize Cell Co-localization on External Data

Furthermore, we assess if Hist2Cell can effectively identify colocalization among these fine-grained cell types on the external dataset. Similar to the human lung data, we used the bi-variate Moran’s R statistic to calculate the spatial correlation and analysis of the cluster maps on this value. Fig.4.d and Fig.4.f indicate that Hist2Cell outperforms the other method, showing a global pattern more akin to that obtained from the probed ST by cell2location and higher cosine similarities as well as correlation. As a notable example, the colocalization between effector memory CD8-positive, alpha-beta T cell with myeloid dendritic cell is a highly ranked interacting pair (Moran’s R=0.38). By directly predicting from histology, Hist2Cell also identifies the high spatial correlation between these two cell types (Moran’s R=0.26), confirming the complex cooperation among immune cells against tumor cells [38]. Specifically, with the observed cell type abundance from ST, mature NK T cell was ranked as the 2nd colocalized cell type for natural killer cell, and it is ranked as the 1st colocalized cell type if using predicted cell type abundance by our method (Fig. 4.e). Detailed cluster maps can be found in Supplementary Fig. 16 to Fig. 18. Taken together these results suggest that the predictions from Hist2Cell uncover promising colocalization patterns despite batch effect and technique difference, potentially facilitating the discovery of context-specific cellular cooperation and signaling on external datasets, which is crucial for its research and clinical usage.

#### Pan-tissue Analysis Potential

For a "boarder sense" of generalizability, an interesting question is whether Hist2Cell can support training in one type of cancer and prediction in another type. Actually, this setting can be referred to as pan-cancer or pan-tissue cell typing which requires both establishing systematic cell type nomenclature across datasets and training in multiple tissues/cancers. Two experiments have been conducted to illustrate that Hist2Cell has the potential to transfer the knowledge learned on one organ to another (from breast cancer tissue to human lung tissue in our study), laying the groundwork for future research that could explore pan-tissue generalization. First, we demonstrated that transferring model parameters pre-trained on breast cancer tissue to lung tissue improves performance under low-data regimes, achieving an 11% increase in average Pearson R over training from scratch, as shown in Supplementary Fig 7. Second, by merging some fine-grained cell types in both datasets into coarse-grained categories like Epithelial Cells, we applied our model trained on breast cancer directly to lung tissue. This approach yielded a PCC of 0.26 for Epithelial cells, suggesting the feasibility of cross-tissue generalization with well-aligned cell type annotations.

### 2.3 Hist2Cell Enables Consensus Cellular Analysis from Large-scale H&E Cohorts

As Hist2Cell generalizes well on external breast cancer samples, we incorporate large-scale breast cancer histology slide datasets from the TCGA repository and adopt Hist2Cell to provide consensus cellular anal-ysis, as shown in Fig.5.a. It is worth noting that there were several notable differences between the TCGA and training breast cancer data. The TCGA slides are collected and scanned at different institutions and only have bulk RNA-seq data that is not conducted on the same tissues used for imaging, which poses a substantial challenge for fine-grained cellular analysis. We applied Hist2Cell directly to H&E slides of 565 TCGA breast tumor samples from 565 patients without any model re-training.

**Fig. 5:**
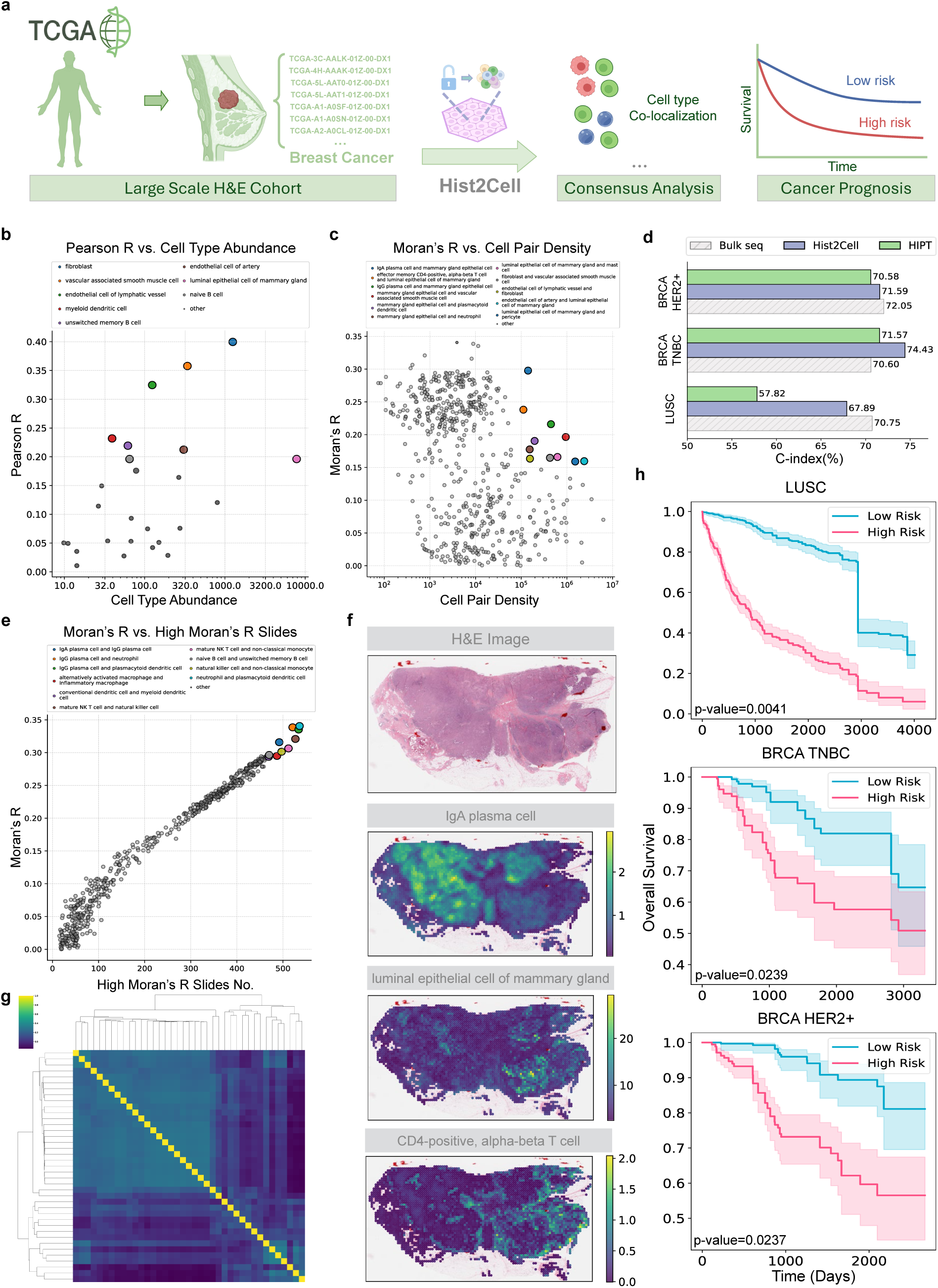
Hist2Cell can provide consensus analysis and precise cancer prognosis on large-scale H&E TCGA cohorts. **a,** An illustration of the large-scale study process using Hist2Cell on TCGA breast cancer cohorts, demonstrating how precise consensus cellular analysis can be achieved and how cancer prognosis (survival risk prediction) can be done on slides spanning hundreds of patients. **b,** A scatter plot illustrating the relationship between cell abundance (x-axis) and Hist2Cell’s prediction performance (y-axis) across all cases in the TCGA breast cancer cohort. Hist2Cell excelled in predicting cell types with higher abundances. **c,** A scatter plot showing the relationship between cell pair density (x-axis) and the average bivariate statistic Moran’s R as calculated by Hist2Cell (y-axis) among all cases in the TCGA breast cancer cohort. Hist2Cell effectively distinguished cell pair colocalization when the density of the cell pairs was high. **e,** A scatter plot illustrating the relationship between the number of slides with high Moran’s R values (x-axis, Moran’s R greater than 0.20) and the average bivariate statistic Moran’s R as calculated by Hist2Cell (y-axis) for all cases in the TCGA breast cancer cohort. Highlighted points represent colocalization patterns Hist2Cell identified among most slides of the entire cohort. **g,** Consensus clustermaps colored by the average bivariate statistic Moran’s R, calculated from Hist2Cell’s predictions for cell type pairs across all cases in the TCGA breast cancer cohort. **f,** H&E images, and visualizations of spatial cell abundance predictions by Hist2Cell for a slide in the TCGA breast cancer cohort. Hist2Cell’s inferred cell abundances effectively differentiated between normal and tumor regions in breast cancer tissue. **d,** Histogram depicting the average C-indices of Cox regression models predicting survival of three different cancer subtypes: LUSC, BRCA-TNBC, and BRCA-HER2+ in a 10-fold cross-validation experiment. Hist2Cell demonstrates superior performance than the previous state-of-the-art model and serves as a promising alternative to bulk RNA-seq. **h,** Cross-validated KM curves for patients split into high-and low-risk groups by the predicted risk scores of Cox regression models using predicted cell abundances from Hist2Cell. The survival risk of the low-risk group is significantly lower than that of the high-risk group (p-value < 0.05, T-test).

#### Pseudo-bulk Cell Abundance Evaluation

To quantitatively evaluate the predictions of Hist2Cell, follow STNet [21], we first estimate the bulk cell abundance of each sample according to their bulk RNA-seq with cell2location. The predictions of Hist2Cell were averaged into a pseudo-bulk cell abundance profile for each sample. For the cell type abundance matrix of 39 cell types by 565 TCGA slides, we calculated Pearson’s correlation between the predictions and the pseudo-bulk cell abundance profile in row wide, i.e., for each cell type across 565 slides, resulting in 39 scores. For all samples in the large cohort, the predicted pseudo-bulk cell abundance profile correlated positively with 26 of 39 cell types. The correlation was significantly positive for 13 cell types (FDR lower than 0.05). Although the Pearson R values are lower than those in the previous section, we assume this is due to the since bulk RNA-seq data are not generated from the same tissue as Hist2Cell’s input slide. However, we believe these positive Pearson R values likely reflect Hist2Cell’s ability to extract meaningful molecular information. Indeed, for a negative control, we performed shuffling of labels between slides (shuffling columns of the label matrix), and we found this null model gives Pearson’s R all near zeros (with FDR consistently failing to pass the 0.05 threshold), proving that the performance of Hist2Cell is non-trivial. Among the predicted fine-grained cell types, fibroblast cells have the highest positive correlation as shown in Supplementary Fig. 31, with an average Jason-Shannon Divergence of 0.26 among all 565 TCGA breast cancer slides. We also noticed that Hist2Cell excelled in predicting cell types with higher bulk cell abundance (Fig.5.b), which were generally more significant in tumor analysis. For instance, in Fig.5.b, Hist2Cell shows better predictions for cell types including fibroblast, luminal epithelial cell of the mammary gland, and naive B cell, which is consistent with the findings in the previous section.

#### Integrated Analysis from Large Cohort

More importantly, by aggregating the cellular architectures from hundreds of patients, Hist2Cell unleashes the remarkable potential of large-scale public H&E cohort and is capable of providing consensus and robust analysis for cell-cell communication analysis for biological research. As shown in Fig.5.c, Hist2Cell identified cell pairs with both high average Moran’s R statistic and high cell pair density (product of bulk cell abundances) among all 565 breast cancer samples. We further investigated recurrence patterns, as depicted in Fig.5.e, by visualizing cell type colocalization relationships characterized by both high average Moran’s R values and a significant number of slides with high Moran’s R (greater than 0.20). Specifically, we highlighted those relationships where the number of slides with high Moran’s R exceeded 470. Also, we show the detailed cell type colocalization analysis provided by Hist2Cell in Fig.5.g (details in Supplementary Figure 30). Among them, the colocalization of T cells with myeloid cells can still be observed in the consensus setting (Moran’s R with T cell ranked 3rd for myeloid cell), which is consistent with the findings in previous study [27]. In addition, Hist2Cell inferred cell abundances had the power to distinguish normal/tumor regions of breast cancer tissues from TCGA as shown in Fig.5.f. Moreover, we explore the impact of large-scale studies on the analytical power of Hist2Cell. As depicted in Supplementary Fig 12, we visualized the cell type colocalization analysis performed by Hist2Cell across various sample sizes. Specifically, *n* denotes the number of randomly sampled slides from the TCGA breast cancer cohort. Using *n* values of 1, 2, 4*, . . . ,* 128, and the entire cohort, we generated consensus clustermaps based on Hist2Cell’s predictions. It was observed that the patterns of cell type colocalization gradually stabilized as the number of slides increased, indicating that more reliable analysis can be achieved with larger sample sizes. These analyses suggest that Hist2Cell can generalize to large-scale public H&E cohorts despite the technical differences in the samples and imaging techniques, and thus provide consensus cellular analysis for biologists on the diverse large cohort studies that current ST technology cannot be scaled due to high cost.

### 2.4 Hist2Cell Provides Precise Cancer Prognosis on Clinical Slides

Apart from analyzing large-scale H&E cohorts for consensus biological analysis, we validate that Hist2Cell is also capable of facilitating real-world applications like cancer prognosis among clinical slides (Fig. 5.a). As previous studies have uncovered the benefit of introducing molecular information into survival risk prediction [39], we envision Hist2Cell’s effectiveness in this significant real-world application. Specifically, we evaluate Hist2Cell’s performance on survival risk prediction tasks for three important cancer subtypes including Lung Squamous Cell Carcinoma (LUSC), Triple-Negative Breast Cancer (BRCA-TNBA), HER2-positive Breast Cancer (BRCA-HER2+), with the clinical slides, bulk RNA-seq and survival information obtained from TCGA. For the evaluation metric, we refer to the concordance index (C-index) to measure the predictive performance of correctly ranking the predicted patient risk scores with respect to overall survival.

#### Supporting Precise Cancer Survival Analysis

To enhance the capability of Hist2Cell on cancer tissues, we first refer to the largest ST dataset for Lung Squamous Cell Carcinoma (LUSC) and Breast Cancer (BRCA) from HEST-1k [40] study. Then, we applied the Cell2location [12] algorithm to these cancer ST data with scRNA-seq references from the Multi-omics Spatial Atlas of the Human Lung^1^ and the Human Breast Cell Atlas^2^ to obtain transcriptional fine-grained cell type abundances that we used as supervision for Hist2Cell on these three cancer subtypes. Note that for each cancer, we train independent Hist2Cell models. We then utilized the trained Hist2Cell model to infer the transcriptional fine-grained cell abundances from TCGA slides. To validate Hist2Cell’s usefulness, we compared it to HIPT [41], which has shown state-of-the-art survival prediction performance. Following the classic multi-instance learning paradigm for WSI analysis, we used HIPT to extract patch-level morphological features from TCGA slides. Then, the Hist2Cell-predicted cell abundance predicted and the HIPT-extracted features were aggregated as slide-level cell abundance profile and slide-level morphological feature respectively. Slide-level Cox regression models are trained with the slide-level cell abundance profile and slide-level morphological feature as input. Results from 10-fold patient-level cross-validation are presented in Fig. 5.d. We observed that Hist2Cell outperformed HIPT among all three cancers, especially for BRCA HER2+ patients, achieving about **10% higher C-index**. Notably, we emphasize that Hist2Cell trained on just **114** breast cancer slides and **34** lung cancer slides, far fewer than HIPT (**10,000** WSIs from TCGA). This finding demonstrates the utilization of additional ST and scRNA-seq supervisions during Hist2Cell’s training could benefit precise cancer prognosis, as the predicted cell abundances reflect distinct tissue micro-environments that aid in analyzing organism function or disease processes [6, 42]. Additionally, we compared with UNI [43], a pathology foundation model pre-trained on over **100,000** diagnostic WSIs from 20 major tissue types. Though performing better than HIPT, UNI still showed lower average performance than Hist2Cell (Supplementary Table 2), with at most **3.8%** performance drop in BRCA-HER2+.

#### Cost-efficient Bulk RNA-seq Alternative

Furthermore, as the expensive bulk RNA-seq collected from the patient’s blood sample serves as an important reference in real-world cancer prognosis, we investigate if the predictions of Hist2Cell have the potential to serve as a comparable but cost-efficient alternative. In Fig. 5.d, we benchmarked Cox regression models trained on bulk RNA-seq data of the patients, finding that Hist2Cell achieved comparable performance and even outperformed the bulk RNA-seq baseline by **4%** C-index on BRCA-TNBC. Interestingly, for LUSC, where molecular information probably plays a significant role, both Hist2Cell and the RNA-seq model showed obvious advantages over HIPT. These results motivate us that Hist2Cell indeed has the potential to serve as a powerful yet cheaper alternative to bulk RNA-seq in cancer prognosis.

#### Helping Patient Risk Stratification

We also visualized the KM curve for the three cancer subtypes across the test patients, separated by the predicted risk score of the Cox regression model trained on Hist2Cell’s predicted cell abundances (Fig. 5.h). We observed a significant difference (p-value<0.05, T-test) in survival risk scores between the low-risk group and the high-risk group, and the survival curves stratified by the risk score of the Hist2Cell-based Cox regression model also show a clear separation between the low-risk and high-risk groups, suggesting that Hist2Cell capture important molecular features relate to tissue micro-environments predictive of patient mortality and could help patient risk stratification in hospital.

#### Interpretation of Predictions and Biological Insights

We can leverage well-established deep learning interpretation methods to understand what the Hist2Cell-prediction-based model pays attention to in the fine-grained transcriptional cell populations when making predictions during the survival analysis. In Supplementary Figure 32 and Figure 33, we utilized the integrated-gradients method [44] to identify the cell types contributing to the prediction of patients’ death risk in different time intervals. Specifically, we calculated the integrated gradients on each cross-validated test slide to obtain an average integrated gradient for each fine-grained transcriptional cell type. We considered these feature attributions to provide a more interpretable relationship between different cell types and the short-time/long-time survival of cancer patients. Taking two breast cancers (Supplementary Figure 32) as an example, we found out that: (1) in the 10 major cell types identified by the original study of the scRNA-seq reference [45], luminal hormone-responsive (LumHR-), luminal secretory (LumSec-), fibroblasts (fibro-) and immune (CD8-activated) cells play important roles (on the top of the ranked heatmap) in predicting patients’ mortality. On the other hand, other cell types, like the vascular (vas-venous) and the rare cells, showed less related (on the lower part of the ranked heatmap) to the patient’s survival status, which aligns with the existing studies analyzing breast cancer survival [46–51]; (2) the effect of a certain cell type might vary between short-time and long-time survival analysis, for instance, we note that CD8-activated cells will have a stronger effect for long-time survival (4 times bigger average integrated gradients for later survival intervals) for HER2+ cancers. Such observation provides potential biological insights for follow-up hypotheses and validation in future cancer research.

### 2.5 Hist2Cell can Provide Super-resolved Fine-grained Cellular Maps

Hist2Cell, leveraging its scalability enabled by subgraph sampling, excels in inferring super-resolved cell abundance maps. Note that, in our study, to produce higher-resolution cellular map is equivalent to predict the relative cell abundances for more positions on the slide. Under this objective, our approach is able to produce high-resolution cellular maps as we could predict the relative cell abundances for a certain position based on its morphological features from the 224×224 image patches centered on this position and its neighboring image patches. As illustrated in Fig.6.a, by sampling subgraphs from higher-resolution spot coordinates, Hist2Cell is capable of providing super-resolved cellular maps by either imputing (fine-tuning) on lower-resolution cellular maps or directly predicting from histology images with its pre-trained parameters. To assess Hist2Cell’s proficiency in resolving higher resolution cell abundance maps, we inferred 2× cell abundance for a sample in the human lung dataset, as shown in Fig.6.b. It was observed that both imputation-based and prediction-based super-resolved cellular maps provide a more detailed mapping in relation to different tissue structures in concordance with the manual annotations in Fig.3.a. The high-resolution cellular maps validate the findings in the previous study that ciliated epithelial cells are in the lumen of the airway surrounded by basal cells, and alveolar types 1 and 2 (AT1 and AT2) cells are in lung parenchym [6]. In addition, we also produced 4× high-resolution cellular maps in Supplementary Fig 10, and we noticed that as the resolution goes higher, we could have more detailed cellular localized patterns. Moreover, we show more detailed 8× and 16× cellular maps of AT1, B naive and Ciliated cells in Supplementary Fig. 23 to Supplementary Fig. 28, which further substantiates the Hist2Cell’s capability of depicting the super-resolved intensity of the cell population distribution within the tissue. We also provide a zoom-in inspection of the 16× cellular maps of Ciliated in Supplementary Fig. 29, in which we could observe that the grid distance between neighboring spots is close and even smaller than a single cell in the slide and this resolution is almost single-cell level. In Supplementary Fig. 9, we further show the super-resolved cellular maps of human breast cancer tissue, which also demonstrate a more fine-grained boundary consistent with the tumor/normal annotations in Fig 4.c. We observe that imputation-based maps are more consistent with the lower-resolution cellular maps, while prediction-based maps exhibit more concentrated cellular clustering patterns based on the morphological features of the histology images. Although some prior studies have focused on super-resolving low-resolution Spatial Transcriptomics (ST) data [16, 18], we highlight that Hist2Cell is uniquely capable of directly providing super-resolved cellular maps from histology images without the need of ST data from the same or consecutive tissue section of the specific patient. Specifically, Hist2Cell can offer super-resolved cellular maps even when low-resolution spatial cell abundances are unavailable, by utilizing the knowledge embedded in its pre-trained parameters. This unique capability renders super-resolved cellular mapping applicable to the vast array of H&E images routinely utilized in clinical settings. In summary, Hist2Cell provides a cost-effective method for achieving cell abundance super-resolution, enabling the acquisition of high-resolution cellular maps directly from histology images.

**Fig. 6:**
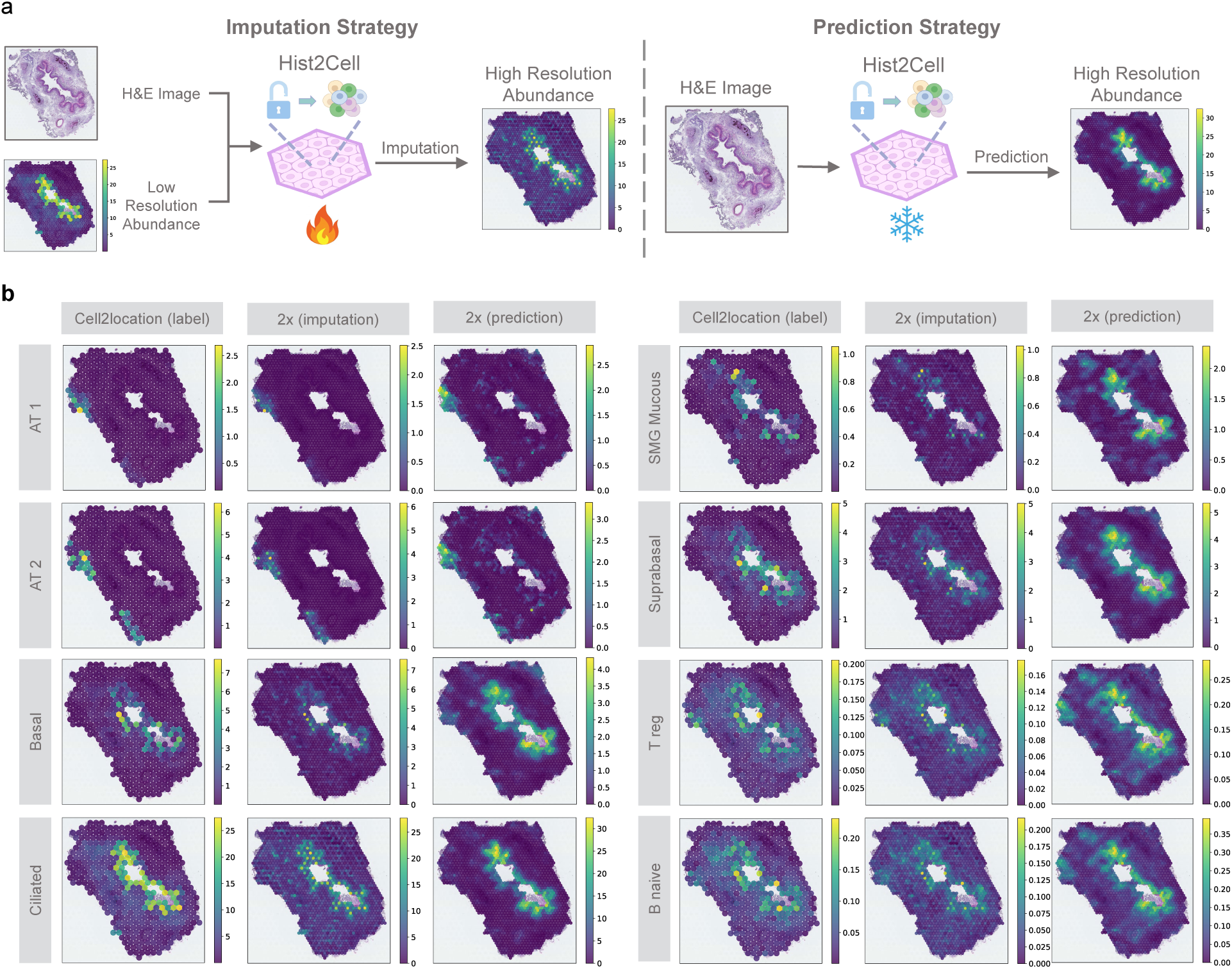
Hist2Cell can provide super-resolved fine-grained cellular maps in a neighbor-aware manner. **a,** An illustration of the two super-resolution strategies based on Hist2Cell. Left: Hist2Cell can impute from low-resolution cellular maps by fine-tuning its pre-trained parameters. Right: Hist2Cellcan also directly predict high-resolution cellular maps with its pre-trained parameters frozen. **b,** Visualizations comparing low-resolution ground truth cell abundance with both imputed and directly predicted high-resolution results by Hist2Cell. Both approaches provide a more detailed mapping in concordance with the manual annotation shown in Fig. 3.a, highlighting different tissue structures.

## 3 Discussion

In this work, we have presented Hist2Cell, a Graph-Transformer framework that accurately decodes fine-grained transcriptional cellular architectures (up to 40 cell types) from tissue morphological features with hematoxylin-and-eosin-stained histology image inputs. The capability of Hist2Cell is beyond previous digital pathology-based methods [3–5], as they can only identify coarse-grained cell type groups (usually 2 to 4 major cell types) and provide limited information for detailed analysis. We demonstrate that the knowledge within spatial transcriptomics and single-cell sequence signatures can be well captured by our deep learning method, and thus transferred to readily available H&E histology images to provide fine-grained transcriptional cellular maps. Hist2Cell produces accurate predictions for held-out donors in human lung tissues. Moreover, it can generalize to external datasets despite batch effect and technique differences of imaging without any re-training, as evident by the breast cancer datasets. We also find that Hist2Cell can capture cell abundance heterogeneity within tumors and different tissue structures. Although this paper focuses on human health lungs and three important cancers, the technical framework of Hist2Cell can be broadly applied to other tissue types.

In addition to predicting and identifying fine-grained cell type abundance, the prediction of Hist2Cell can also be used for analyzing cell colocalization patterns. For example, it reveals the cell type colocalizations in the airway cartilage of human lungs. We also find it promising that Hist2Cell can make predictions of public large-scale H&E cohorts without any model re-training, despite the substantial differences in the data. This opens the door to apply Hist2Cell to large existing cohorts of histology images and provide consensus biology research analysis, such as inferring important bio-markers like cell-cell communication within important tissue sections such as tumor areas. More importantly, we highlight that Hist2Cell could be applied for precise cancer prognosis, such as survival risk prediction. By revealing distinct tissue micro-environments, Hist2Cell has the potential to serve as a comparable yet cost-efficient alternative to bulk RNA-seq in clinical usage, and could provide valuable patient survival risk stratification in real-world applications. Moreover, Hist2Cell could reveal the insightful relationship between different cell populations and patient mortality, helping in validating existing cancer research as well as providing new biological insights. In addition, due to the scalable nature of the designed framework, Hist2Cell can also infer higher resolution cell abundance maps than the current spatial transcriptome technique.

For the first time, Hist2Cell proved that predicting transcriptional fine-grained cell types of biology interest from histology images is feasible and realistically more accurate than individual genes. Although other machine-learning approaches can identify coarse-grained cell types (e.g., 2-4 cell types) from whole histology slides [4, 5], Hist2Cell can decode much more complex cell architectures in a fine-grained manner (up to 80 different cell types) by learning from spatial transcriptomics as well as single-cell sequence signatures. With multiple datasets, we vividly demonstrated that direct cell type prediction provides rich biological insights: our proof-of-concept study finds that the combination of spatial transcriptomic, single-cell sequence, and deep learning can provide direct fine-grained cellular architecture identification solely from histology images, allowing further study of complex cell interactions and variations (e.g., in the human lung dataset) as well as facilitating cancer prognosis by revealing patients’ molecular attributes. Nevertheless, we also recommended image-based cell classification algorithms like HoverNet, if users prefer focusing on only a handful of coarse cell types for their analysis, given their well-established capability of providing cell segmentation masks.

Regarding the fact that the cell types are to some extent continuous and many unknown/novel types are critical for phenotype in cancer, Hist2Cell provides a significantly more comprehensive insight into cellular composition than previous works (2-4 major cell types) as it focuses on predicting fine-grained (up to 80 cell types) and reference-rich cell types, providing the best solution so far for this problem. In addition, Hist2Cell has the potential for novel cell type detection by integrating the posthoc out-of-distribution detection or uncertainty estimation techniques [52, 53], which could be potential future work.

Hist2Cell has a good generalization to unseen datasets, as verified on the external breast cancer datasets and the more diverse and non-standard TCGA datasets. Preliminary experimental results also show that Hist2Cell has the potential to transfer the knowledge learned on one organ to another (from breast cancer tissue to human lung tissue in our study) As different tissues contain an intersection of fine-grained cell types and share similar morphological features, an interesting direction for future work would be to explore the commonalities and differences among different spatial transcriptome datasets with aligned cell type annotation and build a generalized foundation model for pan-cancer analysis.

## 4 Methods

### 4.1 Data Description

The dataset comprising healthy human lung samples was obtained and processed as described in a prior study [6]. This dataset includes 11 slices from tissue sections encompassing the human trachea, bronchi, and both upper and lower parenchyma from 4 patients, totaling 20,770 spots. Visium Spatial Transcriptomics (ST) at 10× magnification was conducted on these slices, and Hematoxylin and Eosin (H&E) images produced during the Visium protocol were captured at a ×20 magnification. The study identified 80 fine-grained cell types and estimated their absolute spatial abundances in the Visium ST data using the cell2location method (version 0.1) [12]. The her2st dataset [27] comprised HER2-positive breast tumors from 8 individuals (Patients A-H). From each individual, either 3 adjacent or 6 evenly spaced sections were obtained (totaling *n* = 36 sections), which were then subjected to Spatial Transcriptomics (ST) [54]. In total, this dataset encompasses 13,620 spots, each with corresponding spatial transcriptomics data. For imaging, the her2st dataset was analyzed using the Metafer VSlide system at ×20 magnification. The STNet dataset [21] included samples from 23 breast cancer patients. For each patient, the dataset contained three microscope images of slides with H&E-stained tissues and their corresponding spatial transcriptomics data, comprising a total of 30,655 spots. The biopsies were collected and processed in accordance with the methods outlined in previous work [54]. Our large-scale study section utilized 565 slides from 565 BRCA patients, (346 for BRCA HER2+, 182 for BRCA-TNBC and rest for other subtypes) and 487 slides from 487 LUSC patients sourced from The Cancer Genome Atlas (TCGA) repository. In line with the approach of previous studies [21], although the bulk RNA-seq for each patient was not performed on the same tissue as the H&E-stained slides, it was utilized as the representative average RNA-seq data for each slide.

### 4.2 Data Preprocessing

#### Spatial Mapping of Cell Types for Breast Cancer Datasets

We employed the fine-grained spatial cell abundance, as estimated from spatial transcriptomics data, as ground truth for training and evaluating Hist2Cell. To maintain consistency with the healthy human lung dataset, we utilized the publicly available Cell2location method to estimate the fine-grained spatial and bulk cell abundances for the her2st, STNet, and TCGA datasets, respectively. The scRNA-seq data from the Human Breast Cell Atlas (HBCA) [45] served as the reference for gene expression signatures of various cell types. This single-cell transcriptomics dataset profiles 714,331 cells from 126 women, uncovering abundant pericyte, endothelial, and immune cell populations, alongside a wide diversity of luminal epithelial cell states. This makes it an excellent reference for studying mammary biology and breast cancer pathology. Specifically, this dataset offers annotations for 39 distinct cell types, encompassing both breast cancer (*n* = 144, 285) and normal breast cells (*n* = 570, 046).

#### H&E Image Cropping

For the healthy human lung, her2st, and STNet datasets, we cropped 224 × 224 image patches from the whole-slide images. These patches were centered on the spatial transcriptomics spots to predict the fine-grained cell type abundances for each spot. For the breast cancer slides from the TCGA repository, we processed the whole-slide images into patches containing substantial tissue amounts, adhering to the pipeline established in previous studies [55]. Initially, candidate 224 × 224 patches were cropped from the whole-slide images without overlap, under a ×10 magnification. Subsequently, patches predominantly consisting of background were discarded. A pixel within a patch was classified as background if its mean RGB value was less than or equal to 220 (indicating a color close to white). A patch was excluded if over 75% of its pixels were categorized as background. Each selected patch was treated as one spot in the TCGA slides. The central pixel coordinates of each spot were utilized for graph construction, visualization, and downstream analysis in our study.

#### Color Normalization

To integrate multiple training/testing samples from diverse sources, such as the TCGA and ST datasets, we employ structure-preserved color normalization techniques, as proposed by Vahadane et al. [56]. This method effectively normalizes color variations that typically arise from different microscopes or scanners and inconsistencies in slide preparation methods. By selecting a representative slide from the ST dataset as a reference, we standardize the color profiles of all training images to this target, thereby reducing one of the most prominent sources of batch effects in histological analyses.

### 4.3 Hist2Cell Design and Training

Hist2Cell is constructed by leveraging the advantages and addressing the limitations of previous works focused on predicting gene expression from histology images, incorporating their key findings. Current approaches can be divided into two groups: (1) global-based methods and (2) local-based methods. In global-based methods, the core concept involves using Whole Slide Images (WSIs) to simultaneously predict an expression map for all coordinates. This enables the model to consider spatial associations between spots. However, this method suffers from severe data scarcity, making models prone to overfitting. Additionally, processing the entire WSI incurs in a high computational cost. Conversely, local-based methods rely solely on the visual information available at each coordinate for predicting gene expression. Training on spot patches, local-based methods benefit from abundant data for deep learning training. Nevertheless, such methods do not take into consideration characteristics such as the vicinity of the patch or long-range interactions, resulting in sub-optimal performance. Previous studies have also discovered that images with similar features tend to exhibit similar gene expression patterns, regardless of their location within the tissue [57, 58]. Building on these findings, we propose employing k-hop subgraphs as the fundamental unit of our framework and utilize a Graph-Transformer architecture to learn both local context and distant relations, thus benefiting from the advantages of local-based and global-based methods without succumbing to their respective limitations.

To be specific, the key idea of our approach is to execute a minibatch of local subgraphs sampled from a WSI instead of using an entire WSI as a single data element. This strategy conserves computational resources and mitigates the overfitting risks commonly associated with using large WSIs, which makes our framework suitable for super-resolved cellular maps inference and improves the generalization of our framework. To better model the relationships within each subgraph and among different subgraphs, we employed a Graph-Transformer architecture. The Graph Attention layers facilitate message passing between spots and their spatial neighbors, learning the local spatial context within each subgraph, while the Transformer layers calculate the attention score among any pair of spots despite the physical distance, modeling long-range correlations existing among different subgraphs. In Supplement Fig 5, we validate the effectiveness of leveraging both local and global relations from the tissue, results showing that both the GNN part and the Transformer part contribute to the performance increase of Hist2Cell. Specifically, we further investigate the effectiveness of GNN over typical spatial smoothing in predicting fine-grained cell types. Previous research in this domain such as [24] has demonstrated the benefit of learning associations between spatially adjacent spots when predicting spatial gene expression from histology images. We propose that similar benefits extend to predicting fine-grained cell type abundances, which are inherently correlated with spatial gene expressions. GNNs are particularly suited for this task because they do not simply perform spatial smoothing; they learn advanced feature representations by integrating adjacency information with the attributes of each node (spot). This capability allows GNNs to capture complex structures and relationships within spatial data more effectively than traditional smoothing techniques. Empirical results in Supplement Fig 6 validate our hypothesis and demonstrate the effectiveness of GNN in understanding and modeling the complex spatial interactions that are crucial for the accurate prediction of fine-grained cell type abundance. More details of the framework are illustrated in the subsequent sections.

#### Spatial Graph Construction

As illustrated in Fig. 1b, we abstract the whole-slide image (WSI) into a spatial graph. Within this spatial graph, each node corresponds to a 224 × 224 image patch, centered on specific spatial spots, targeted for spatial cell abundance prediction. Utilizing the spatial coordinate information of all nodes, we calculate the distances between nodes using the Euclidean distance. For each node, we select *k* nearest nodes based on the smallest Euclidean distances, where *k* = 6 for 10× Visium data and *k* = 8 for ST data [54]. Leveraging the spatial neighbor relationships, we construct an undirected graph, denoted as *G* = {*V, E*}, to represent the corresponding whole-slide image. Here, the node set *V* represents the image patches, and the edge set *E* indicates the spatial neighbor relationships among these patches.

#### Local Subgraph Sampling and Embedding

The core concept involves initially sampling all the nodes required for computation. We begin by randomly sampling a minibatch of central nodes, denoted as *B*, from the spatial graph *G*. Subsequently, using a pre-defined radius *k* for local subgraphs, we gather all necessary nodes for computation, denoted as *B^k^* = *B* ∪ *N*_1_(*B*) *. . .* ∪ *N_k_*(*B*). We use the notation *N_i_*(*B*) to denote a deterministic function that specifies the set of all *i*-hop neighbor nodes of all center nodes in *B*. In this study, we use *k* = 2 for both training and inference of Hist2Cell. Together with *B^k^*, we obtain the spatial neighbor relationship *E^k^* among all nodes in *B^k^* as the corresponding subset of edge set *E*. To embed the image patches within *B^k^*, we employ a feature extractor, represented as *I*(·), to learn fine-grained spatial heterogeneity in cell type abundances within the tissue. This yields a low-dimensional embedding 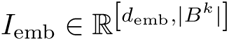, where *d*_emb_ is the dimension of extracted node embedding, and *I*_emb_ denotes all node embedding of *B^k^*. In our experiments, we used a ResNet-18 [59] model with pre-trained ImageNet [60] weights as its initialization. The ResNet-18 model comprises 18 convolutional layers with residual connections, followed by a fully connected layer. We remove the fully connected layer and use the feature embedding before it as the node embedding in our study. Throughout the training of Hist2Cell, the parameters of the ResNet-18 model are optimized in an end-to-end fashion.

#### Short-range Local Message Aggregation

Adhering to the principles of the Graph-Transformer architecture, Hist2Cell initially employs a Graph Attention layer to learn local representations of sampled subgraphs *B^k^*. As demonstrated in prior studies [61, 62], the Graph Neural Network (GNN) component of a Graph-Transformer architecture derives node representations from neighborhood features. This neighborhood aggregation in GNN plays a crucial role in learning local and short-range correlations among graph nodes. Specifically, we implement the GATv2 layer [63] for message propagation and aggregation. The message propagation and aggregation of node *i* in *B^k^*are defined as:

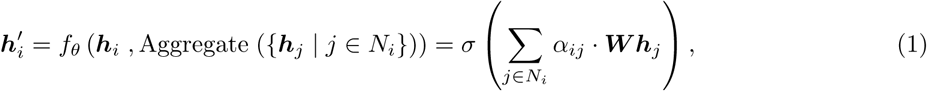

where *h_i_* and *h_j_* are the node embeddings of node *i* and node *j* from the previous layer, and *N_i_* denotes the set of all spatial neighbor nodes of node *i* in *B^k^*. Thus, the GATv2 layer computes a weighted average of the transformed features of neighbor nodes (followed by a nonlinearity *σ*) as the new representation ***h****_i_*^′^ of node *i*, using normalized attention coefficients *α_ij_*. For attention coefficient calculation, a scoring function *e* : ℝ*^d^*^emb^ × ℝ*^d^*^emb^ → ℝ computes scores for each edge (*j, i*), indicating the importance of neighbor node *j*’s features to node *i*:

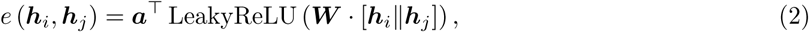

where ***a*** ∈ ℝ^2*d*^*^′^* , ***W*** ∈ ℝ*^d′^*^×*d*^ are learned during the training process, and ∥ denotes vector concatenation. The attention scores are normalized across all neighboring nodes in *N_i_* using the softmax function, and the attention function is defined as:

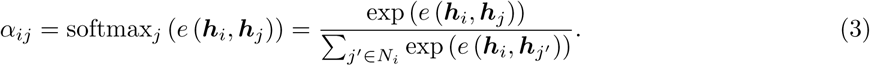

#### Long-range Global Correlation Learning

In Hist2Cell, apart from the local aggregation of neighbor nodes, we also learn the long-range correlations among all sampled nodes in *B^k^*. The underlying premise is that spots exhibiting similar morphological features should have similar cell abundances, regardless of their distance within the tissue. Hist2Cell employs the Transformer [64] to learn these long-range correlations, treating each node in *B^k^* as tokens. The core component of the Transformer is its multi-head attention mechanism, denoted as:

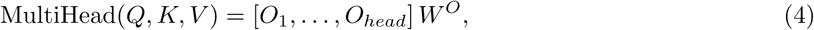

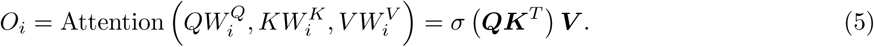

Within this formula, *head* represents the number of parallel attention layers, and *σ* denotes an activation function. Utilizing the components mentioned above, the Transformer layer is defined as:

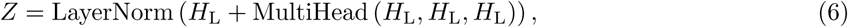

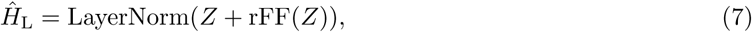

where *H*_L_ is the features of *B^k^* fed into the *L*-th Transformer layer, rFF is a row-wise feedforward layer, LayerNorm is layer normalization [65], and *H̑_j_* represents the transformed features of all spots in *B^k^* after considering the long-range correlations.

#### Multi-scale Feature Fusion and Prediction

In the final stage, Hist2Cell fuses multi-scale features from the feature extractor, the Graph Attention layers, and the Transformer layers using a residual block as shown in Fig.1b, which can be denoted as:

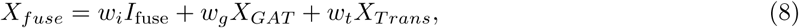

where *X_fuse_*, *I*_fuse_, *X_GAT_* , and *X_T rans_* are the fused features, features from the feature extractor, Graph Attention part, and Transformer part, respectively, and *w_i_*, *w_g_*, and *w_t_* are pre-defined weights for feature fusion. Subsequently, we apply Multi-Layer Perceptron (MLP) heads to each node in *B^k^* for spatial cell abundance prediction:

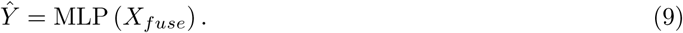

The Mean Squared Error (MSE) loss is employed to optimize the parameters in Hist2Cell using gradient descent. As the training of Hist2Cellis conducted on randomly sampled subgraphs, and the loss term is calculated among all sampled nodes in *B^k^*, the inference phase is slightly different for Hist2Cell. During inference, every spot in the whole-slide image is sampled as the center node of subgraphs. For more reliable cell abundance prediction, we only use the predictions from the center nodes of the subgraphs and merge them into slide-level spatial cell abundances.

### 4.4 Cell-cell Colocalization Analysis via Spatial Correlation

Cell-cell colocalization analysis, leveraging spatial cell abundance data, was conducted using the spatial correlation statistic, bivariate Moran’s R, as implemented in the SpatialDM toolbox [28]. Specifically, to evaluate the spatial correlation between cell type pairs, we calculated the bivariate Moran’s R statistic using SpatialDM. The bivariate Moran’s R statistic, originally proposed for detecting ligand-receptor pairs exhibiting significant spatial co-expression, facilitates reliable analysis of cell-cell communication in spatial transcriptomics data. This statistic is an extension of techniques previously utilized in geography [66]. Utilizing the available spatial abundance data for various cell types in our study, we directly calculated the bivariate Moran’s R statistic between different pairs of cell types. Specifically, the statistic is defined as:

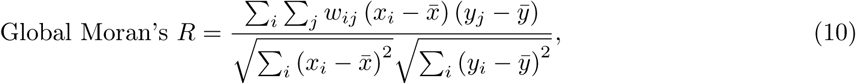

where *x_i_* and *y_j_* denote the cell abundances of cell types *x* and *y* at spots *i* and *j*, respectively. The spatial weight matrix computation employs a Radial Basis Function (RBF) kernel with element-wise normalization:

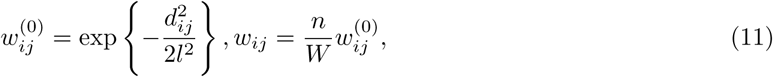

where *d_ij_* represents the geographical distance between spots *i* and *j* (Euclidean distance on spatial coordinates), *W* is the sum of *w_ij_*^(0)^, and *n* is the total number of spots. To assess the efficacy of Hist2Cell in predicting spatial cell abundance, we utilized clustermaps colored according to the bivariate Moran’s R statistic. These were derived from both the ground truth spatial cell abundances and the predictions made by Hist2Cell and STNet. The pattern similarities between the clustermaps of the ground truth and those generated by Hist2Cell underscore the effectiveness of our method.

### 4.5 Survival Analysis

Survival analysis plays a pivotal role in clinical scenarios, offering critical insights into outcomes such as mortality, disease progression, and recovery by examining survival data that spans time periods encompassing censored time points [67]. With the advent of deep learning, leveraging whole slide images (WSIs) and patients’ molecular information for survival prediction has shown promising results in various cancer types. Features extracted from these two modalities could utilized to train a survival model, such as Cox regression models, for predicting patient outcomes. The concordance index is usually employed to evaluate the accuracy of the survival risk prediction.

#### Problem Formulation

Building upon prior work [39], we applied a deep learning-based survival prediction technique that discretizes the survival time into intervals, with each interval being represented by a distinct output neuron. This approach mitigates the requirement for large mini-batch sizes, enabling the model to be trained using individual observations. Specifically, for right-censored survival data, we construct a discrete-time survival model by dividing the continuous time axis into distinct intervals: [*t*_0_*, t*_1_) , [*t*_1_*, t*_2_) , [*t*_2_*, t*_3_) , [*t*_3_*, t*_4_), defined by the quartiles of uncensored patients’ survival times (in months) in each TCGA cohort. The discrete event time for each patient, indexed by *j*, with a continuous event time *T_j_*, _cont_ , is then described as:

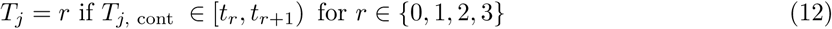

Given the discrete-time label for the *j*^th^ patient as *Y_j_*, and the patient’s slide-level feature vector **h**_final_ *_j_*, the network’s final layer applies a sigmoid activation function, modeling the hazard function as:

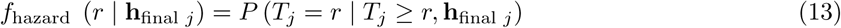

which relates to the survival function by:

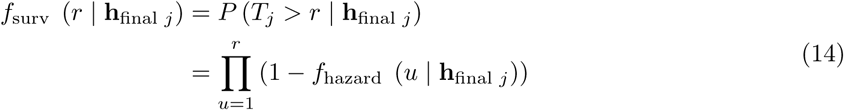

The model parameters are updated during training via the log-likelihood function of a discrete survival model [68], considering the censorship status of each patient ( *c_j_* = 1 if the patient survives beyond the follow-up period, and *c_j_* = 0 if the patient dies within the recorded event time *T_j_* ):

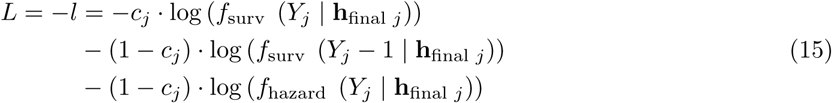

In training, we further enhance the model by up-weighting uncensored patient cases using a weighted sum of *L* and *L*_uncensored_ :

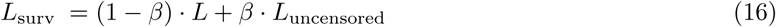

The second term in the loss function, specific to uncensored patients, is computed as:

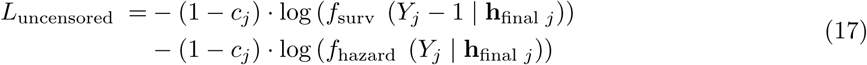

Specifically, for the patient’s slide-level feature vector **h**_final_ *_j_* , we have slide-level transcriptional fine-grained cell abundances, slide-level morphological features, and patient-wise bulk RNA-seq in our study. To obtain the slide-level transcriptional fine-grained cell abundances and morphological features, we applied average and attention-based pooling among Hist2Cell’s patch-level predictions and patch-level features extracted by HIPT [41] or UNI [43]. For the patient-wise bulk RNA-seq, we select the top 1000 highly variable expressions from the patients’ whole profile.

#### Concordance Index

The concordance index (C-index) is a widely used metric to evaluate the predictive accuracy of survival models, particularly in scenarios involving censored data. It measures the ability of a model to correctly rank the survival times of individuals, providing an estimate of the discriminative power of the model. Formally, the C-index can be defined as the proportion of all possible pairs of individuals whose predicted survival times are correctly ordered, given that one individual has a longer observed survival time than the other. For a pair of individuals (*i, j*) with true survival times *T_i_* and *T_j_*, and corresponding predicted risks *T̑_i_* and *T̑_j_* , the C-index is computed as:

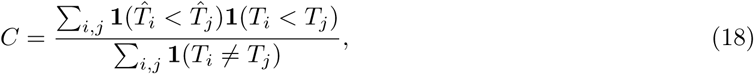

where **1**(·) is the indicator function that returns 1 if the condition is true and 0 otherwise. A C-index value of 0.5 indicates a model with no predictive power, equivalent to random guessing, while a value of 1.0 indicates perfect concordance between predicted and actual outcomes. In the presence of censored data, the C-index is adjusted by considering only those pairs where one individual is known to have a longer survival time than the other.

### 4.6 Super-resolved Spatial Cell Abundance

In our study, to produce higher-resolution cellular map is equivalent to predict the relative cell abundances for more positions on the slide. For instance, for the 2× resolution results in Fig 6, we predict the relative cell abundances for sub-spots, whose number is four times the original spots in the slide. Under this objective, our approach is able to produce high-resolution cellular maps as we could predict the relative cell abundances for a certain position based on its morphological features from the 224×224 image patches centered on this position and its neighboring image patches. For instance, for the 2× resolution results in Fig 6, for the original resolution, we crop 224×224 spot patches using a radius of about 112-pixel distances on the tissue, then, for producing the 2× resolution results, we will crop 224×224 sub-spot patches using a radius of about 56-pixel distances. We want to point out that this sliding-window strategy is reasonable due to two reasons: (1) Hist2Cell is trained with 224×224 image patches and it is likely to work as we did not change the input image size when producing high-resolution cellular maps; (2) producing high-resolution from 224×224 would not affect the reliability and the biological utility of the predicted fine-grained cell abundance as it is actually a relative value that high related to the gene expression in a specific location.

Particularly, we also noticed that Hist2Cell share some principles with HistToGene [20], as both methods involve dense sliding window sampling. However, we note two major differences including: (1) The model architectures are fundamentally different. HisToGene relies solely on a Vision Transformer to model entire slides, whereas Hist2Cell combines a convolutional feature extractor with a Graph-Transformer model. This design enables Hist2Cell to extract morphology features for each spot, capture spatial contextual relations between neighboring spots, and understand representation similarities among distant tissue regions. With these capabilities, Hist2Cell demonstrates stronger representation power than HistToGene, making it more suitable for decoding complex transcriptional information from histology images; and (2) Hist2Cell has a GPU memory advantage. While HistToGene requires GPU memory that scales quadratically with the number of spots, Hist2Cell only needs a constant amount of GPU memory by using subgraphs as basic computational units. This is particularly important in super-resolution settings, where increased resolution results in a higher spot count per WSI. Indeed, our study provides 16× super-resolved results, whereas HistToGene only presents 2× super-resolution results in their study.

As illustrated in Fig.5.a, we could infer high-resolution cellular maps from the spatial graphs generated under high-resolutions in two ways:

#### Impute from Low-resolution Cellular Map

Given a low-resolution cellular map as priors, Hist2Cell can be fine-tuned on this map to make predictions on a super-resolved spatial graph derived from the same whole-slide image. This approach yields higher-resolution spatial cell abundances that are consistent with the low-resolution priors.

#### Predict from H&E Image

Furthermore, Hist2Cell can directly predict super-resolved spatial cell abundances from histology images of human lung and breast tissues, using its parameters optimized on these two tissue sections. These predictions could serve as reliable references, given Hist2Cell’s strong capabilities to overcome batch effects on the external data.

## 5 Data Availability

All datasets employed in this study are previously published and publicly accessible. The healthy lung dataset was downloaded from the GitHub repository at https://5locationslung.cellgeni.sanger.ac.uk/. The her2st dataset was obtained from https://github.com/almaan/her2st. The STNet dataset was sourced from https://data.mendeley.com/datasets/29ntw7sh4r/5. The TCGA dataset was acquired from the Genomic Data Commons Data Portal at https://portal.gdc.cancer.gov/. The scRNA-seq data from the Human Breast Cell Atlas (HBCA) was downloaded from CELLxGENE at https://cellxgene.cziscience.com/collections/ 4195ab4c-20bd-4cd3-8b3d-65601277e731. Source data for this study are provided alongside this paper.

## 6 Code Availability

The Hist2Cell source code are available in Github (https://github.com/Weiqin-Zhao/Hist2Cell). We have uploaded the processed data and necessary files including the prepossessing code to GitHub. Furthermore, A comprehensive demo is available, providing users with a practical step-by-step tutorial for environment installation, model training, and reproducing the main results of our study.

## Acknowledgement

This work was partially supported by the Research Grants Council of the Hong Kong SAR, China (Project No. 27206123 and T45-401/22-N) and the Hong Kong Innovation and Technology Fund (Project No. ITS/274/22).

## 7 Author contributions

LY and YH conceived and supervised the study. WZ implemented the framwork and performed all data analysis, with support from ZL and XH. WZ, LY and YH wrote the manuscript with inputs from all authors.

## 9 Supplementary

**Supplementary Table 1:** Conceptual comparison between our Hist2Cell with conventional digital pathol-ogy (DPath), HE2RNA/ST prediction works, and XFuse/iStar works.

| Group | Method | Input |  | Label Source |  | Output |  | Generalization |  |
| --- | --- | --- | --- | --- | --- | --- | --- | --- | --- |
|  |  | Histology Image | Spatial Gene | Manual Annotation | Spatial Gene | Cell Abundance | Cell Granularity | Within Section | Cross Patient |
| 1 | Conventional DPath [4, 5] | ✓ | ✗ | ✓ | ✗ | ✓ | 2-4 types | ✓ | ✓ |
| 2 | HE2RNA [69] | ✓ | ✗ | ✗ | ✗ | N/A | N/A | ✓ | ✓ |
|  | ST Prediction [21, 23, 24, 29] | ✓ | ✗ | ✗ | ✓ | N/A | N/A | ✓ | ✓ |
| 3 | XFuse [16], iSTAR [18] | ✓ | ✓ | ✗ | ✓ | N/A | N/A | ✓ | ✗ |
| Ours | Hist2Cell | ✓ | ✗ | ✗ | ✓ | ✓ | up to 80 types | ✓ | ✓ |

**Supplementary Fig 1:**
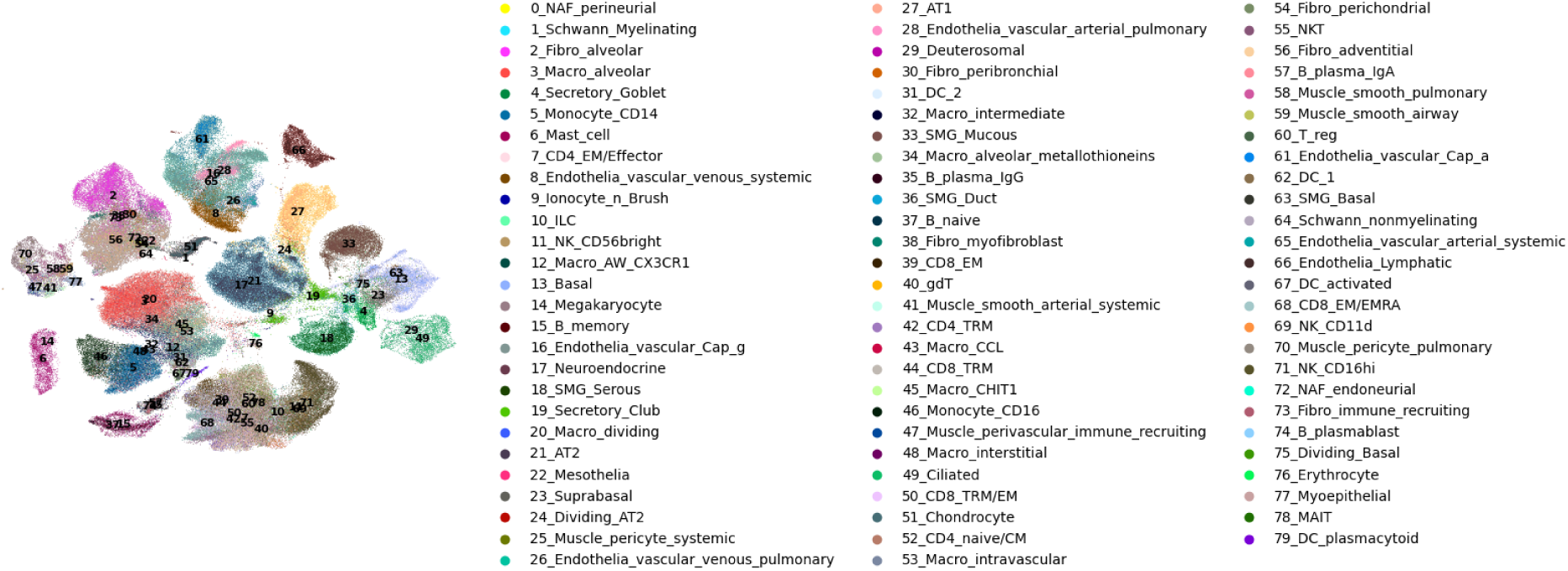
UMAP representation of 80 cell types identified by Louvain clustering on the reference single cell reference dataset of human lung dataset.

**Supplementary Fig 2:**
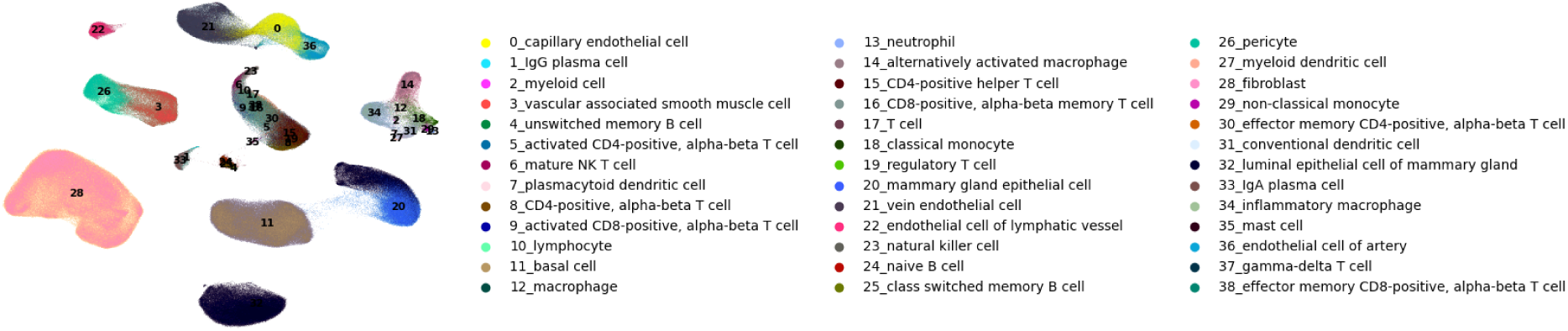
UMAP representation of 39 cell types identified by Louvain clustering on the reference single cell reference dataset of human breast cancer dataset.

**Supplementary Fig 3:**
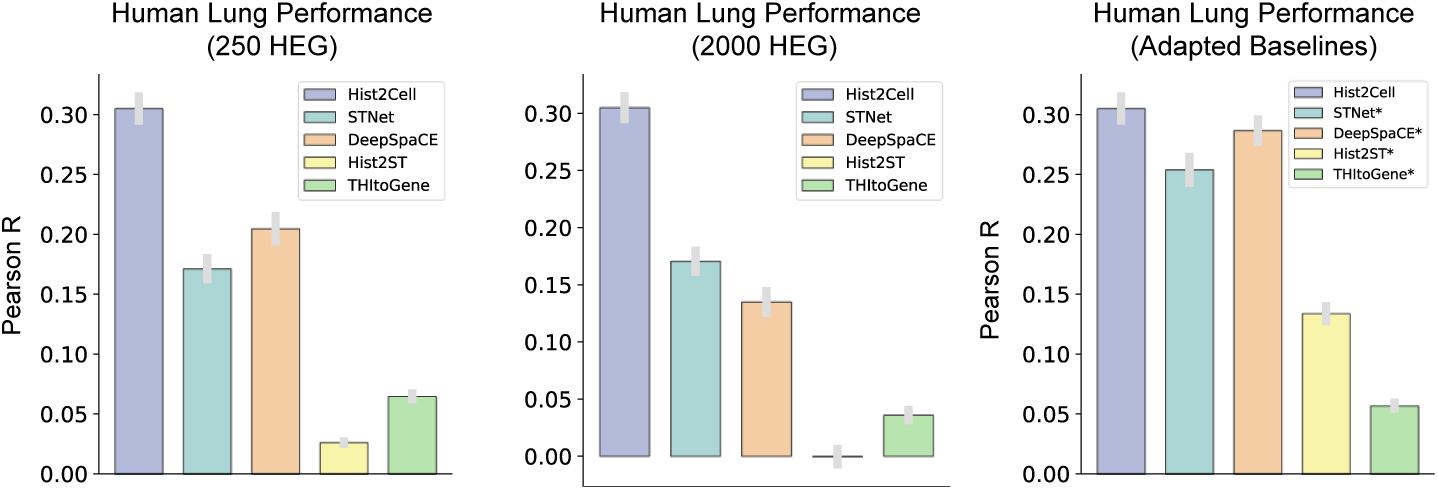
Histogram depicting the average Pearson’s R values for cell abundance prediction in the leave-one-donor-out cross-validation experiment conducted on the healthy human lung dataset. "250 HEG" and "2000 HEG" represent that, for the previous ST prediction baselines, we follow their setup to first predict the 250/2000 highly expressed genes, and then use the predicted highly expressed genes to esti-mate the cell abundance. "Adapted Baselines" represents that, for the previous ST prediction baselines, we adapted them to our one-stage prediction strategy to directly predict the fine-grained cell abundances from the histology image. We use **\*** to distinguish the adapted baselines from the original ones. Results show that Hist2Cell demonstrated more accurate predictions of fine-grained cell type abundances across locations, as previous ST prediction baselines suffer from low performance in fine-grained cell type abundances prediction under both "250 HEG" and "2000 HEG" and setups as their noisy and unstable predictions for a relatively small group of genes. When adapted, the performance of previous baselines increases obviously, showing the efficacy of our one-stage prediction strategy.

**Supplementary Fig 4:**
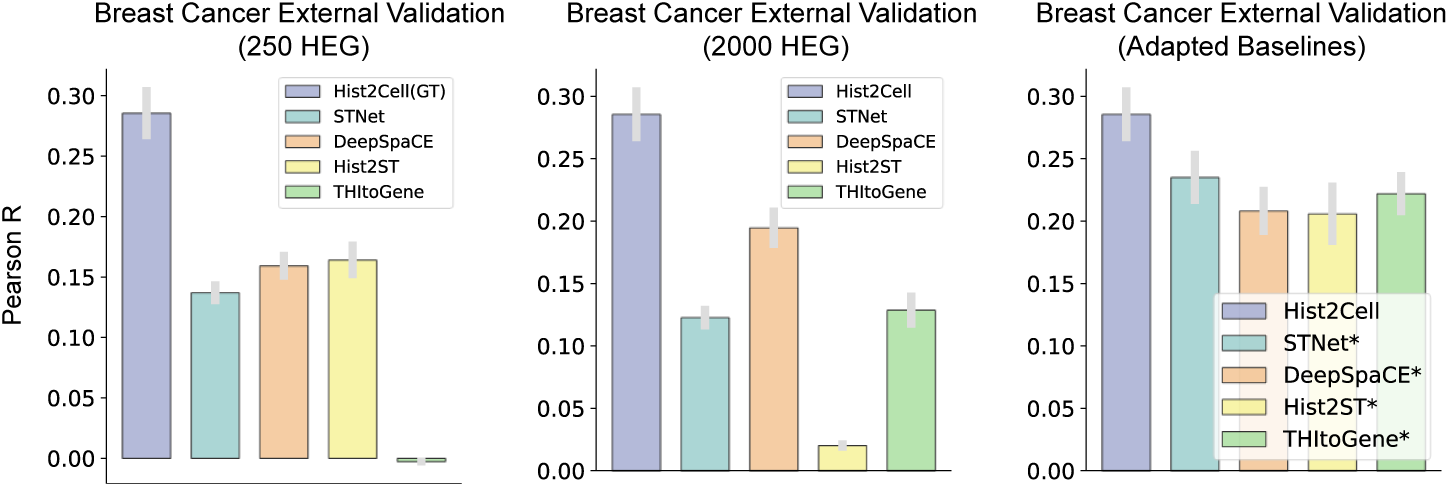
Histogram depicting average Pearson’s R values for cell abundance prediction in the external breast cancer dataset. "250 HEG" and "2000 HEG" represent that, for the previous ST prediction baselines, we follow their setup to first predict the 250/2000 highly expressed genes, and then use the predicted highly expressed genes to estimate the cell abundance. "Adapted Baselines" represents that, for the previous ST prediction baselines, we adapted them to our one-stage prediction strategy to directly predict the fine-grained cell abundances from the histology image. We use **\*** to distinguish the adapted baselines from the original ones. Results show that Hist2Cell demonstrated more accurate predictions of fine-grained cell type abundances across locations, as previous ST prediction baselines suffer from low performance in fine-grained cell type abundances prediction under both "250 HEG" and "2000 HEG" and setups as their noisy and unstable predictions for a relatively small group of genes. When adapted, the performance of previous baselines increases obviously, showing the efficacy of our one-stage prediction strategy.

**Supplementary Fig 5:**
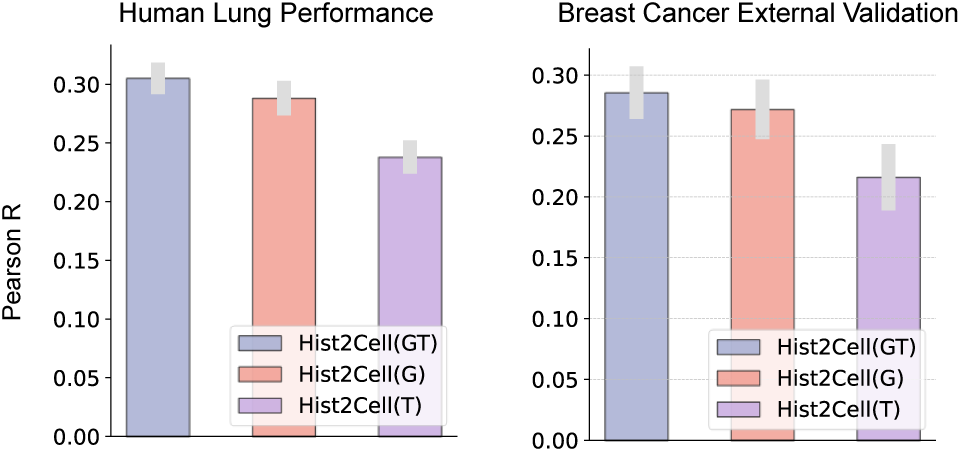
Histogram depicting the average Pearson’s R values for cell abundance prediction for human lung dataset and external breast cancer dataset. "GT" represents the complete Hist2Cell, "G" represents removing the Transformer part of Hist2Cell, and "T" represents removing the GNN part of Hist2Cell. Results show that both the GNN part and the Transformer part contribute to the performance increase of Hist2Cell, validating the efficacy of utilizing both local and global information within the tissue.

**Supplementary Fig 6:**
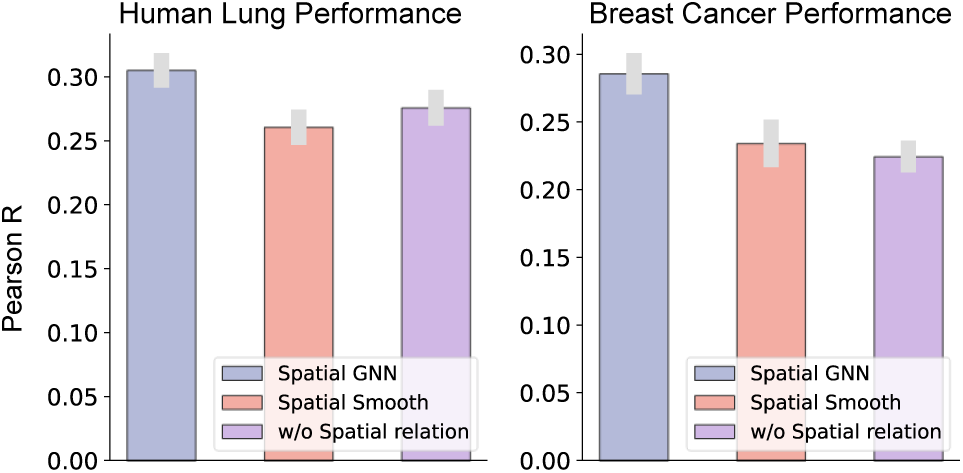
Histogram depicting the average Pearson’s R values for cell abundance prediction for human lung dataset and breast cancer datasets. We compare three different settings: using GNN, using spatial smoothing from STNet and removing the spatial relation modeling in Hist2Cell.

**Supplementary Fig 7:**
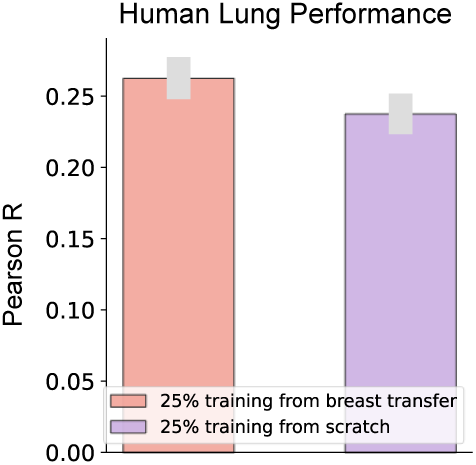
Histogram depicting the average Pearson’s R values for cell abundance prediction for the human lung dataset under low data regime (25% training data). Training from breast transfer denotes that we use the model weights trained on the breast cancer dataset as the initialization. Results show that the knowledge of predicting fine-grained cell type abundances could be transferred across different tissues via the pre-trained model parameters.

**Supplementary Fig 8:**
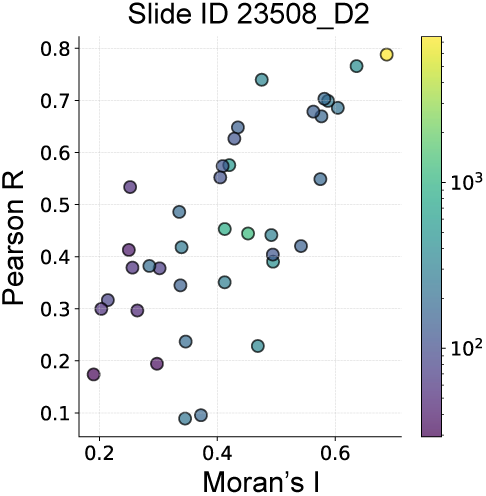
Scatter plot for one test slide illustrating the relationship between spatial auto-correlation (Moran’s *I*; x-axis) and Hist2Cell’s prediction performance (y-axis) in the healthy human lung dataset. Hist2Cell excelled in predicting cell types exhibiting higher spatial auto-correlation on the external breast cancer dataset.

**Supplementary Fig 9:**
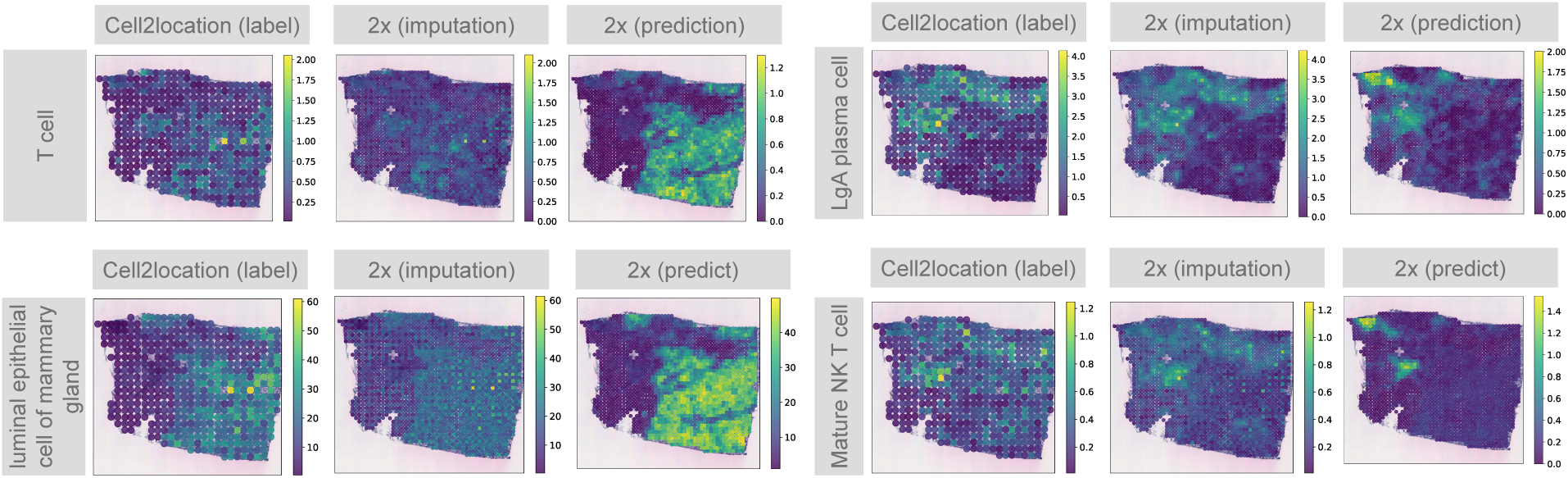
Hist2Cell provides super-resolved spatial cell abundances with more accurate alignment to tumor/normal annotation than STNet. Visualizations comparing low-resolution ground truth cell abundance with both imputed and directly predicted high-resolution results by Hist2Cell. Both approaches provide a more detailed mapping in concordance with the tumor/normal anno-tation shown in Fig. 4.c.

**Supplementary Table 2:** Comparison average C-indices of the cox regression models predicting survival of three different cancer subtypes: LUSC, BRCA-TNBC, and BRCA-HER2+. C-indices were calculated from the test sets of a 10-fold Cross-Validation experiment. The Cox regression models used the predicted cell abundances of Hist2Cell and the image feature extracted by HIPT/UNI, and the bulk-RNA seq of the patients as inputs, respectively. Top performance in the histology image-based models is shown in **bold**.

| Model | Pre-training Slide No. | C-index(%) |  |  |  |
| --- | --- | --- | --- | --- | --- |
|  |  | BRCA-HER2+ | BRCA-TNBC | LUSC | Average |
| HIPT [41] | 10,678 | 57.82 | 71.57 | 70.58 | 66.65 |
| UNI [43] | 100,426 | 64.05 | 74.35 | **71.75** | 70.05 |
| Hist2Cell | 114 (BRCA) / 34 (LUSC) | **67.89** | **74.43** | 71.59 | **71.30** |
| Bulk RNA-seq | N/A | 70.75 | 70.60 | 72.05 | 71.13 |

**Supplementary Fig 10:**
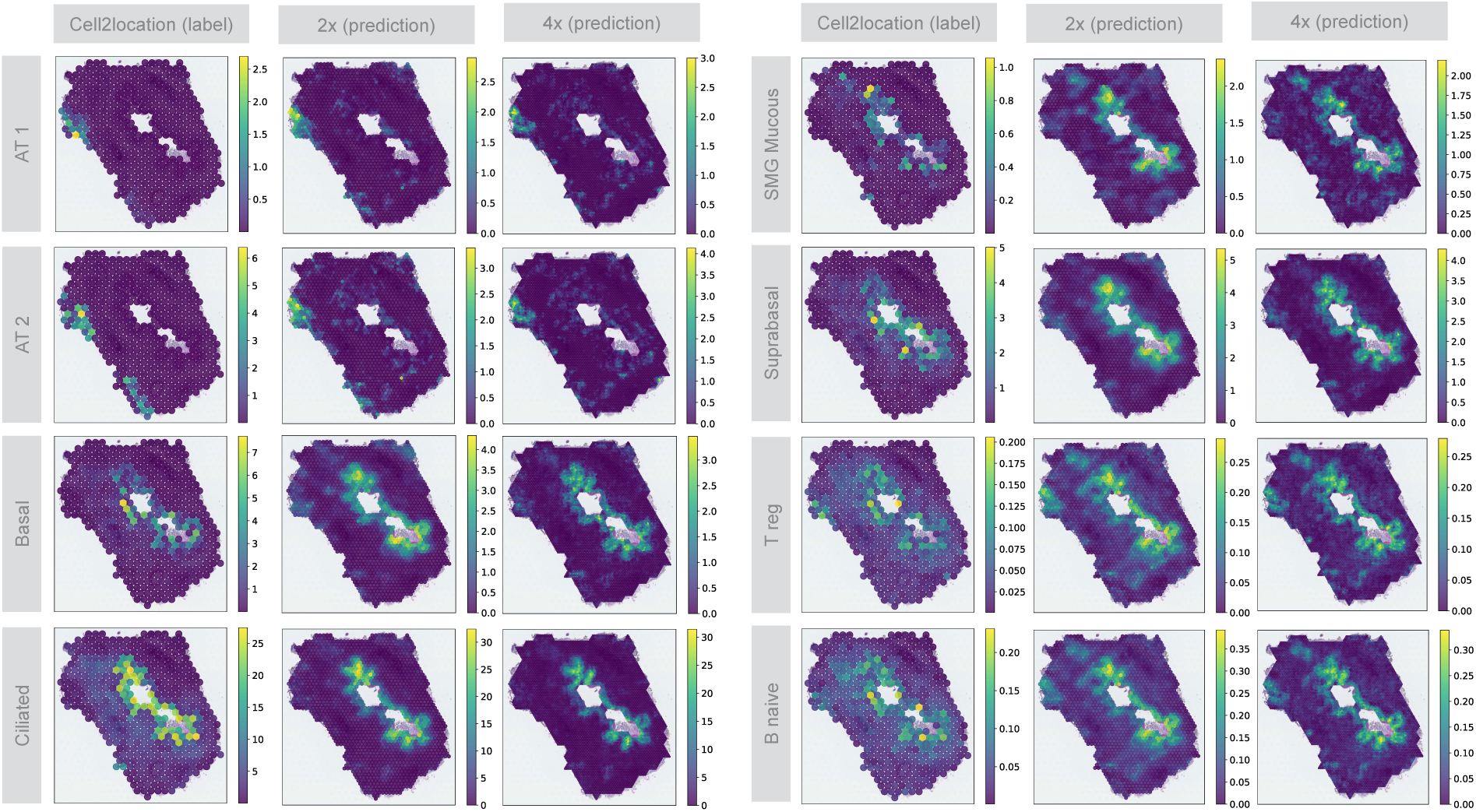
Visualizations comparing low-resolution ground truth cell abundance with directly predicted 2× and 4× high-resolution results by Hist2Cell. Higher resolutions provide more detailed mapping in concordance with the manual annotation shown in Fig 3.a.

**Supplementary Fig 11:**
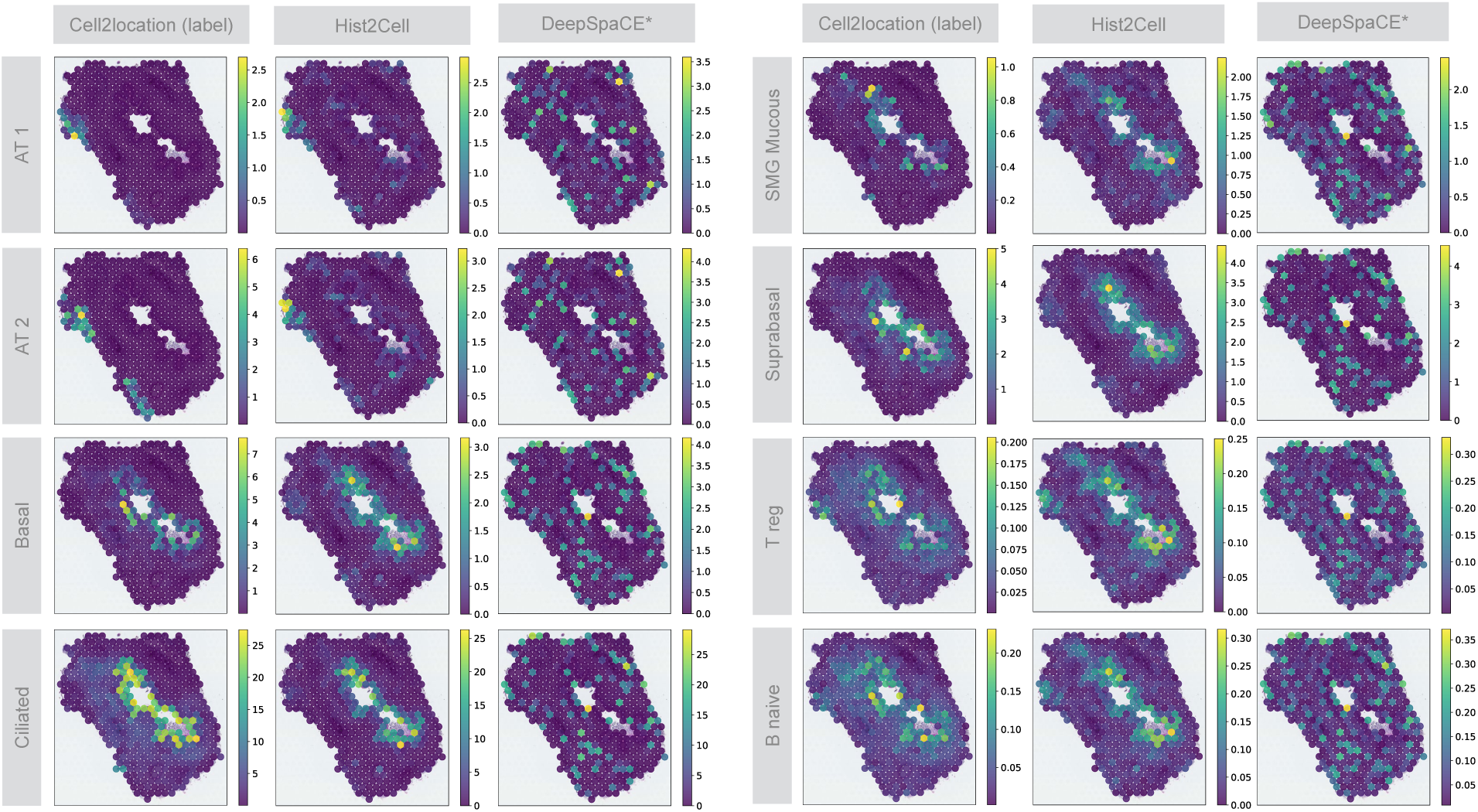
Hist2Cell provide more accurate alignment to manual annotation than DeepSpaCE*. The visualizations comparing the spatial cell abundances as determined by ground truth, Hist2Cell, and DeepSpaCE* predictions for select key cell types. Hist2Cell show less false positives than DeepSpaCE* when comparing to both ground truth and manual annotation in Fig.3.b.

**Supplementary Fig 12:**
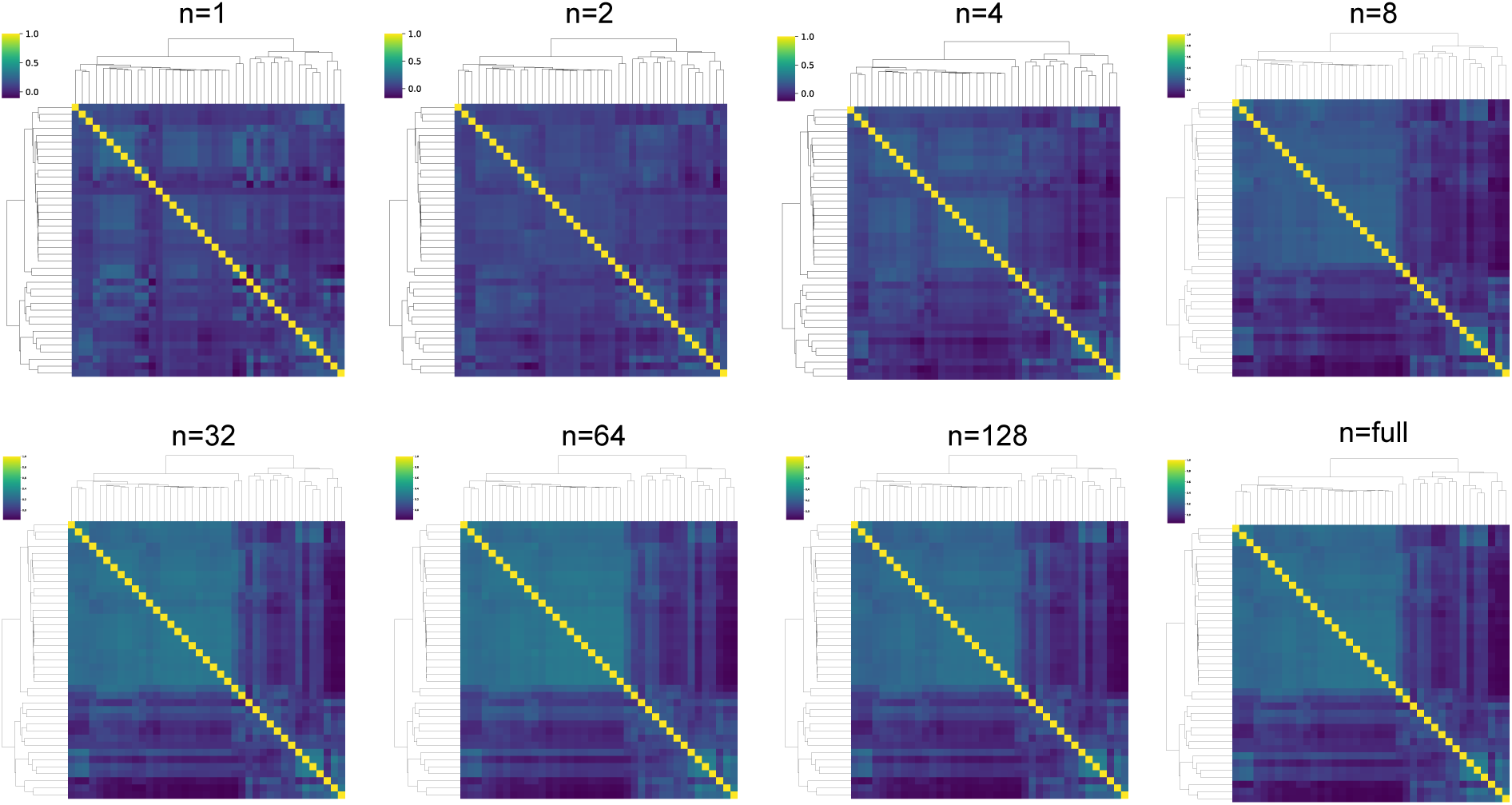
Hist2Cell provides more stabilized consensus analysis with larger sam-ple sizes. Consensus clustermaps generated by Hist2Cell using varying numbers of samples from the TCGA breast cancer cohort. *n* denotes the number of randomly sampled slides, with *n* = full indicating the use of the entire cohort. The consensus clustermaps become more stable as the number of sampled slides increases.

**Supplementary Fig 13:**
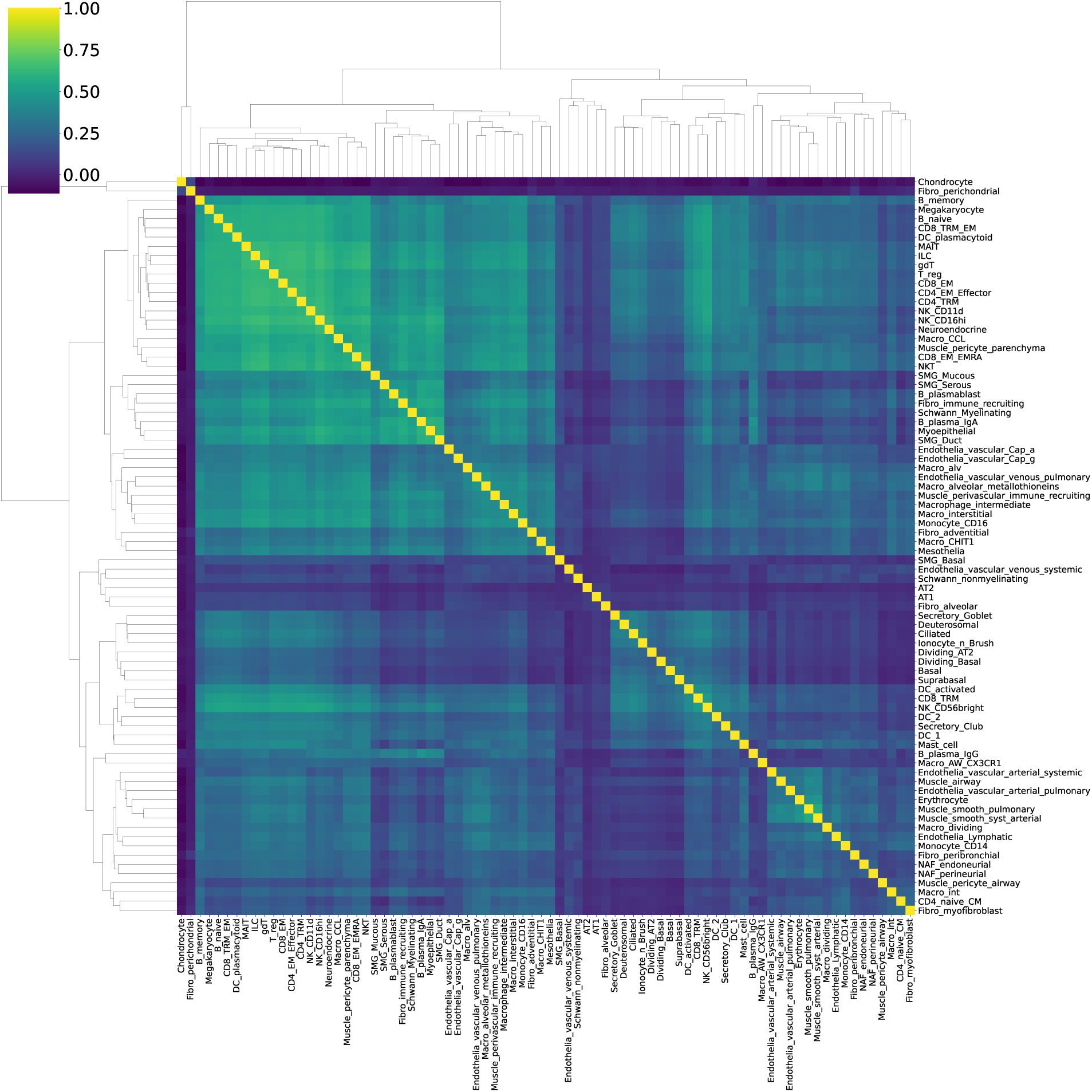
Detailed Human lung Moran’s R Clustermaps calculated from ground truth cell type abundances (cell2location algorithm) in Fig.2e.

**Supplementary Fig 14:**
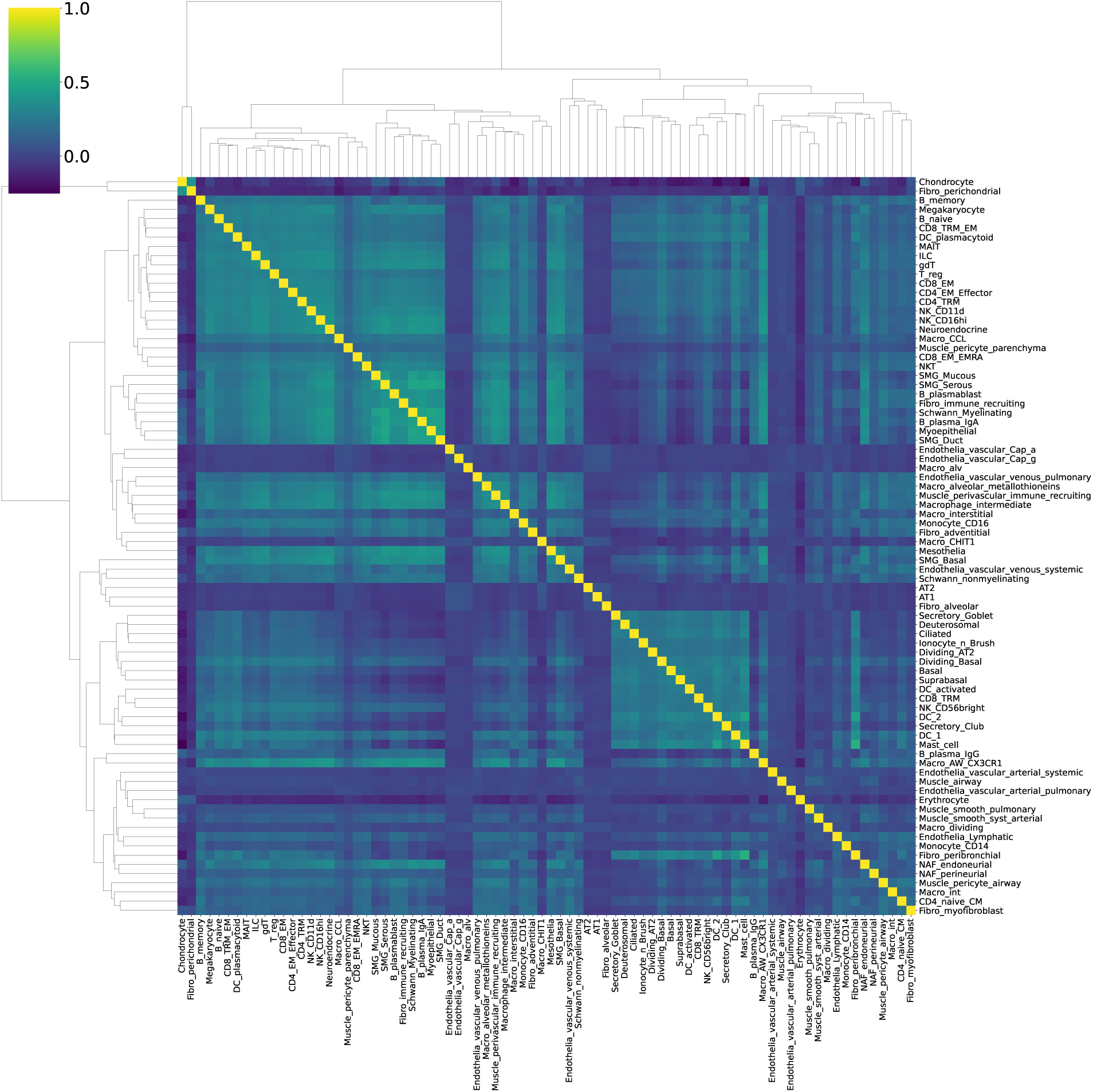
Detailed Human lung Moran’s R Clustermaps calculated from Hist2Cell predic-tions in Fig.2e.

**Supplementary Fig 15.**
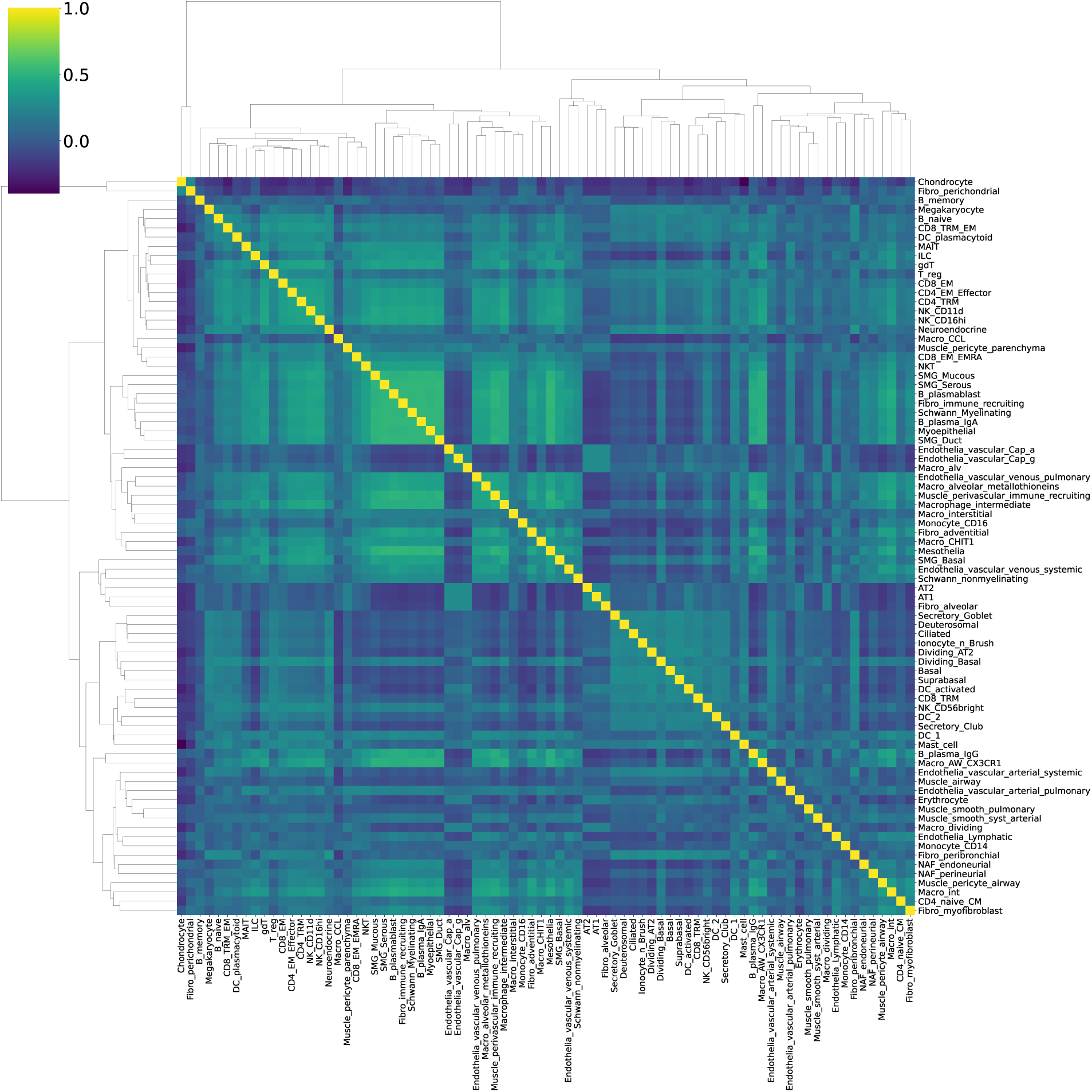
: Detailed Human lung Moran’s R Clustermaps calculated from DeepSpaCE* predictions in Fig.2e.

**Supplementary Fig 16:**
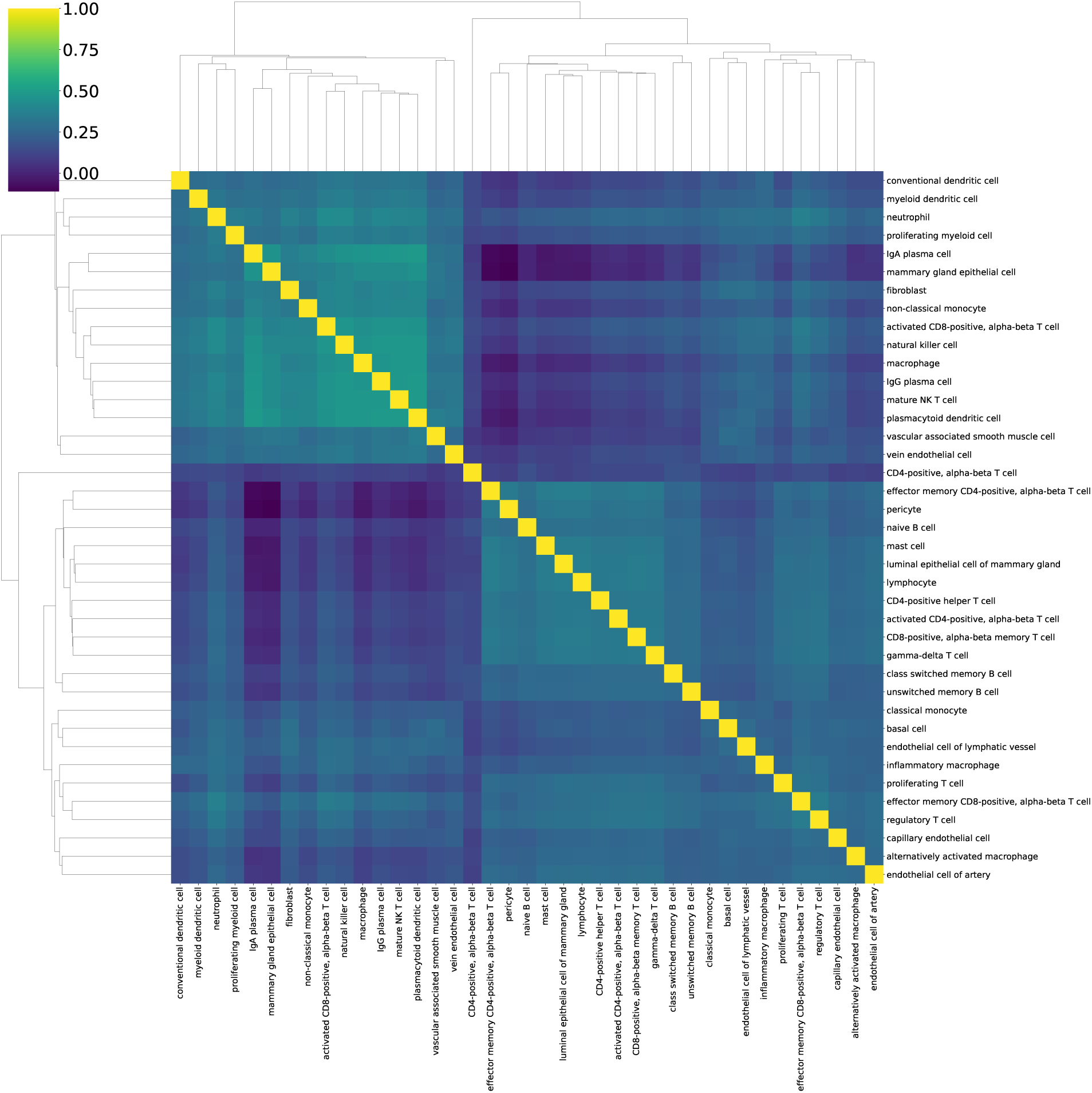
Detailed Human breast cancer Moran’s R Clustermaps calculated from ground truth cell type abundances (cell2location algorithm) in Fig.4.e.

**Supplementary Fig 17:**
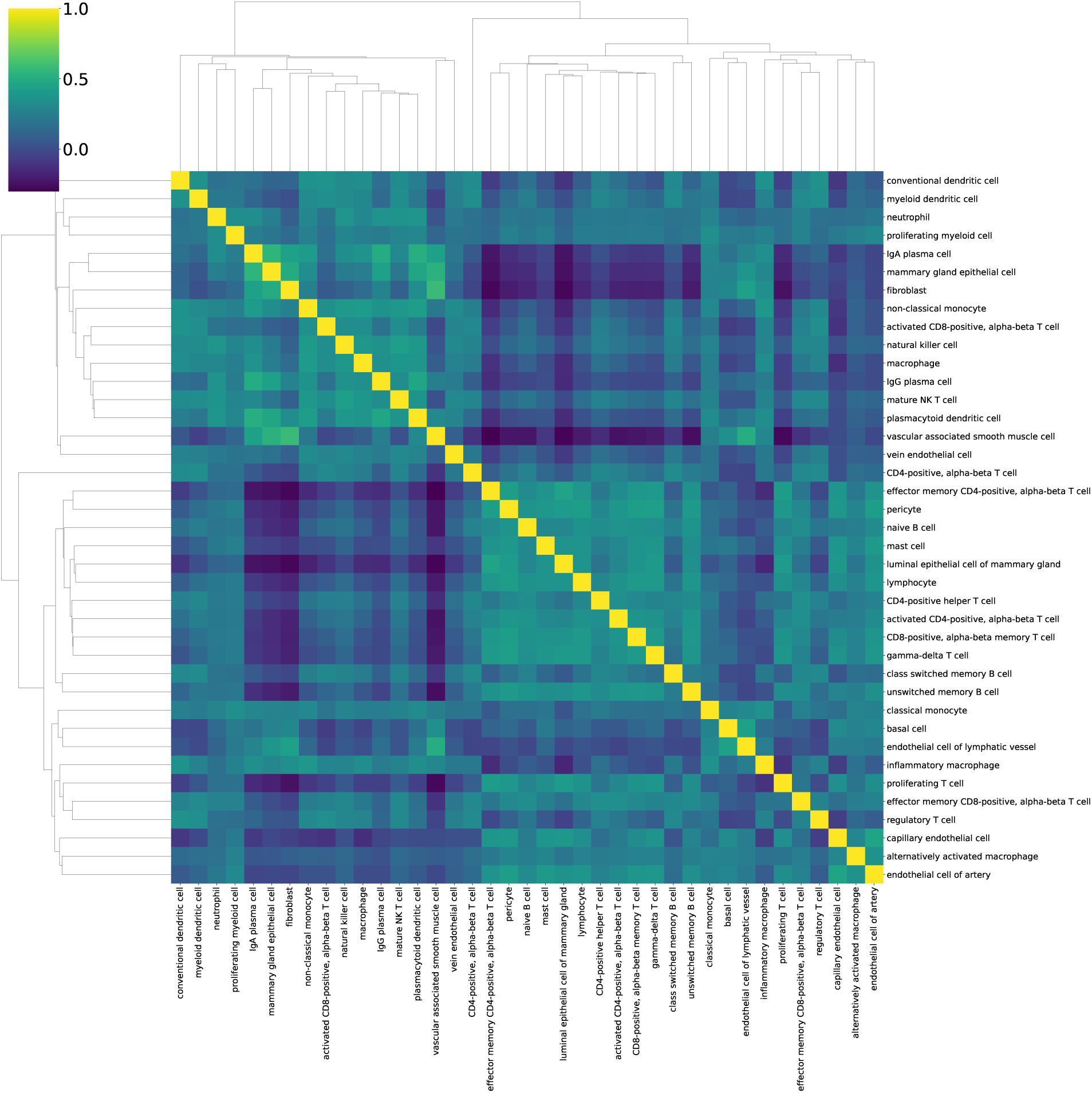
Detailed Human breast cancer Moran’s R Clustermaps calculated from Hist2Cell predictions in Fig.4.e.

**Supplementary Fig 18:**
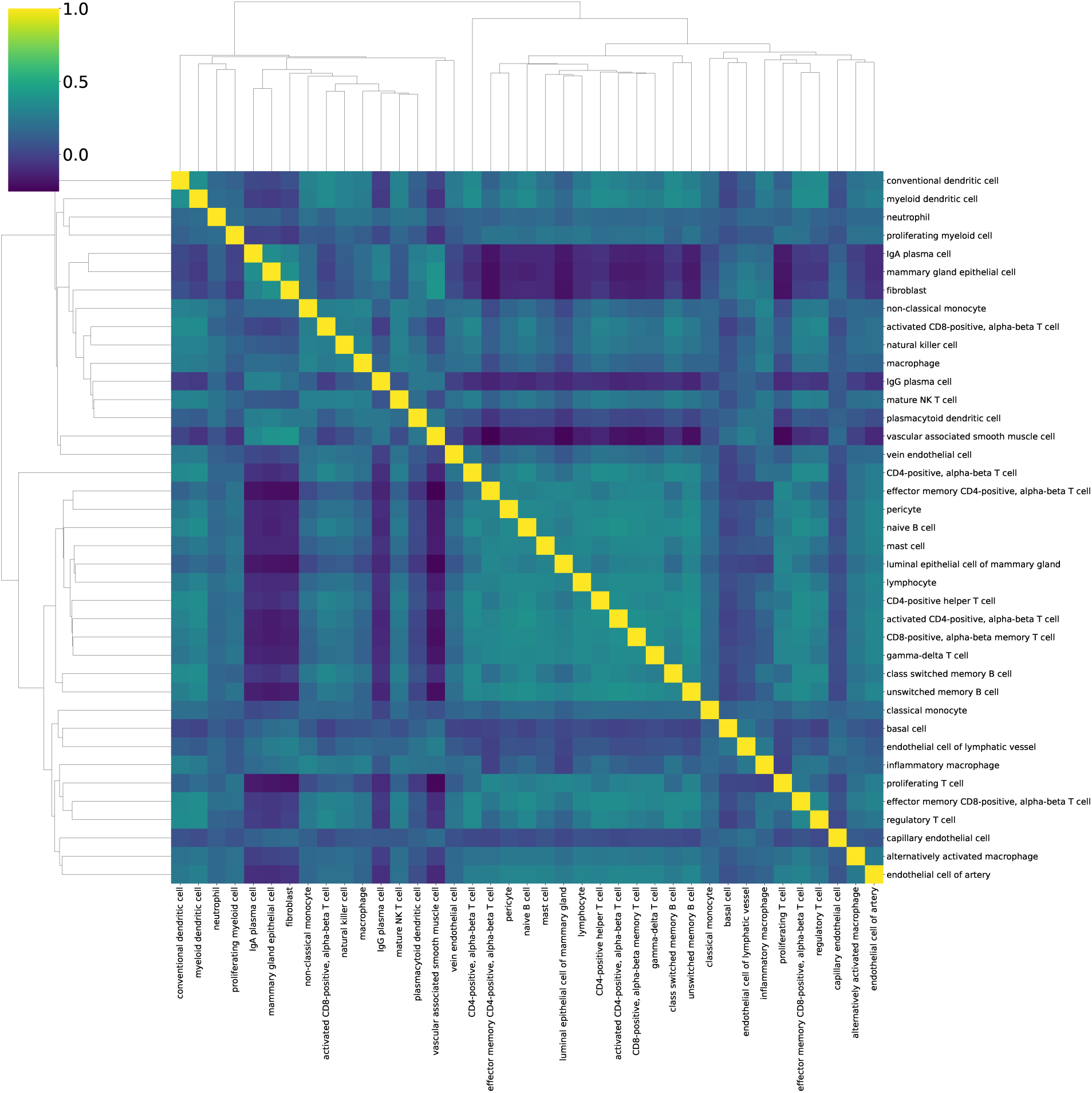
Detailed Human breast cancer Moran’s R Clustermaps calculated from STNet* predictions in Fig.4.e.

**Supplementary Fig 19:**
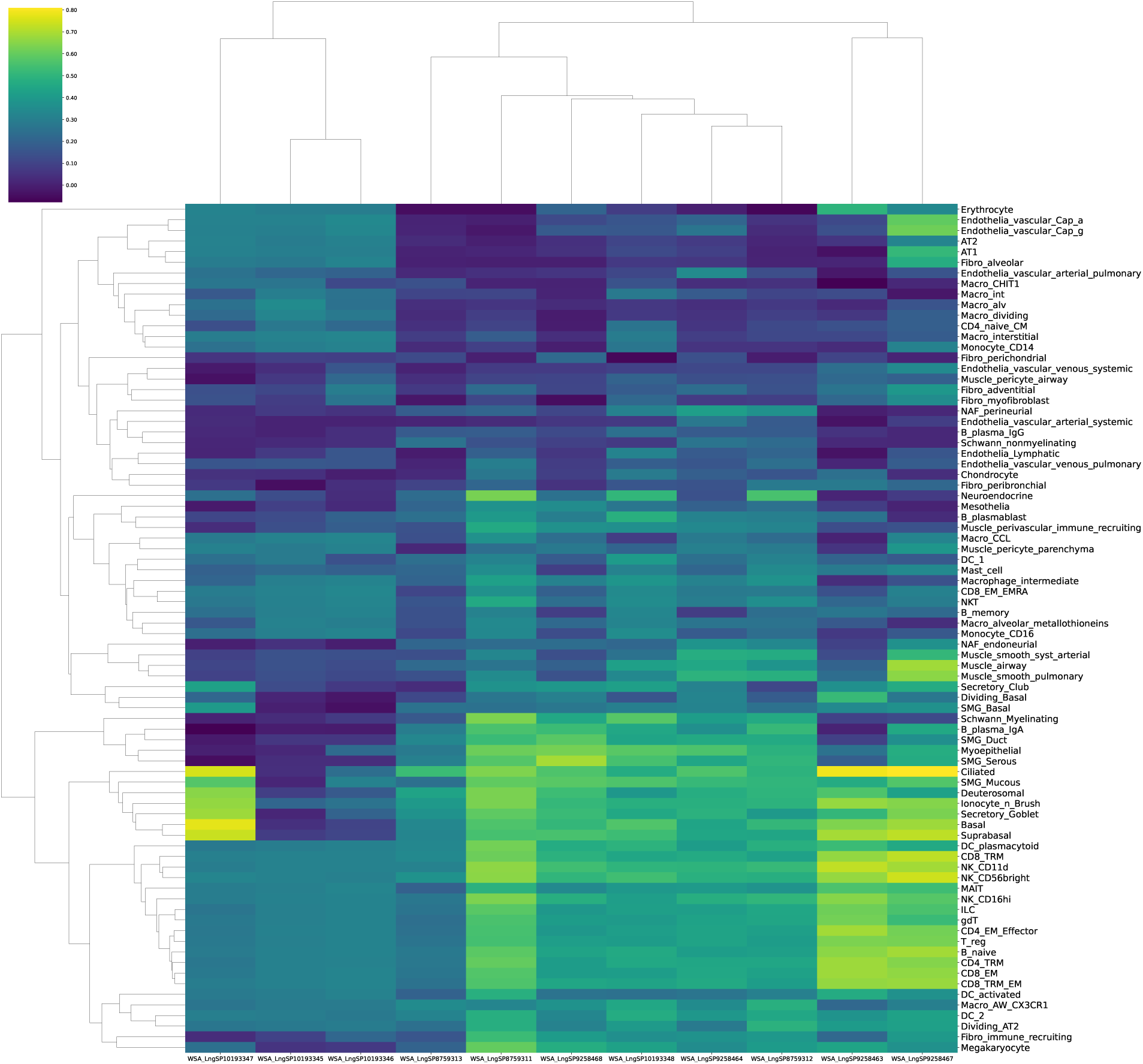
Clustermap colored by the Pearson R from Hist2Cell for different cell types (y-axis) on different slides (x-axis) during testing in the human lung dataset experiments. The top 30% cell types across 11 slides have a mean Pearson R 0.50.

**Supplementary Fig 20:**
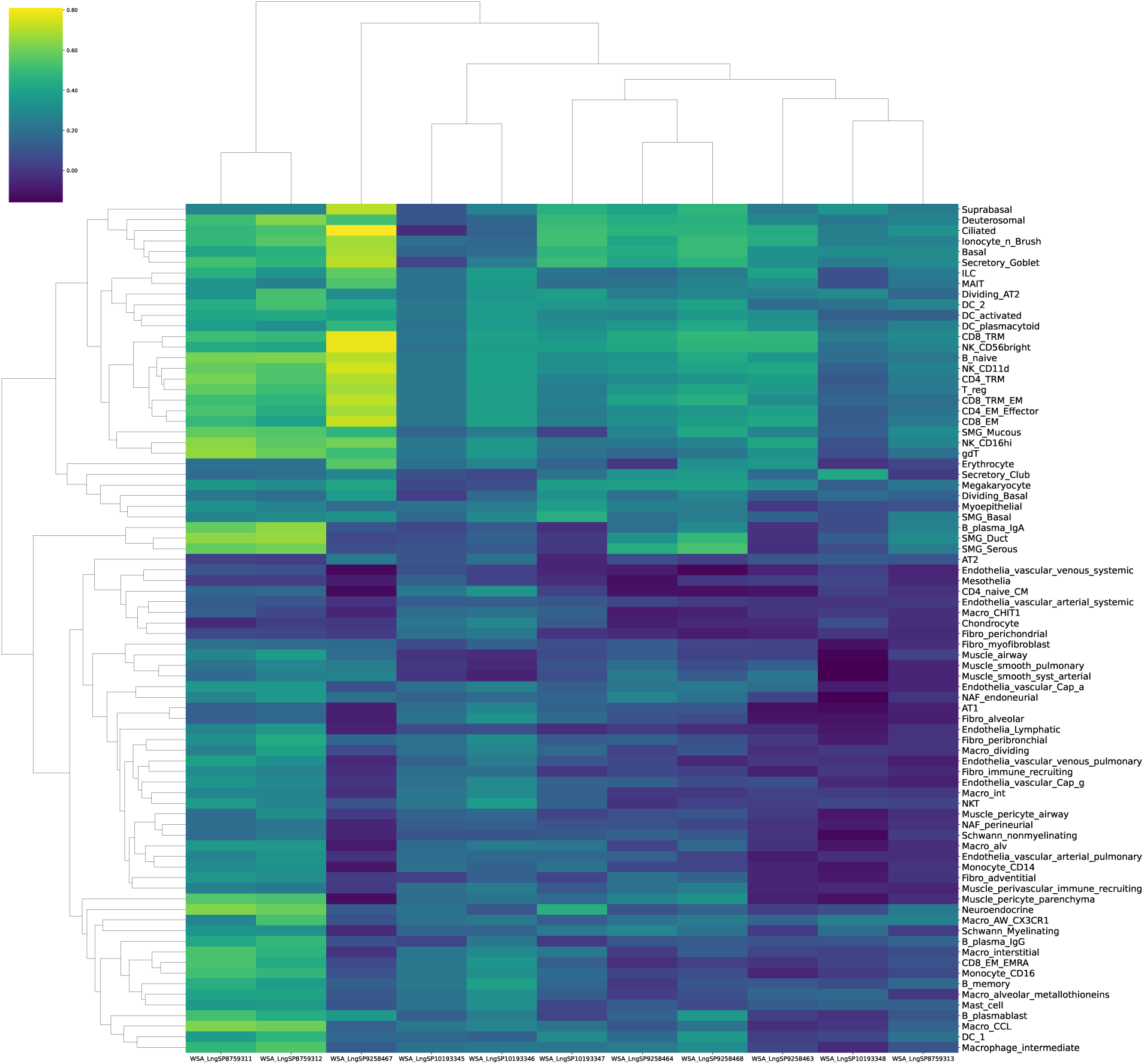
Clustermap colored by the Pearson R from DeepSpaCE baseline for different cell types (y-axis) on different slides (x-axis) during testing in the human lung dataset experiments. The top 30% cell types across 11 slides have a mean Pearson R 0.43.

**Supplementary Fig 21:**
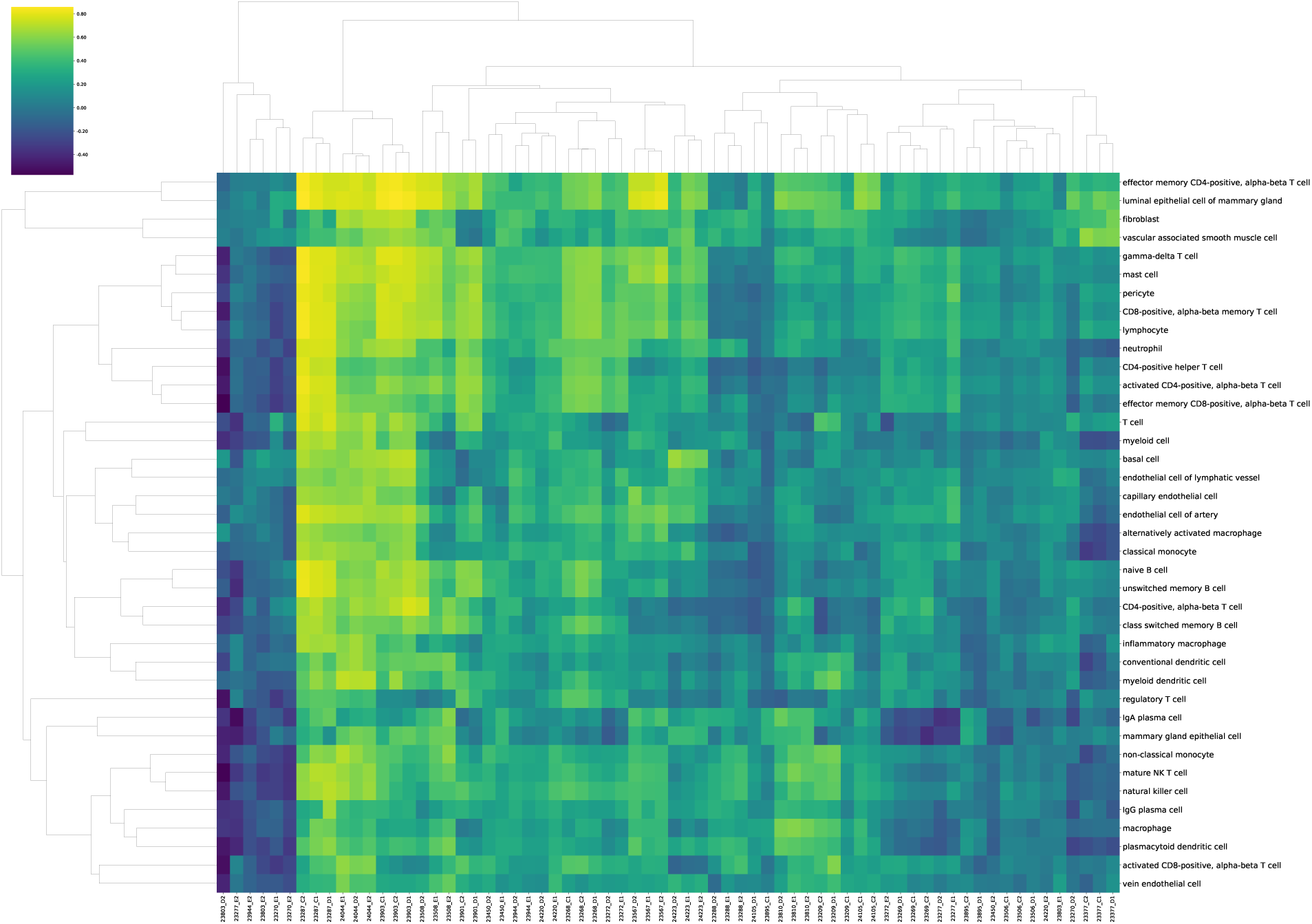
Clustermap colored by the Pearson R from Hist2Cell for different cell types (y-axis) on different slides (x-axis) during testing in the external breast cancer dataset experiments. The top 30% cell types across 68 slides have a mean Pearson R 0.54.

**Supplementary Fig 22:**
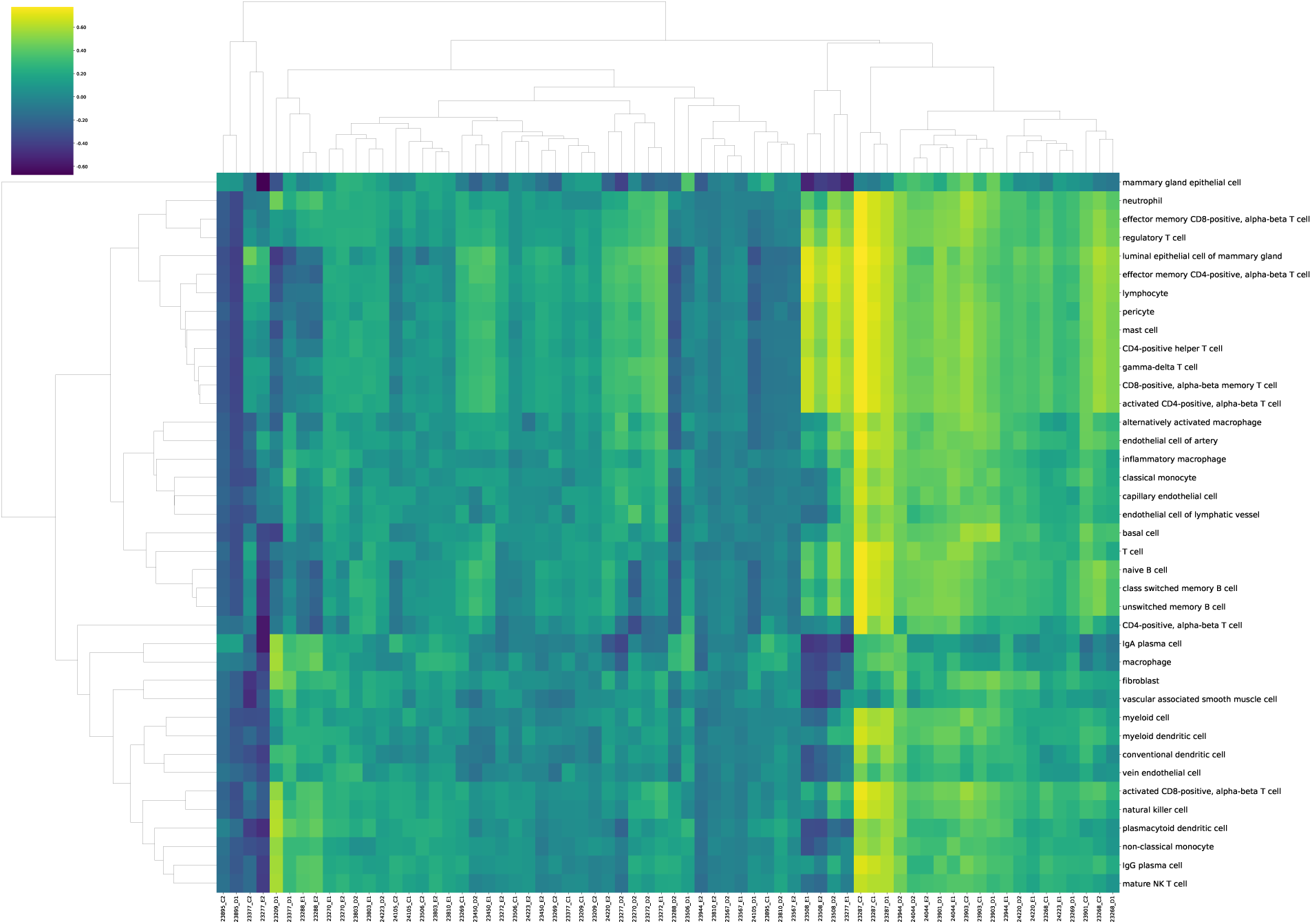
Clustermap colored by the Pearson R from DeepSpaCE for different cell types (y-axis) on different slides (x-axis) during testing in the external breast cancer dataset experiments. The top 30% cell types across 68 slides have a mean Pearson R 0.44.

**Supplementary Fig 23:**
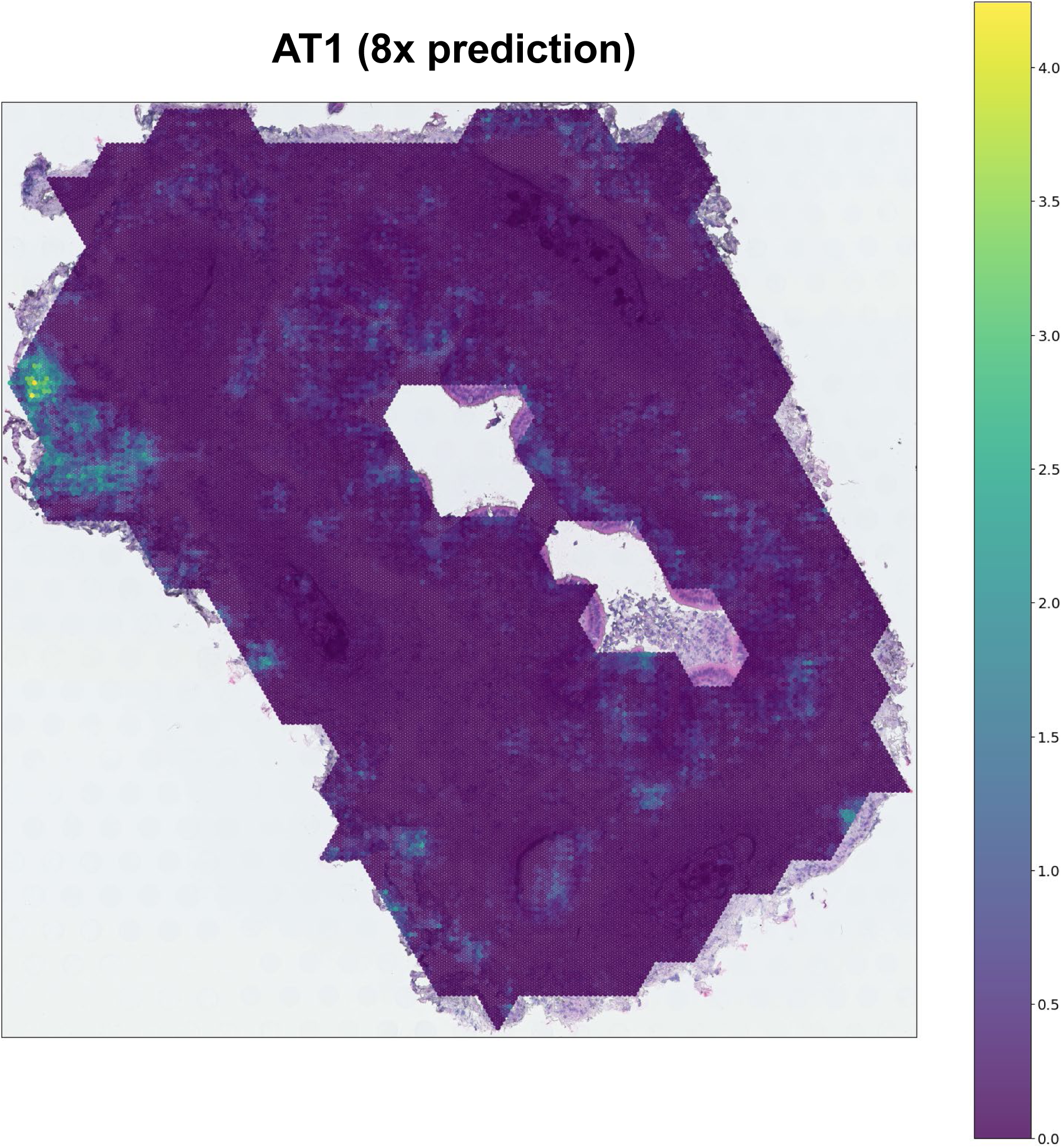
Visualization of the directly predicted 8× high-resolution results of AT1 cells by Hist2Cell.

**Supplementary Fig 24:**
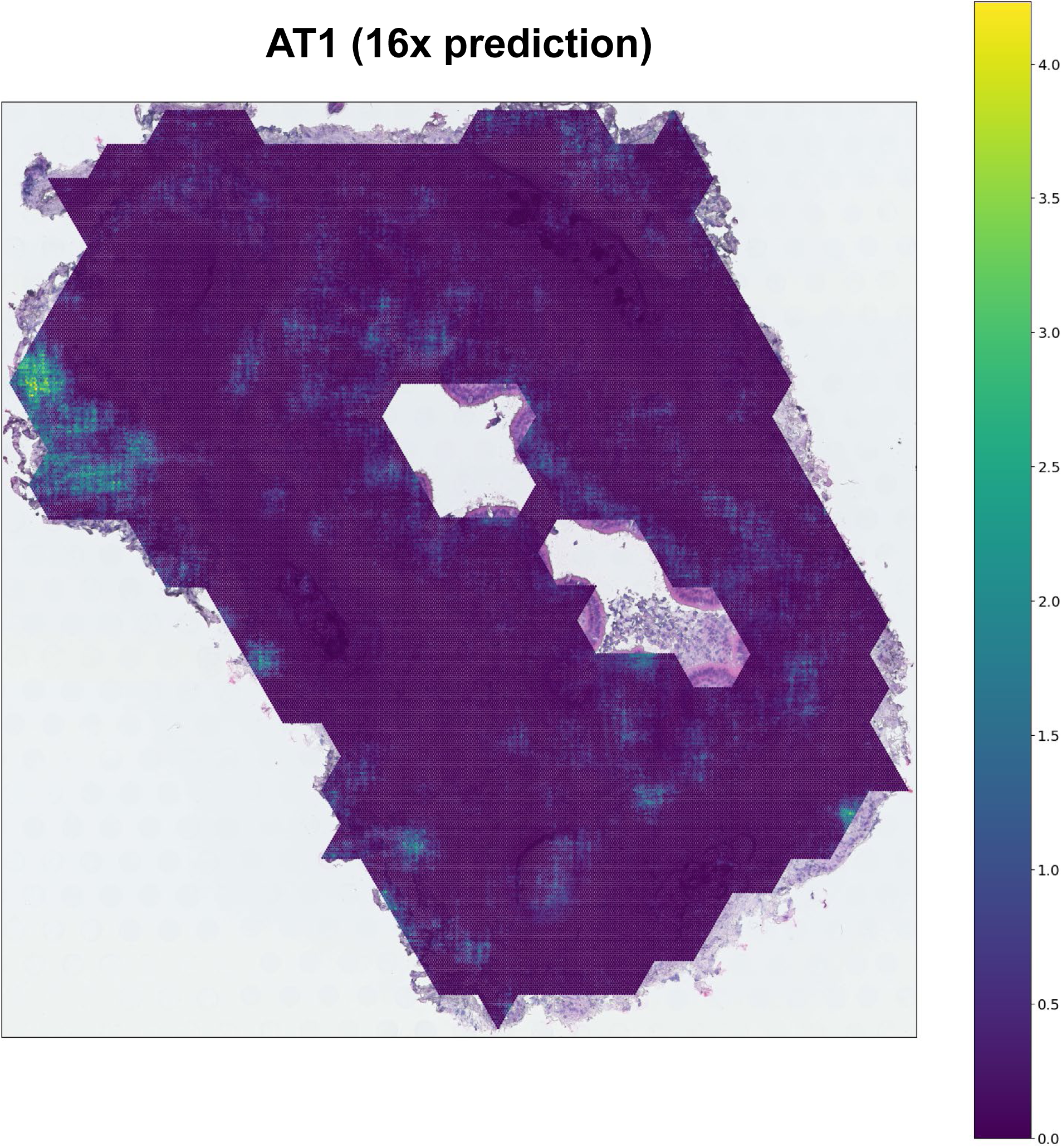
Visualization of the directly predicted 16× high-resolution results of AT1 cells by Hist2Cell.

**Supplementary Fig 25:**
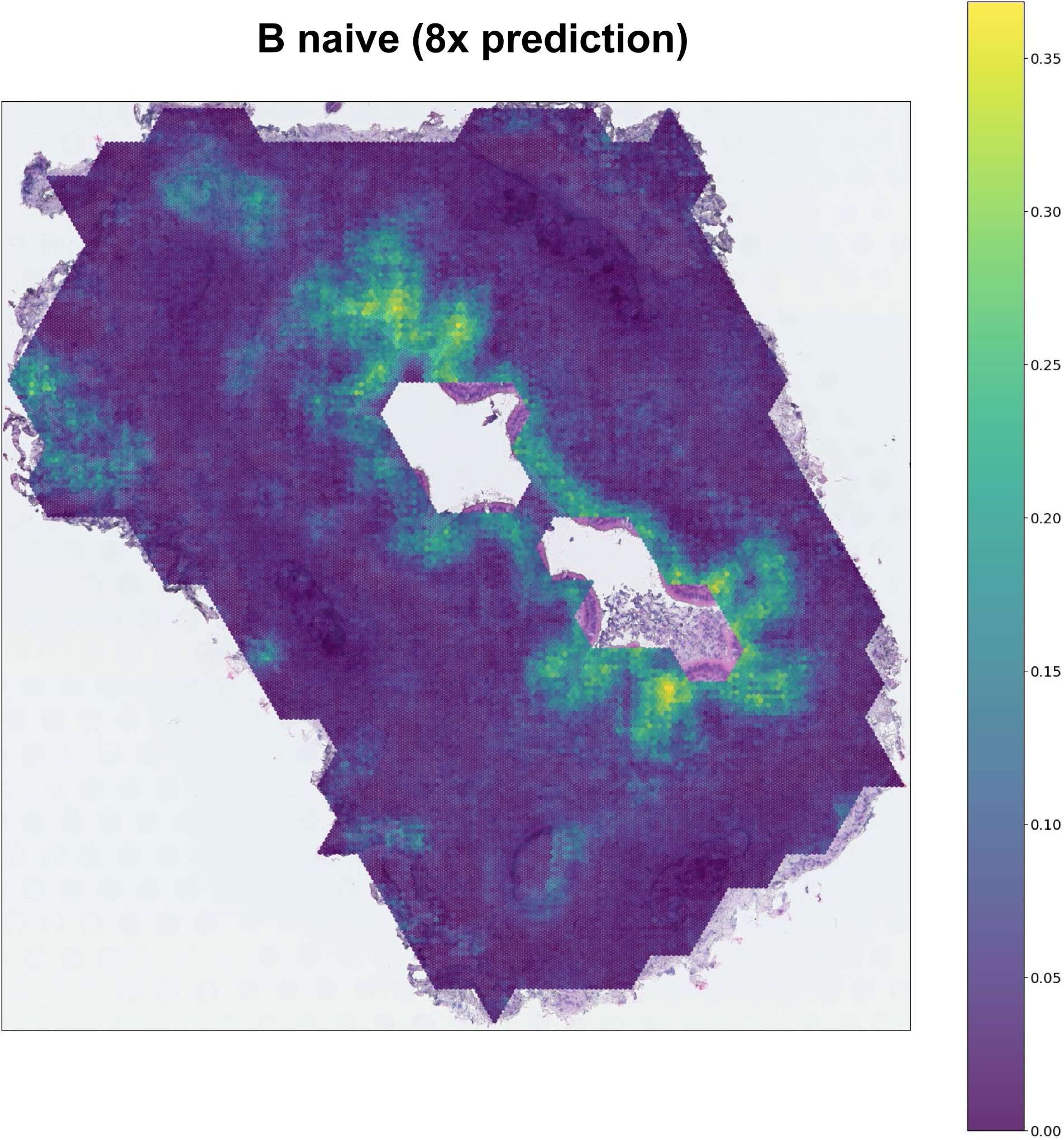
Visualization of the directly predicted 8× high-resolution results of B naive cells by Hist2Cell.

**Supplementary Fig 26:**
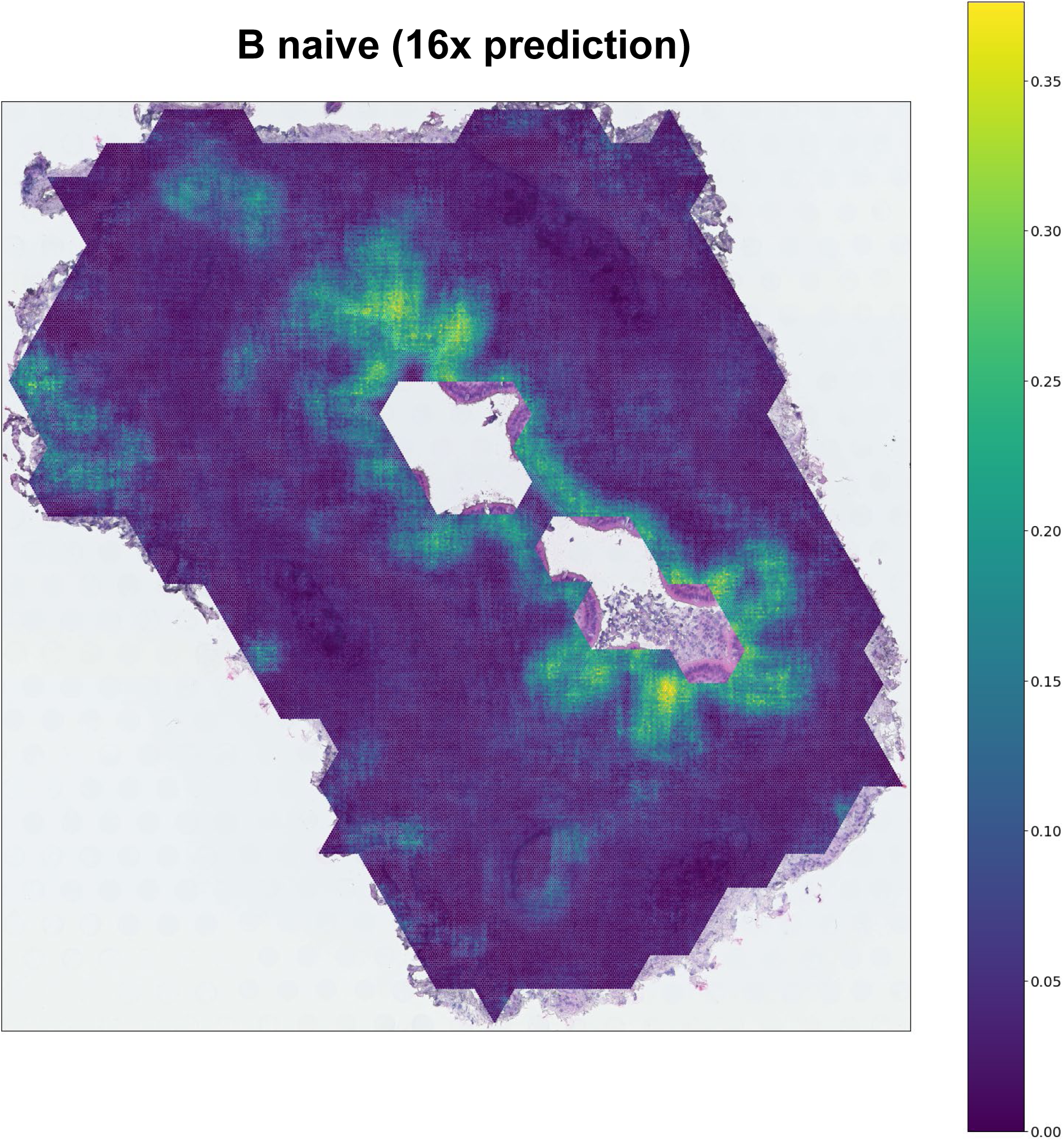
Visualization of the directly predicted 16× high-resolution results of B naive cells by Hist2Cell.

**Supplementary Fig 27:**
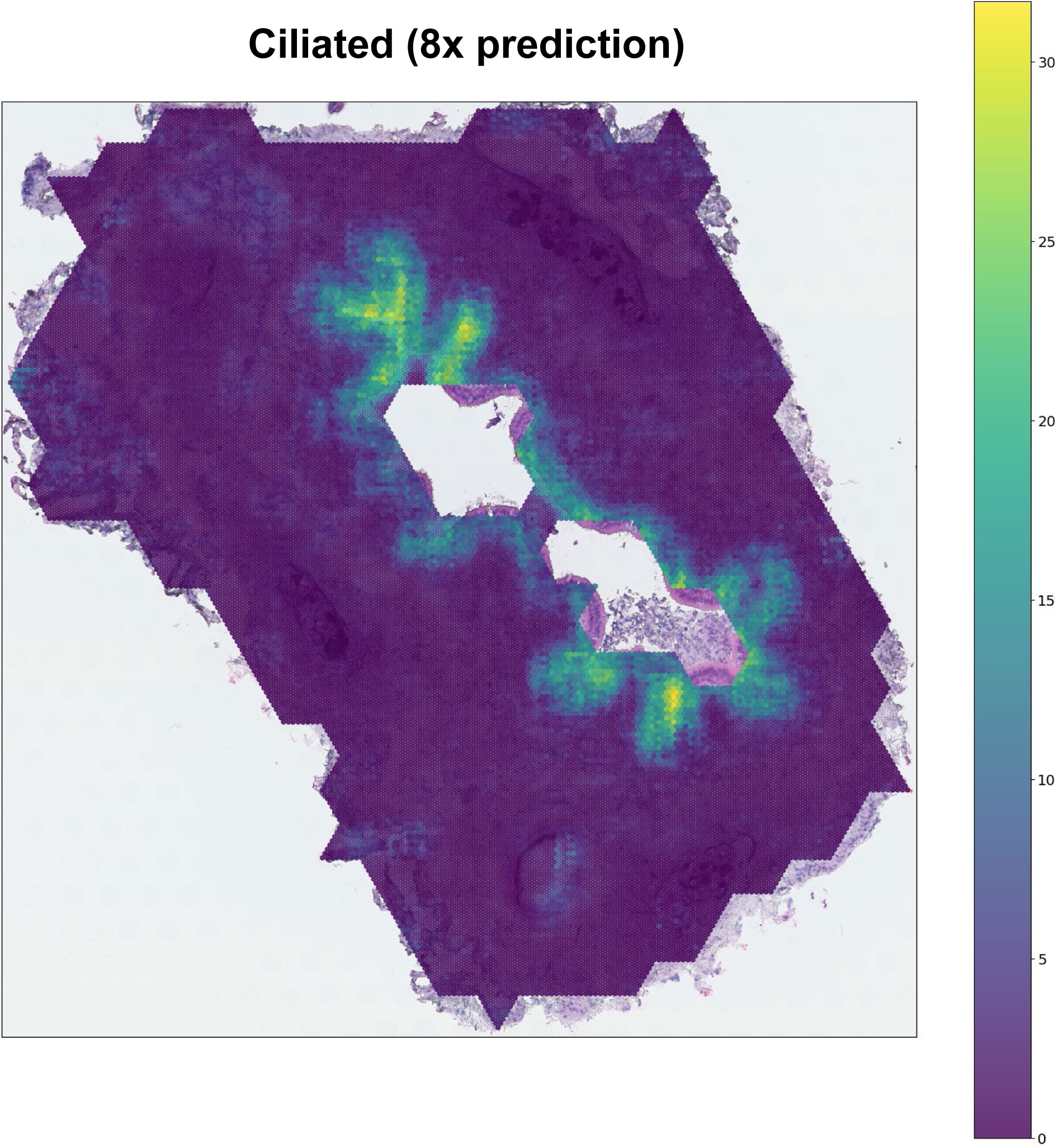
Visualization of the directly predicted 8× high-resolution results of Ciliated cells by Hist2Cell.

**Supplementary Fig 28:**
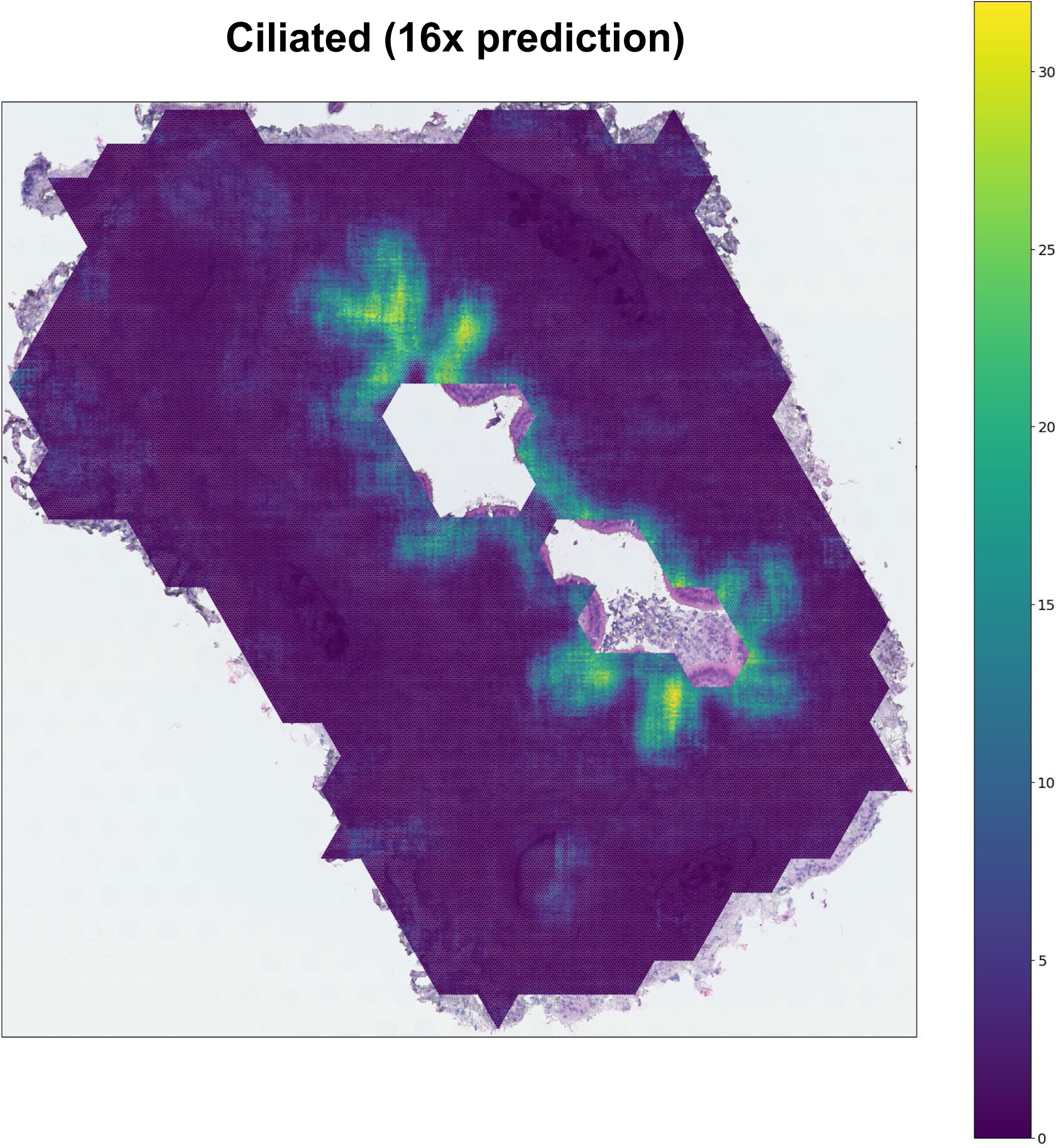
Visualization of the directly predicted 16× high-resolution results of Ciliated cells by Hist2Cell.

**Supplementary Fig 29:**
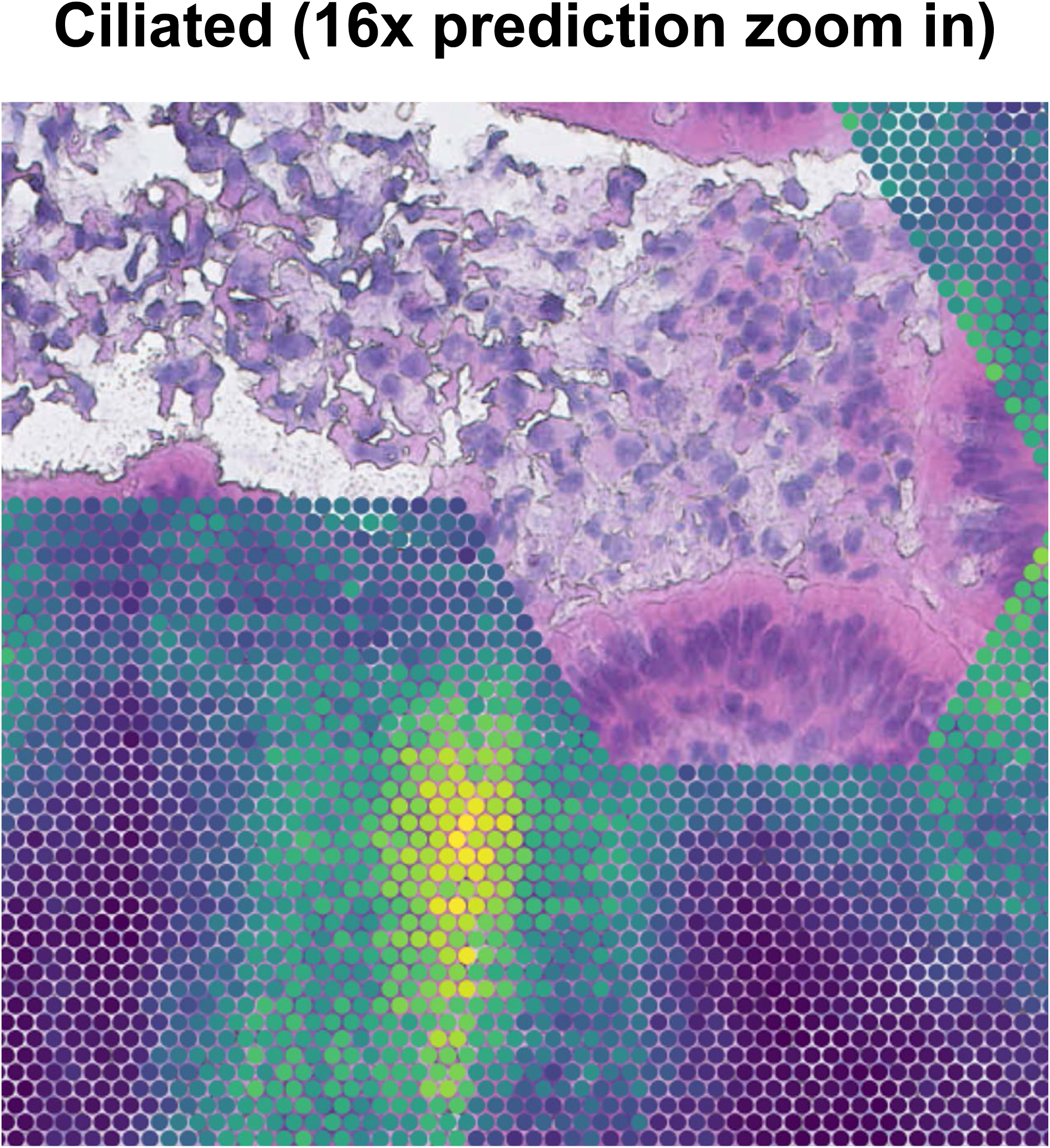
The zoom-in visualization of the directly predicted 16× high-resolution results of Ciliated cells by Hist2Cell, the sub-spot size is close and even smaller than a single cell.

**Supplementary Fig 30:**
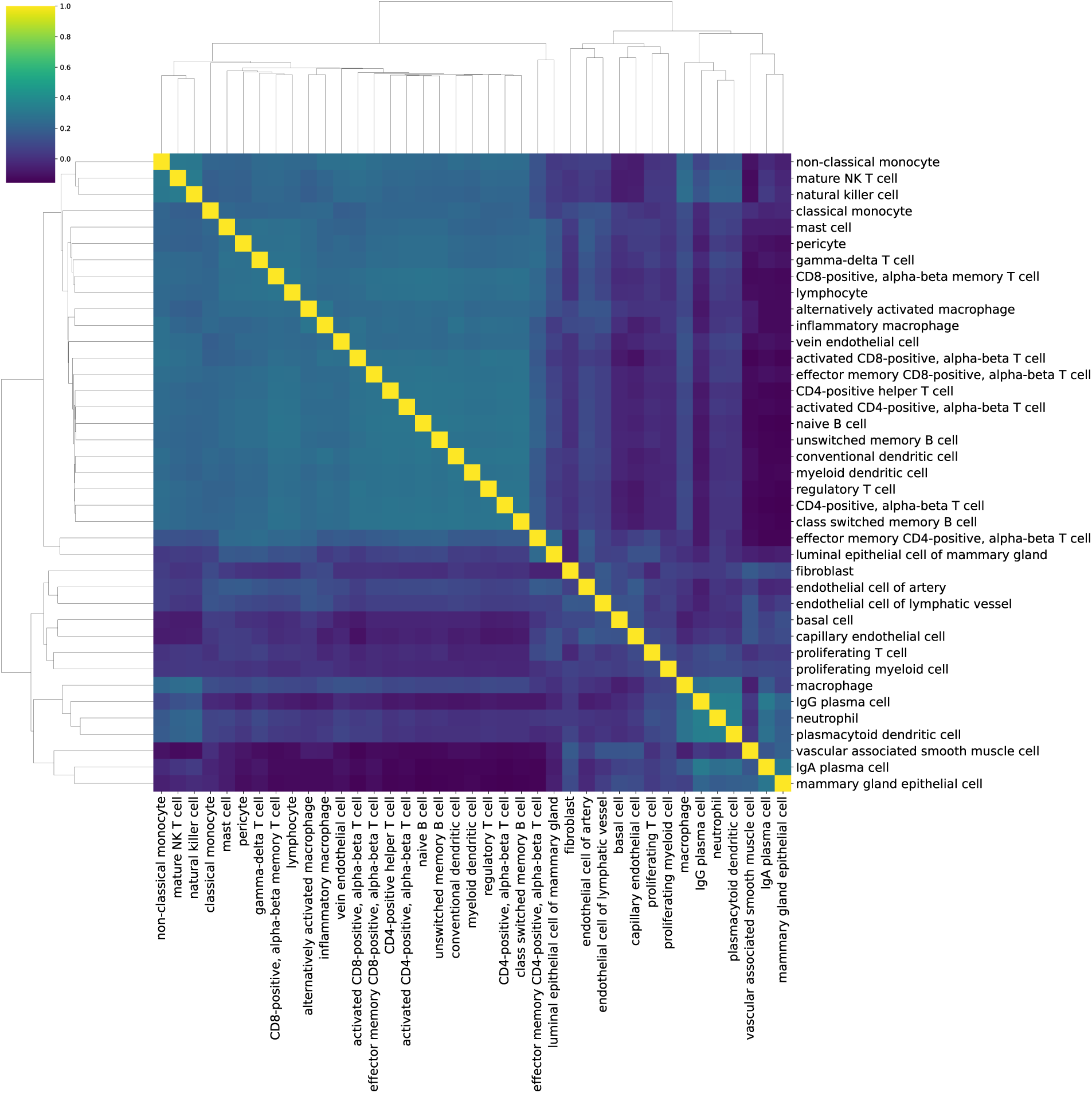
Detailed Human breast cancer Moran’s R Clustermaps calculated from Hist2Cell predictions in Fig.5.g.

**Supplementary Fig 31:**
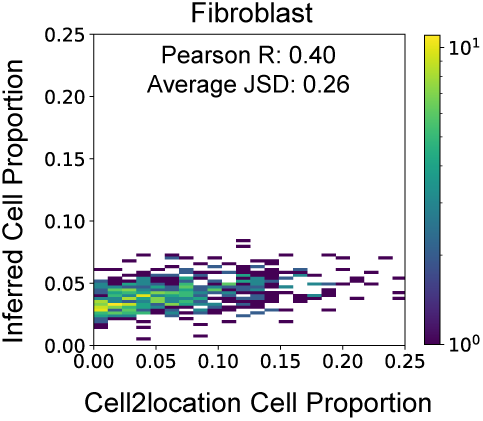
2D histogram plots showing the concordance between the ground truth cell proportion (x-axis) and the cell proportion predicted by Hist2Cell (y-axis) for fibroblast cells across all cases in the TCGA breast cancer cohort.

**Supplementary Fig 32:**
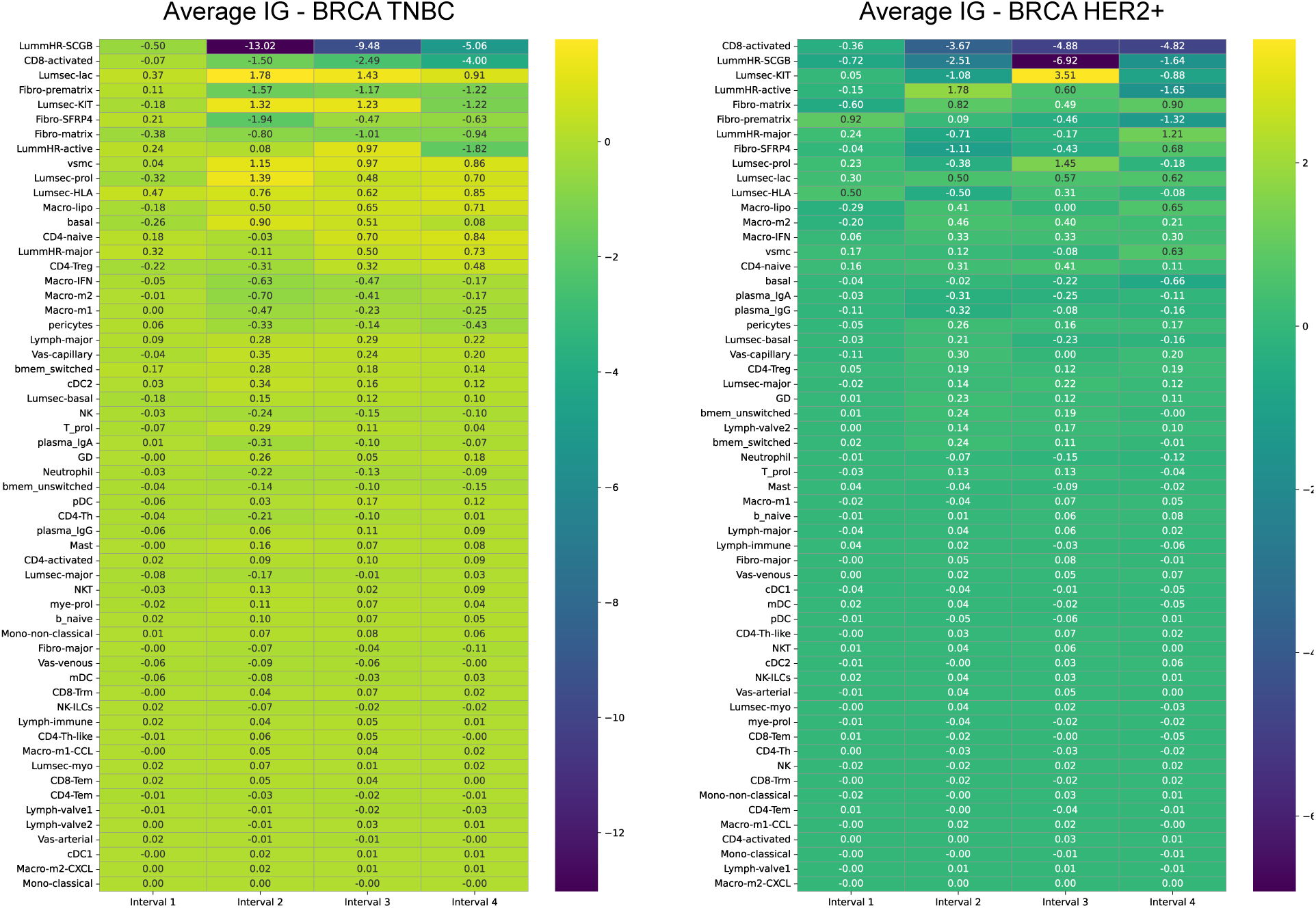
Average integrated gradients (representing the feature attribution of the Hist2Cell-prediction-based Cox regression model) for the survival analysis of BRCA TNBC and BRCA HER2+. The Y-axis is the different fine-grained transcriptional cell types and the X-axis is the four survival intervals ranging from shortest to longest survival time among the patients. The fine-grained transcriptional cell types are ranked according to their importance (mean value) among four intervals. Interesting observations could be found: (1) in the ten 10 major cell types identified by the original study of the scRNA-seq reference [45], luminal hormone-responsive (LumHR-), luminal secretory (LumSec-), fibroblasts (fibro-) and immune (CD8-activated) cells play important roles (on the top of the ranked heatmap) in predicting patients’ mortality while other cell types, like the vascular (vas-venous) and the rare cells, less related (on the lower part of the ranked heatmap) to the patient’s survival status, this aligns with the existing studies analyzing breast cancer survival [46–51]; (2) the effect of a certain cell type might vary between short-time and long-time survival analysis, for instance, we note that CD8-activated cells will have a stronger effect for long-time survival (4 times bigger average integrated gradients for later survival intervals) for HER2+ cancers, such observation provides potential biological insights for future cancer research and probably motivates customized BRCA-HER2+ treatment plan to promote the proliferation of certain cell types.

**Supplementary Fig 33:**
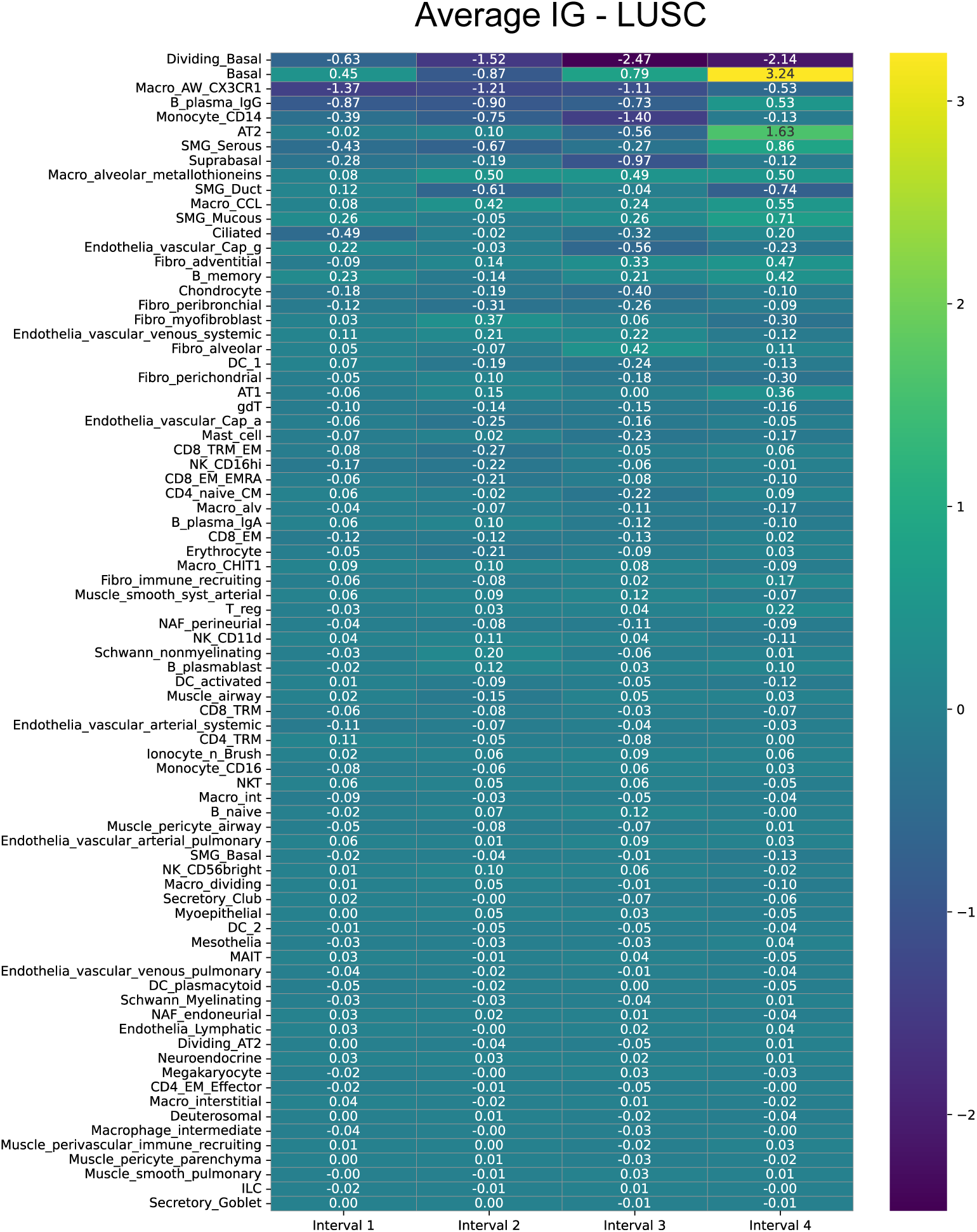
Average integrated gradients (representing the feature attribution of the modelname-prediction-based Cox regression model) for the survival analysis of LUSC. The Y-axis is the different fine-grained transcriptional cell types and the X-axis is the four survival intervals ranging from shortest to longest survival time among the patients. The fine-grained transcriptional cell types are ranked according to their importance (mean value) among four intervals.

## Footnotes

1 https://5locationslung.cellgeni.sanger.ac.uk/

2 https://cellxgene.cziscience.com/collections/4195ab4c-20bd-4cd3-8b3d-65601277e731

## References

[1] Armingol, E., Officer, A., Harismendy, O., Lewis, N.E.: Deciphering cell–cell interactions and communication from gene expression. Nature Reviews Genetics 22(2), 71–88 (2021)

[2] Bloemendal, S., Kück, U.: Cell-to-cell communication in plants, animals, and fungi: a comparative review. Naturwissenschaften 100, 3–19 (2013)

[3] Graham, S., Vu, Q.D., Raza, S.E.A., Azam, A., Tsang, Y.W., Kwak, J.T., Rajpoot, N.: Hover-net: Simultaneous segmentation and classification of nuclei in multi-tissue histology images. Medical image analysis 58, 101563 (2019)

[4] Dawood, M., Branson, K., Rajpoot, N.M., Minhas, F.: Albrt: Cellular composition prediction in routine histology images. In: Proceedings of the IEEE/CVF International Conference on Computer Vision, pp. 664–673 (2021)

[5] Song, T.-H., Sanchez, V., Daly, H.E., Rajpoot, N.M.: Simultaneous cell detection and classification in bone marrow histology images. IEEE journal of biomedical and health informatics 23(4), 1469–1476 (2018)

[6] Madissoon, E., Oliver, A.J., Kleshchevnikov, V., Wilbrey-Clark, A., Polanski, K., Richoz, N., Ribeiro Orsi, A., Mamanova, L., Bolt, L., Elmentaite, R., et al.: A spatially resolved atlas of the human lung characterizes a gland-associated immune niche. Nature Genetics 55(1), 66–77 (2023)

[7] Stickels, R.R., Murray, E., Kumar, P., Li, J., Marshall, J.L., Di Bella, D.J., Arlotta, P., Macosko, E.Z., Chen, F.: Highly sensitive spatial transcriptomics at near-cellular resolution with Slide-seqV2. Nature biotechnology 39(3), 313–319 (2021)

[8] Chen, A., Liao, S., Cheng, M., Ma, K., Wu, L., Lai, Y., Qiu, X., Yang, J., Xu, J., Hao, S., et al.: Spatiotemporal transcriptomic atlas of mouse organogenesis using DNA nanoball-patterned arrays. Cell 185(10), 1777–1792 (2022)

[9] Stuart, T., Butler, A., Hoffman, P., Hafemeister, C., Papalexi, E., Mauck, W.M., Hao, Y., Stoeckius, M., Smibert, P., Satija, R.: Comprehensive integration of single-cell data. Cell 177(7), 1888–1902 (2019)

[10] Andersson, A., Bergenstråhle, J., Asp, M., Bergenstråhle, L., Jurek, A., Fernández Navarro, J., Lunde-berg, J.: Single-cell and spatial transcriptomics enables probabilistic inference of cell type topography. Communications biology 3(1), 565 (2020)

[11] Cable, D.M., Murray, E., Zou, L.S., Goeva, A., Macosko, E.Z., Chen, F., Irizarry, R.A.: Robust decomposition of cell type mixtures in spatial transcriptomics. Nature biotechnology 40(4), 517–526 (2022)

[12] Kleshchevnikov, V., Shmatko, A., Dann, E., Aivazidis, A., King, H.W., Li, T., Elmentaite, R., Lomakin, A., Kedlian, V., Gayoso, A., et al.: Cell2location maps fine-grained cell types in spatial transcriptomics. Nature biotechnology 40(5), 661–671 (2022)

[13] Elosua-Bayes, M., Nieto, P., Mereu, E., Gut, I., Heyn, H.: Spotlight: seeded nmf regression to deconvolute spatial transcriptomics spots with single-cell transcriptomes. Nucleic acids research 49(9), 50–50 (2021)

[14] Biancalani, T., Scalia, G., Buffoni, L., Avasthi, R., Lu, Z., Sanger, A., Tokcan, N., Vanderburg, C.R., Segerstolpe, Å., Zhang, M., et al.: Deep learning and alignment of spatially resolved single-cell transcriptomes with Tangram. Nature methods 18(11), 1352–1362 (2021)

[15] Qiao, C., Huang, Y.: Reliable imputation of spatial transcriptome with uncertainty estimation and spatial regularization. bioRxiv, 2023–01 (2023)

[16] Bergenstråhle, L., He, B., Bergenstråhle, J., Abalo, X., Mirzazadeh, R., Thrane, K., Ji, A.L., Andersson, A., Larsson, L., Stakenborg, N., et al.: Super-resolved spatial transcriptomics by deep data fusion. Nature biotechnology 40(4), 476–479 (2022)

[17] Hu, J., Coleman, K., Zhang, D., Lee, E.B., Kadara, H., Wang, L., Li, M.: Deciphering tumor ecosystems at super resolution from spatial transcriptomics with TESLA. Cell systems 14(5), 404–417 (2023)

[18] Zhang, D., Schroeder, A., Yan, H., Yang, H., Hu, J., Lee, M.Y., Cho, K.S., Susztak, K., Xu, G.X., Feldman, M.D., et al.: Inferring super-resolution tissue architecture by integrating spatial transcriptomics with histology. Nature Biotechnology, 1–6 (2024)

[19] Wan, X., Xiao, J., Tam, S.S.T., Cai, M., Sugimura, R., Wang, Y., Wan, X., Lin, Z., Wu, A.R., Yang, C.: Integrating spatial and single-cell transcriptomics data using deep generative models with spatialscope. Nature Communications 14(1), 7848 (2023)

[20] Pang, M., Su, K., Li, M.: Leveraging information in spatial transcriptomics to predict super-resolution gene expression from histology images in tumors. bioRxiv, 2021–11 (2021)

[21] He, B., Bergenstråhle, L., Stenbeck, L., Abid, A., Andersson, A., Borg, Å., Maaskola, J., Lundeberg, J., Zou, J.: Integrating spatial gene expression and breast tumour morphology via deep learning. Nature biomedical engineering 4(8), 827–834 (2020)

[22] Fu, Y., Jung, A.W., Torne, R.V., Gonzalez, S., Vöhringer, H., Shmatko, A., Yates, L.R., Jimenez-Linan, M., Moore, L., Gerstung, M.: Pan-cancer computational histopathology reveals mutations, tumor composition and prognosis. Nature cancer 1(8), 800–810 (2020)

[23] Zeng, Y., Wei, Z., Yu, W., Yin, R., Yuan, Y., Li, B., Tang, Z., Lu, Y., Yang, Y.: Spatial transcriptomics prediction from histology jointly through transformer and graph neural networks. Briefings in Bioinformatics 23(5), 297 (2022)

[24] Jia, Y., Liu, J., Chen, L., Zhao, T., Wang, Y.: Thitogene: a deep learning method for predicting spatial transcriptomics from histological images. Briefings in Bioinformatics 25(1), 464 (2024)

[25] Xie, R., Pang, K., Bader, G.D., Wang, B.: Spatially Resolved Gene Expression Prediction from H&E Histology Images via Bi-modal Contrastive Learning. arXiv preprint arXiv:2306.01859 (2023)

[26] Comiter, C., Vaishnav, E.D., Ciapmricotti, M., Li, B., Yang, Y., Rodig, S.J., Turner, M., Pfaff, K.L., Jané-Valbuena, J., Slyper, M., et al.: Inference of single cell profiles from histology stains with the single-cell omics from histology analysis framework (schaf). BioRxiv, 2023–03 (2023)

[27] Andersson, A., Larsson, L., Stenbeck, L., Salmén, F., Ehinger, A., Wu, S.Z., Al-Eryani, G., Roden, D., Swarbrick, A., Borg, Å., et al.: Spatial deconvolution of her2-positive breast cancer delineates tumor-associated cell type interactions. Nature communications 12(1), 6012 (2021)

[28] Li, Z., Wang, T., Liu, P., Huang, Y.: Spatialdm: Rapid identification of spatially co-expressed ligand-receptor reveals cell-cell communication patterns. bioRxiv, 2022–08 (2022)

[29] Monjo, T., Koido, M., Nagasawa, S., Suzuki, Y., Kamatani, Y.: Efficient prediction of a spatial transcriptomics profile better characterizes breast cancer tissue sections without costly experimentation. Scientific Reports 12(1), 4133 (2022)

[30] Buggert, M., Price, D.A., Mackay, L.K., Betts, M.R.: Human circulating and tissue-resident memory cd8+ t cells. Nature Immunology, 1–11 (2023)

[31] Davis, J.D., Wypych, T.P.: Cellular and functional heterogeneity of the airway epithelium. Mucosal immunology 14(5), 978–990 (2021)

[32] Shaban, M., Bai, Y., Qiu, H., Mao, S., Yeung, J., Yeo, Y.Y., Shanmugam, V., Chen, H., Zhu, B., Nolan, G.P., et al.: Maps: Pathologist-level cell type annotation from tissue images through machine learning. bioRxiv (2023)

[33] Hu, T., Allam, M., Cai, S., Henderson, W., Yueh, B., Garipcan, A., Ievlev, A.V., Afkarian, M., Beyaz, S., Coskun, A.F.: Single-cell spatial metabolomics with cell-type specific protein profiling for tissue systems biology. Nature Communications 14(1), 8260 (2023)

[34] Zahir, N., Sun, R., Gallahan, D., Gatenby, R.A., Curtis, C.: Characterizing the ecological and evolutionary dynamics of cancer. Nature genetics 52(8), 759–767 (2020)

[35] Bartoschek, M., Oskolkov, N., Bocci, M., Lövrot, J., Larsson, C., Sommarin, M., Madsen, C.D., Lindgren, D., Pekar, G., Karlsson, G., et al.: Spatially and functionally distinct subclasses of breast cancer-associated fibroblasts revealed by single cell rna sequencing. Nature communications 9(1), 5150 (2018)

[36] Hein, S.M., Haricharan, S., Johnston, A.N., Toneff, M.J., Reddy, J.P., Dong, J., Bu, W., Li, Y.: Luminal epithelial cells within the mammary gland can produce basal cells upon oncogenic stress. Oncogene 35(11), 1461–1467 (2016)

[37] Seow, D.Y.B., Yeong, J.P.S., Lim, J.X., Chia, N., Lim, J.C.T., Ong, C.C.H., Tan, P.H., Iqbal, J.: Tertiary lymphoid structures and associated plasma cells play an important role in the biology of triple-negative breast cancers. Breast Cancer Research and Treatment 180, 369–377 (2020)

[38] DeNardo, D.G., Andreu, P., Coussens, L.M.: Interactions between lymphocytes and myeloid cells regulate pro-versus anti-tumor immunity. Cancer and Metastasis Reviews 29, 309–316 (2010)

39. Chen, R.J., Lu, M.Y., Weng, W.-H., Chen, T.Y., Williamson, D.F., Manz, T., Shady, M., Mahmood, F.: Multimodal co-attention transformer for survival prediction in gigapixel whole slide images. In: Proceedings of the IEEE/CVF International Conference on Computer Vision, pp. 4015–4025 (2021)

[40] Jaume, G., Doucet, P., Song, A.H., Lu, M.Y., Almagro-Pérez, C., Wagner, S.J., Vaidya, A.J., Chen, R.J., Williamson, D.F., Kim, A., et al.: Hest-1k: A dataset for spatial transcriptomics and histology image analysis. arXiv preprint arXiv:2406.16192 (2024)

[41] Chen, R.J., Chen, C., Li, Y., Chen, T.Y., Trister, A.D., Krishnan, R.G., Mahmood, F.: Scaling vision transformers to gigapixel images via hierarchical self-supervised learning. In: Proceedings of the IEEE/CVF Conference on Computer Vision and Pattern Recognition, pp. 16144–16155 (2022)

[42] Huuki-Myers, L.A., Spangler, A., Eagles, N.J., Montgomery, K.D., Kwon, S.H., Guo, B., Grant-Peters, M., Divecha, H.R., Tippani, M., Sriworarat, C., et al.: A data-driven single-cell and spatial transcriptomic map of the human prefrontal cortex. Science 384(6698), 1938 (2024)

[43] Chen, R.J., Ding, T., Lu, M.Y., Williamson, D.F., Jaume, G., Song, A.H., Chen, B., Zhang, A., Shao, D., Shaban, M., et al.: Towards a general-purpose foundation model for computational pathology. Nature Medicine 30(3), 850–862 (2024)

[44] Sundararajan, M., Taly, A., Yan, Q.: Axiomatic attribution for deep networks. In: International Conference on Machine Learning, pp. 3319–3328 (2017). PMLR

[45] Kumar, T., Nee, K., Wei, R., He, S., Nguyen, Q.H., Bai, S., Blake, K., Gong, Y., Pein, M., Sei, E., et al.: A spatially resolved single cell genomic atlas of the adult human breast. bioRxiv, 2023–04 (2023)

[46] Ali, H., Provenzano, E., Dawson, S.-J., Blows, F., Liu, B., Shah, M., Earl, H., Poole, C., Hiller, L., Dunn, J.A., et al.: Association between cd8+ t-cell infiltration and breast cancer survival in 12 439 patients. Annals of oncology 25(8), 1536–1543 (2014)

[47] Elsas, M.J., Middelburg, J., Labrie, C., Roelands, J., Schaap, G., Sluijter, M., Tonea, R., Ovcinnikovs, V., Lloyd, K., Schuurman, J., et al.: Immunotherapy-activated t cells recruit and skew late-stage activated m1-like macrophages that are critical for therapeutic efficacy. Cancer Cell 42(6), 1032–1050 (2024)

[48] Virassamy, B., Caramia, F., Savas, P., Sant, S., Wang, J., Christo, S.N., Byrne, A., Clarke, K., Brown, E., Teo, Z.L., et al.: Intratumoral cd8+ t cells with a tissue-resident memory phenotype mediate local immunity and immune checkpoint responses in breast cancer. Cancer Cell 41(3), 585–601 (2023)

[49] Buchsbaum, R.J., Oh, S.Y.: Breast cancer-associated fibroblasts: where we are and where we need to go. Cancers 8(2), 19 (2016)

[50] Yang, D., Liu, J., Qian, H., Zhuang, Q.: Cancer-associated fibroblasts: from basic science to anticancer therapy. Experimental & Molecular Medicine 55(7), 1322–1332 (2023)

[51] Guo, Z., Zhang, H., Fu, Y., Kuang, J., Zhao, B., Zhang, L., Lin, J., Lin, S., Wu, D., Xie, G.: Cancer-associated fibroblasts induce growth and radioresistance of breast cancer cells through paracrine il-6. Cell Death Discovery 9(1), 6 (2023)

[52] Wang, H., Li, Z., Feng, L., Zhang, W.: Vim: Out-of-distribution with virtual-logit matching. In: Proceedings of the IEEE/CVF Conference on Computer Vision and Pattern Recognition, pp. 4921–4930 (2022)

[53] Hendrycks, D., Basart, S., Mazeika, M., Zou, A., Kwon, J., Mostajabi, M., Steinhardt, J., Song, D.: Scaling out-of-distribution detection for real-world settings. arXiv preprint arXiv:1911.11132 (2019)

[54] Ståhl, P.L., Salmén, F., Vickovic, S., Lundmark, A., Navarro, J.F., Magnusson, J., Giacomello, S., Asp, M., Westholm, J.O., Huss, M., et al.: Visualization and analysis of gene expression in tissue sections by spatial transcriptomics. Science 353(6294), 78–82 (2016)

[55] Li, B., Eliceiri, K.W.: Dual-stream maximum self-attention multi-instance learning. arXiv preprint arXiv:2006.05538 (2020)

[56] Vahadane, A., Peng, T., Albarqouni, S., Baust, M., Steiger, K., Schlitter, A.M., Sethi, A., Esposito, I., Navab, N.: Structure-preserved color normalization for histological images. In: 2015 IEEE 12th International Symposium on Biomedical Imaging (ISBI), pp. 1012–1015 (2015). IEEE

[57] Yang, Y., Hossain, M.Z., Stone, E.A., Rahman, S.: Exemplar guided deep neural network for spatial transcriptomics analysis of gene expression prediction. In: Proceedings of the IEEE/CVF Winter Conference on Applications of Computer Vision, pp. 5039–5048 (2023)

[58] Yang, Y., Hossain, M.Z., Stone, E., Rahman, S.: Spatial transcriptomics analysis of gene expression prediction using exemplar guided graph neural network. Pattern Recognition 145, 109966 (2024)

[59] He, K., Zhang, X., Ren, S., Sun, J.: Deep residual learning for image recognition. In: Proceedings of the IEEE Conference on Computer Vision and Pattern Recognition, pp. 770–778 (2016)

[60] Russakovsky, O., Deng, J., Su, H., Krause, J., Satheesh, S., Ma, S., Huang, Z., Karpathy, A., Khosla, A., Bernstein, M., et al.: Imagenet large scale visual recognition challenge. International journal of computer vision 115, 211–252 (2015)

[61] Wu, Z., Jain, P., Wright, M., Mirhoseini, A., Gonzalez, J.E., Stoica, I.: Representing long-range context for graph neural networks with global attention. Advances in Neural Information Processing Systems 34, 13266–13279 (2021)

[62] Rong, Y., Bian, Y., Xu, T., Xie, W., Wei, Y., Huang, W., Huang, J.: Self-supervised graph transformer on large-scale molecular data. Advances in Neural Information Processing Systems 33, 12559–12571 (2020)

[63] Brody, S., Alon, U., Yahav, E.: How attentive are graph attention networks? arXiv preprint arXiv:2105.14491 (2021)

[64] Vaswani, A., Shazeer, N., Parmar, N., Uszkoreit, J., Jones, L., Gomez, A.N., Kaiser, Ł., Polosukhin, I.: Attention is all you need. Advances in neural information processing systems 30 (2017)

[65] Ba, J.L., Kiros, J.R., Hinton, G.E.: Layer normalization. arXiv preprint arXiv:1607.06450 (2016)

[66] Wartenberg, D.: Multivariate spatial correlation: A method for exploratory geographical analysis. Geographical Analysis 17, 263–283 (2010) 10.1111/j.1538-4632.1985.tb00849.x

[67] Rai, S., Mishra, P., Ghoshal, U.C.: Survival analysis: A primer for the clinician scientists. Indian Journal of Gastroenterology 40(5), 541–549 (2021)

[68] Zadeh, S.G., Schmid, M.: Bias in cross-entropy-based training of deep survival networks. IEEE transactions on pattern analysis and machine intelligence 43(9), 3126–3137 (2020)

[69] Schmauch, B., Romagnoni, A., Pronier, E., Saillard, C., Maillé, P., Calderaro, J., Kamoun, A., Sefta, M., Toldo, S., Zaslavskiy, M., et al.: A deep learning model to predict rna-seq expression of tumours from whole slide images. Nature communications 11(1), 3877 (2020)

